# A hydrogenase-like activity associated with mitochondrial Complex I under hypoxia in vascular plants

**DOI:** 10.64898/2026.06.02.729431

**Authors:** Xin Zhang, Zhao Zhang, Xujuan Zhang, Mengyu Liu, Yanqi Li, Pengxiang Zhao, Fei Xie, Xuemei Ma

## Abstract

Molecular hydrogen (H₂) evolution by higher plants under oxygen limitation was reported more than six decades ago, yet its origin, biochemical basis and physiological significance remain unresolved. Here, we localize this activity to a mitochondrial, Complex I-associated electron-transfer system: fractions from *etiolated Vigna radiata* hypocotyls accumulated H₂ under hypoxia at mildly acidic pH. H₂ accumulation was nearly abolished by rotenone and sensitive to perturbations of Complex II, quinone-pool turnover, terminal oxidases and protonmotive force. NADH-generating substrates and succinate-fumarate stimulated H₂ accumulation, while metabolic profiling linked H₂ production to succinate and NADH accumulation. Cumulative H₂ production exceeded the measured NADH pool by approximately two orders of magnitude, requiring sustained NADH regeneration. Our data support a model in which mature Complex I and CI*, a Complex I assembly or remodeling state, have distinct roles. TCA-cycle dehydrogenation and potentially succinate-driven reverse electron transfer (RET) through mature Complex I replenish the matrix NADH pool, whereas CI* may consume NADH through an H₂-evolving branch, regenerating NAD⁺. This model identifies a conditional redox function for CI* and an electron outlet coupled to continuous NADH cycling. Diversion of reducing equivalents into H₂ could attenuate the corresponding ROS signature, helping explain the elusive physiological evidence for plant RET. Structural and computational analyses support flavin-based electron bifurcation across the FMN-Fe-S branch, although the catalytic steps remain unverified. Together, these findings suggest that Complex I assembly states may reshape electron fate under hypoxia, exposing latent hydrogenase-related chemistry linked to Complex I’s evolutionary ancestry.

## Introduction

Plants frequently encounter oxygen limitation during seed germination, root waterlogging, and within bulky organs where gas diffusion is restricted (Bailey-Serres et al. 2012; Bailey-Serres and Voesenek 2008). Under these conditions, the mitochondrial electron transport chain (ETC) is deprived of its terminal electron acceptor, leading to over-reduction of respiratory carriers, accumulation of reducing equivalents, and an increased risk of reactive oxygen species formation upon reoxygenation (Bailey-Serres et al. 2012; Suzuki et al. 2012). How plant mitochondria dissipate excess reductive pressure during hypoxia remains a central question.

Molecular hydrogen (H₂) has long been detected in plants. H₂ emission from germinating seeds was first reported by Renwick et al. (1964), yet its biochemical source has remained unresolved for six decades. Exogenous H₂ can enhance stress tolerance and antioxidant capacity in diverse species (Jin et al. 2016; Xie et al. 2012). Unlike *Chlamydomonas reinhardtii*, which contains a canonical [FeFe]-hydrogenase (Happe and Kaminski 2002), angiosperm genomes encode no hydrogenase. The only previously reported plant H₂-producing activity, light-dependent PSII photoproduction, requires illumination, anaerobiosis and an artificial electron donor, and cannot operate in non-photosynthetic seeds (Mal’tsev et al. 1988). Thus, the enzymatic system generating H₂ in higher plants under low oxygen remains unknown.

Mitochondria are compelling candidates because hypoxia creates a redox environment that could unmask latent catalytic activities. As oxygen declines, NADH and ubiquinol accumulate, the TCA cycle is rerouted, and succinate becomes prominent (Millar et al. 2011; Møller and Sweetlove 2010). These changes are accompanied by matrix acidification and altered protonmotive force, conditions that may favor atypical electron-transfer reactions. In mammalian mitochondria, succinate-driven RET through Complex I produces superoxide at the flavin site during ischaemia–reperfusion (Chouchani et al. 2014; Murphy 2009). In plants, RET has only early biochemical indications (Rustin and Lance 1991; Storey 1971) and lacks physiological support (Huang et al., 2019). Complex I (CI; NADH:ubiquinone oxidoreductase) is therefore of interest. Its NADH-oxidizing N-module shares deep evolutionary ancestry with hydrogenase-related redox systems (Efremov and Sazanov 2012; Friedrich and Scheide 2000), and its FMN plus proximal Fe–S network shows architectural parallels to the electron-bifurcating machinery of [FeFe]-hydrogenases such as HydABC (Buckel and Thauer 2018; Katsyv et al. 2023). These observations suggest CI may retain a cryptic hydrogenase-like potential.

A plant-specific Complex I assembly intermediate, CI*, accumulates stably during plant mitochondrial CI biogenesis and contains the N, Q, and proximal-pumping modules (PP), but lacks the distal-pumping module (PD) (Braun et al. 2014; Ligas et al. 2019; Maldonado et al. 2020; Meyer, Welchen, and Carrie 2019). In animals, the N-module is added last, so no assembly intermediate is NADH-competent (Guerrero-Castillo et al. 2017); in plants, the distal membrane module is added last, leaving CI catalytically competent for NADH oxidation throughout its lifetime (Ligas et al. 2019; Schimmeyer, Bock, and Meyer 2016), yet whether this abundant intermediate serves any physiological function has remained unknown. Structural analyses show CI preserves FMN and Fe–S chain but has a more open quinone-cavity architecture than mature holo-Complex I (Klusch et al. 2021; Maldonado et al. 2020). Plant mitochondria also contain AOX, which oxidizes ubiquinol independently of complexes III and IV and is strongly associated with stress and low-oxygen metabolism (Millar et al. 2011; Vanlerberghe 2013). These features converge in germinating seeds, the classic source of plant H₂ emission.

Here, we combine thermodynamic analysis, kinetic measurements, substrate and inhibitor perturbations, metabolite profiling, and cross-species comparisons to characterize hypoxia-associated H₂ production in plant mitochondria. The results support a mitochondrial, respiratory-state-dependent process and define biochemical constraints that are consistent with a Complex I-associated electron-partitioning model. They do not, however, directly establish CI* as the catalytic species or demonstrate electron bifurcation. We therefore present the CI* model as a falsifiable mechanistic hypothesis.

## Results

### Thermodynamic and kinetic modeling supports an FMN–N1a branched pathway for hypoxic H₂ production

To identify a plausible source of endogenous H₂, we asked whether mitochondrial Complex I-related redox chemistry could support proton reduction under hypoxia, focusing on the plant Complex I assembly intermediate CI* (Maldonado et al. 2020). CI* retains the N, Q and PP modules but lacks the distal PD module, leaving FMN-N5 solvent-exposed (Figure 1B; Supplementary Calculations S1–S13).

**Figure 1.**
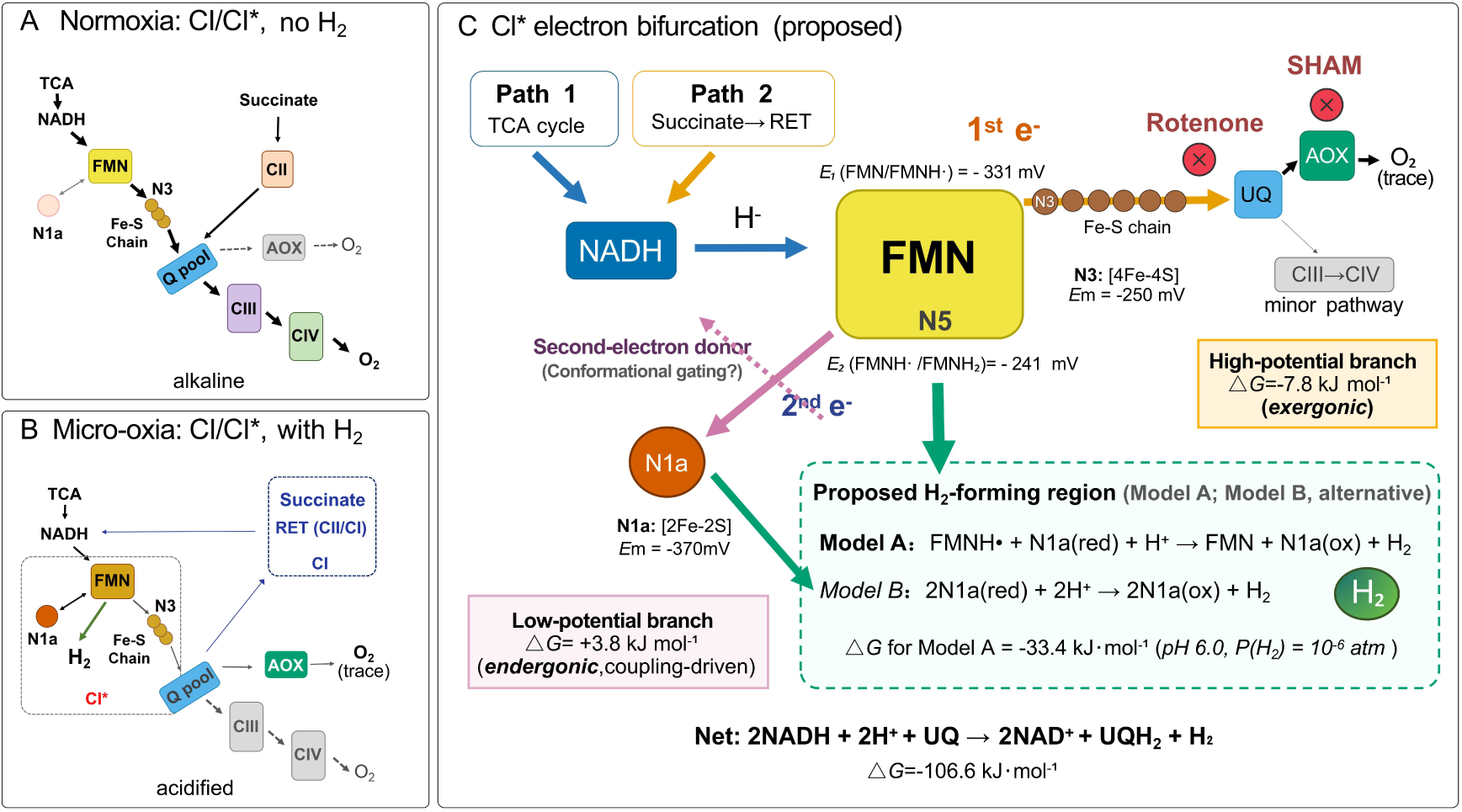
A proposed CI-associated mechanism for hypoxic H₂ production. **(A)** Normoxia. Mature Complex I (CI) and the proposed assembly intermediate CI* may coexist in the same mitochondria. CI/CI* transfers electrons from NADH through FMN, the Fe–S chain, ubiquinone, and complexes III and IV to O₂. The N1a branch is not proposed to support H₂ production under these conditions. **(B)** Micro-oxia. AOX becomes the dominant terminal acceptor (trace); succinate-driven RET through mature CI supplements NADH and may help maintain the redox conditions; CI* lacks the distal PD module, leaving the FMN-N5 region more solvent-exposed than in mature CI. CI* is the proposed H₂-producing entity in which N1a participates in the low-potential branch. **(C)** The proposed model assigns two electron-transfer branches to CI-bound FMN: NADH supplies 2e⁻ via Path 1 (TCA-cycle oxidation) or Path 2 (succinate-driven RET through mature CI). An exergonic high-potential branch toward N3 and the ubiquinone pool, and a low-potential branch toward N1a that is mildly endergonic when considered separately but favorable when coupled to H₂ formation. Model A places H–H bond formation in the FMN-N5 environment, whereas Model B places it at a transiently altered N1a iron site. Rotenone (N2 → UQ) and SHAM (AOX) abolish H₂. The calculated free-energy values are provided in Supplementary Calculations S1–S13. Structural homology with HydABC and the relevant cofactor distances are described in the main text and Supplementary Information.

Nernst-based modeling showed that direct H₂ evolution from CI-bound FMNH₂ is unfavorable under standard conditions (ΔG ≈ +14.3 kJ mol⁻¹) but becomes strongly favored by low H₂ partial pressure and a high FMNH₂/FMN ratio (ΔG ≈ −31.5 kJ mol⁻¹ at P(H₂) ≈ 10⁻⁶ atm and FMNH₂/FMN ≈ 100; Supplementary Calculation S2). Because this direct route has no net proton stoichiometry, matrix acidification does not by itself drive H₂ production.

We therefore evaluated an FMN-centered branched electron-transfer model. At pH 7.0, the one-electron potentials of CI-bound FMN are crossed, with E₁ = −390 mV (FMN/FMNH·) and E₂ = −300 mV (FMNH·/FMNH₂) (Sazanov and Hinchliffe 2006). Assuming the corresponding 1e⁻/1H⁺ pH dependence, these values shift to approximately −331 and −241 mV at pH 6.0. This relationship implies a thermodynamically unstable semiquinone, a feature commonly associated with flavin-based electron bifurcation (Buckel and Thauer 2018). The CI* N-module is structurally homologous to the bifurcating [FeFe]-hydrogenase HydABC (Furlan et al. 2022).

In this model, one electron is transferred through the high-potential branch to UQ, favorable by approximately −7.8 kJ mol⁻¹, whereas the second electron transfers to the low-potential N1a cluster, mildly unfavorable at pH 6.0. The N1a-directed transfer crosses thermodynamic neutrality near pH 6.7; coupling the two branches yields a summed free-energy change of approximately −4.0 kJ mol⁻¹, consistent with the experimental pH 6.0 optimum (Supplementary Calculation S4). The FMNH·/N1a(red) precursor is generated over two successive NADH oxidation events: the first reduces N1a while regenerating oxidized FMN, and the second traps the flavin as FMNH· (Supplementary Calculation S6). The cooperative H₂-forming reaction (FMNH· + N1a(red) + H⁺ → FMN + N1a(ox) + H₂) is calculated to be favorable by approximately −33.4 kJ mol⁻¹. Because a freely equilibrating FMNH₂ would return its reducing equivalent to the main Fe–S pathway rather than to protons, productive H₂ formation is proposed to require tight coupling of N1a electron delivery to proton transfer and H–H bond formation, without a stable FMNH₂ intermediate; a sequential back-transfer that regenerates FMNH₂ is instead excluded by the rotenone response (Supplementary Calculations S5 and S7). Unlike the direct route (FMNH₂ → H₂, Supplementary Calculation S2), this reaction consumes a matrix proton and becomes approximately 5.7 kJ mol⁻¹ more exergonic per unit decrease in pH, providing a possible explanation for the acidic pH optimum (Supplementary Calculation S5, S10). The corresponding electron-balanced net reaction (2 NADH + 2H⁺ + UQ → 2 NAD⁺ + UQH₂ + H₂) has ΔG ≈ −106.6 kJ mol⁻¹ (Supplementary Calculation S6).

Moser–Dutton analysis indicates that the modeled cofactor geometry imposes no obvious electron-tunneling barrier to the proposed pathway. Estimated rates were approximately 1.4 × 10⁹ s⁻¹ for FMN→N3 and 4.6 × 10⁵ s⁻¹ for FMN→N1a, both substantially faster than the observed H₂ flux (Supplementary Calculation S8). The branched model also provides a possible explanation for rotenone inhibition: unlike a simple FMNH₂-discharge mechanism, which might predict enhanced H₂ formation when downstream quinone reduction is blocked, the proposed pathway requires continued Fe–S/quinone electron transfer to generate the FMNH·/N1a(red) precursor. In this model, rotenone could therefore suppress H₂ production by limiting quinone-directed electron flow (Supplementary Calculation S7).

Under this framework, H₂ evolution requires three primary conditions: an active NADH/NAD⁺ cycle to sustain FMN turnover; controlled, low-flux transfer of one electron from FMNH₂ through the Fe–S chain to ubiquinone; and an acidic matrix pH, with a predicted thermodynamic crossover near pH 6.7. These conditions may in turn be established or maintained by secondary factors: micro-oxia (which accumulates NADH and acidifies the matrix, while a residual oxidized quinone fraction is preserved as the high-potential-branch acceptor), AOX-dependent quinone reoxidation (which maintains that acceptor fraction), and, where applicable, Δp-driven reverse electron transfer from succinate, which supplements the NADH pool. Two candidate H₂-forming sites remain to be distinguished experimentally (Model A, FMN-N5; Model B, N1a iron site; Supplementary Calculations S5 and S13).

Collectively, these analyses support a thermodynamically permissible and tunneling-compatible FMN–N1a branched pathway. However, obligatory branched electron-transfer chemistry, the detailed redox-state transitions and the identity of the H–H bond-forming catalyst remain to be established. The principal testable predictions are summarized in Table 1 and evaluated below.

**Table 1.** Testable predictions of the proposed CI*-associated branched electron-transfer model for H₂ production.

| Prediction | Mechanistic rationale | Experimental test |
|---|---|---|
| P1. Rotenone suppresses H<sub>2</sub> despite continued access of NADH to FMN (R2). | Blocking N2-to-UQ transfer restricts Fe–S-chain turnover and formation of the proposed H<sub>2</sub>-forming FMNH•/N1a(red) pair. | Quinone-site inhibition with H<sub>2</sub> and redox measurements; distinguish branch inhibition from reduced RET-dependent NADH supply. |
| P2. FCCP suppresses H<sub>2</sub>, whereas oligomycin has weaker or distinct effects (R1). | Δp supports NADH regeneration through RET in mature CI; ATP synthesis is not a direct requirement of the proposed H<sub>2</sub>-forming reaction. | Compare FCCP and oligomycin while monitoring NADH, Δp and matrix pH. |
| P3. AOX inhibition suppresses H<sub>2</sub> under micro-oxia (R2). | AOX reoxidizes ubiquinol, maintaining an oxidized-UQ fraction for high-potential-branch turnover. | AOX inhibition with H<sub>2</sub>, O<sub>2</sub>-consumption and quinone-redox measurements. |
| P4. Complex III inhibition is less inhibitory than AOX inhibition under micro-oxia (R2). | AOX can sustain quinol oxidation when the cytochrome pathway is blocked, although reduced proton pumping may limit RET. | Compare antimycin A and AOX inhibition under matched O<sub>2</sub> conditions. |
| P5. Sustained H<sub>2</sub> production requires matrix NADH/NAD<sup>+</sup> cycling (R1). | CI* consumes NADH and regenerates NAD<sup>+</sup>; TCA-cycle dehydrogenation and proposed RET through mature CI replenish the shared NADH pool. | Compare cumulative H<sub>2</sub> output with NADH/NAD<sup>+</sup> pools and perturb NADH regeneration. |
| P6. Complex II inhibition reduces succinate-stimulated H<sub>2</sub> production (R1). | Succinate oxidation supplies ubiquinol that may drive Δp-dependent NADH regeneration through mature CI. | Test succinate stimulation with malonate and FCCP; measure NADH formation, quinone redox state and Δp. |
| P7. Malate and succinate support H<sub>2</sub> production, whereas excess oxidized UQ suppresses it (R1, R2). | TCA-linked and RET-compatible inputs converge on matrix NADH; excess oxidized UQ is predicted to favor the quinone branch over H<sub>2</sub> formation. | Substrate supplementation and oxidized-UQ titration, with rotenone and malonate controls. |
| P8. Moderately acidic matrix conditions favor H<sub>2</sub> formation (R3). | The proposed cooperative H<sub>2</sub>-forming step consumes a matrix proton; excessive acidification may impair supporting reactions. | Measure H<sub>2</sub> across a pH series together with matrix pH; bulk-medium pH alone cannot establish the catalytic optimum. |
| P9. H<sub>2</sub> production is favored within a micro-oxic redox window (R1, R2). | O<sub>2</sub> limitation increases reducing pressure, while residual O<sub>2</sub>-dependent quinol oxidation prevents complete acceptor depletion. | Controlled O<sub>2</sub>-gradient assays with H<sub>2</sub> and redox measurements; explicitly test sustained strict anoxia. |
| P10. H<sub>2</sub>-forming activity is proposed to associate with CI* abundance and assembly state (M). | CI* is proposed to permit H<sub>2</sub>-forming chemistry, while mature CI supports the RET-dependent NADH supply route. | Native fractionation, immunodetection and activity assays; selective depletion or reconstitution to establish catalytic attribution. |

*R1, sustained matrix NADH/NAD<sup>+</sup> cycling; R2, controlled electron withdrawal through the Fe–S/UQ branch; R3, sufficiently acidic matrix conditions; M, proposed CI catalytic identity. The model assigns distinct roles to coexisting mature CI and CI\*; RET competence of CI\* is not assumed. These tests assess model compatibility but do not individually establish RET, electron bifurcation or direct H<sub>2</sub> catalysis by CI\*.*

### Etiolated mung bean seedlings accumulate H₂ under low-oxygen conditions

We first examined whether etiolated mung bean seedlings accumulated detectable H₂ in sealed vials (Figure 2A). GC–BID resolved H₂ and O₂ as distinct peaks against authentic standards (Figure 2B). GC–BID peak area increased linearly with H₂ amount over the relevant analytical ranges, permitting peak-area-based comparison across whole-seedling and excised-organ assays and absolute quantification in the same 1.9-mL vial/500-μL system used for mitochondrial measurements (Figure 2—figure supplement 1). H₂ accumulation was readily detected in mung bean seedlings but not in empty vials or sterile-water controls, with substantially stronger signals under N₂-flushed than under air-started conditions (Figure 2C–E). Bacterial contamination was not detected by 16S rRNA PCR or LB plating, consistent with earlier evidence that seedlings possess an intrinsic H₂-producing capacity independent of bacteria (Renwick et al. 1964; Torres, Ballesteros, and Fernández 1986).

**Figure 2.**
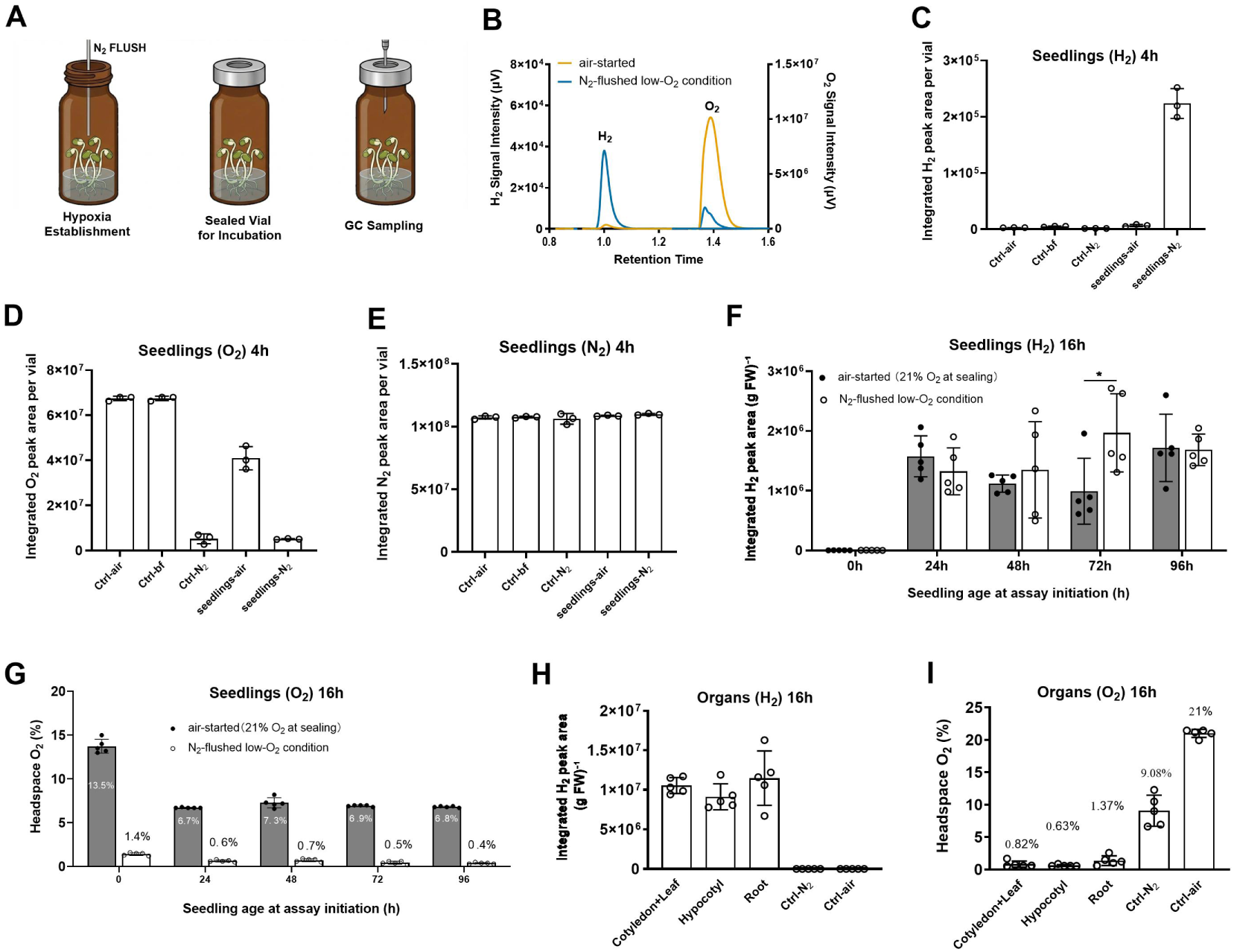
Detection, developmental dynamics, and organ distribution of H₂ accumulation in etiolated mung bean seedlings. **(A)** Schematic of the sealed-vial headspace assay. Seedlings (10 per 40-mL vial) or excised organs (4–5 g fresh weight per 22.5-mL vial) were sealed under ambient air (air-started; ≈21% O₂ at sealing) or after flushing the headspace with N₂ for 2 min; vials were incubated in darkness with single endpoint sampling. **(B)** Representative GC–BID chromatograms of seedlings in air-started or N₂-flushed vials. H₂ and O₂ were baseline-separated (retention times 1.0 ± 0.1 and 1.4 ± 0.1 min) and identified against authentic standards; separate *y*-axes reflect the large difference in detector response. Calibration is detailed in Figure 2—figure supplement 1.**(C–E)** Integrated peak areas of H₂ (C), O₂ (D), and N₂ (**E**) after 4 h in empty air-filled vials (Ctrl-air), buffer-only vials (Ctrl-bf), N₂-flushed empty vials (Ctrl-N₂), air-started vials containing seedlings (seedlings-air), and N₂-flushed vials containing seedlings (seedlings-N₂) (*n* = 3). Comparable N₂ peak areas across conditions indicate consistent sampling and injection. **(F, G)** Endpoint headspace H₂ (F) and O₂ (**G**) from seedlings of the indicated ages after 16 h in independently prepared air-started or N₂-flushed vials (*n* = 5); H₂ normalized to seedling fresh weight. Numbers in (**G**) indicate endpoint headspace O₂ concentrations. Inter-condition differences at each time point were tested by two-way ANOVA with Šídák’s multiple-comparisons correction. **(H, I)** Endpoint headspace H₂ (**H**) and O₂ (**I**) from cotyledon–leaf, hypocotyl, and root organs excised from 72-h-old seedlings and incubated for 16 h in N₂-flushed vials (*n* = 5); H₂ normalized to organ fresh weight, numbers in (**I**) indicate endpoint headspace O₂ concentrations. Ctrl-N₂ and Ctrl-air are sterile-water controls without plant organs. Seedlings (*Vigna radiata*) were germinated in darkness at 25 °C. Data are shown without background subtraction as mean ± SD; points represent independent vials. Absolute GC responses should be compared only within panels because experiments were performed independently under assay-specific conditions. Statistical significance: \**P* < 0.05; \*\**P* < 0.01; \*\*\**P* < 0.001; \*\*\*\**P* < 0.0001; ns, not significant. GC calibration and analytical details are provided in Figure 2—figure supplement 1 and Methods.

Seedlings germinated for 24-96 h accumulated H₂ during a subsequent 16-h incubation under both air-started and N₂-flushed conditions (Figure 2F). Residual headspace O₂ was 6.7-13.5% in air-started vials versus 0.4-1.4% in N₂-flushed vials (Figure 2G), consistent with the stronger H₂ signal under low-O₂ but not fully anoxic conditions and providing physiological support for prediction P9 (Table 1); likewise, earlier work reported that headspace H₂ accumulated to a peak as O₂ fell to ≈15% (Renwick et al. 1964). Because only air-started and N₂-flushed conditions were compared, these experiments do not define an O₂ optimum or test the effect of complete anoxia. Excised cotyledon–leaf, hypocotyl, and root organs from 72-h-old seedlings all accumulated H₂ under N₂-flushed conditions, with organ-dependent differences in signal intensity (Figure 2H, I). Roots showed a comparable H₂ signal despite retaining more residual O₂ (1.37%) than cotyledons plus true leaves (0.82%) or hypocotyls (0.63%) (Figure 2I), suggesting that organ-specific H₂ accumulation was not determined solely by terminal headspace O₂ levels. Together, these results demonstrate developmental and spatial variation in H₂ accumulation in mung bean seedlings and identify 72-96-h-old seedlings as suitable material for subsequent biochemical analyses.

### H₂-accumulating activity is recovered in crude mitochondrial fractions and depends strongly on assay conditions

Differential centrifugation showed that H₂-accumulating activity was enriched in sedimentable fractions of mung bean hypocotyl homogenates: activity was recovered in the 12,000 × *g* crude organelle pellet and was further enriched in the 80,000 × *g* pellet derived from the corresponding supernatant, whereas little activity remained in the final supernatant (Figure 3—figure supplement 1). These findings indicate association with particulate cellular material but, by themselves, do not establish exclusive mitochondrial localization; the crude mitochondrial fraction was therefore used for subsequent characterization.

**Figure 3.**
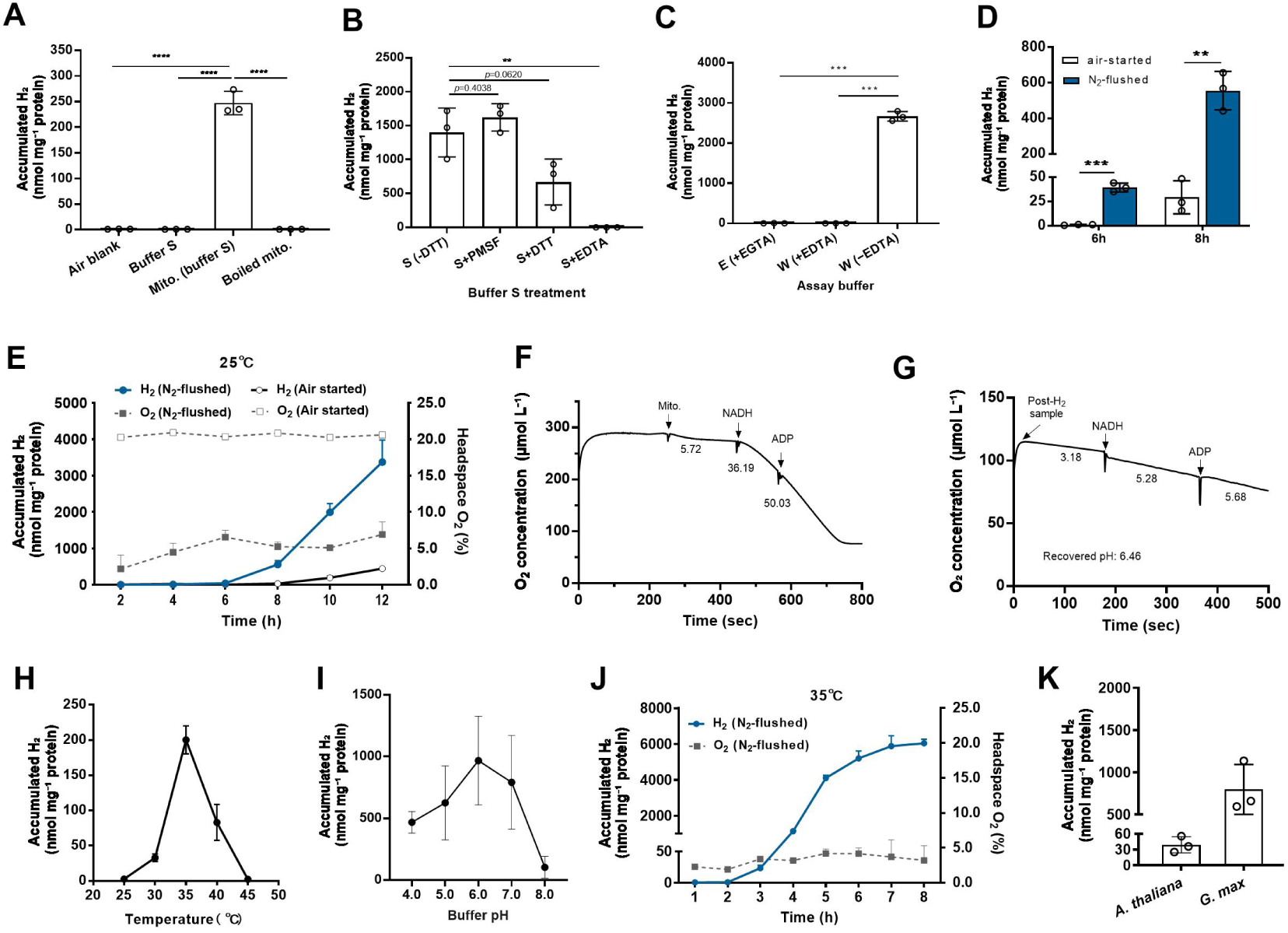
Development and characterization of the mitochondrial H₂-accumulation assay. Unless otherwise indicated, crude mitochondrial fractions were isolated from hypocotyls of 72-96-h-old etiolated *Vigna radiata* seedlings and incubated in sealed 1.9-mL GC vials containing 500 μg protein in 500 μL reaction mixture without added respiratory substrates. Vial headspaces were flushed with N₂ for 2 min unless otherwise indicated. Except in **(A)**, H₂ values were corrected against buffer-only controls and normalized to mitochondrial protein. Buffer compositions are provided in Methods. **(A–D)** Initial assay-condition screening at 25 °C. Reactions were incubated for 8 h unless otherwise indicated. **(A)** H₂ detected in an empty air-started vial, a Buffer S blank, untreated mitochondria in Buffer S, and mitochondria boiled at 100 °C for 5 min before incubation. Blank values were expressed on a protein-equivalent basis using a nominal input of 0.5 mg protein per vial. Buffer S contained 1 mM DTT. **(B)** Effects of adding PMSF, DTT, or EDTA-Na₂ (1 mM each) to Buffer S lacking these additives [S (−DTT)]. **(C)** H₂ accumulation in Buffer E containing EGTA [E (+EGTA)], Buffer W containing EDTA-Na₂ [W (+EDTA)], or EDTA-free Buffer W [W (−EDTA)]. **(D)** H₂ accumulation in EDTA-free Buffer W after 6 or 8 h under air-started or N₂-flushed conditions. Data were replotted from the corresponding time points in **(E)**. **(E)** Time courses of H₂ accumulation and headspace O₂ concentration in EDTA-free Buffer W at pH 7.2 and 25 °C. Filled and open symbols indicate N₂-flushed and air-started vials, respectively. H₂ and O₂ were quantified from the same GC injection. Headspace O₂ was estimated from the integrated GC peak area using an air-started control as the 21% O₂ reference; each time point was measured using an independent endpoint vial. **(F, G)** Oxygen-electrode measurements in EDTA-free Buffer W at pH 7.2 and 25 °C. Numbers adjacent to the traces indicate O₂-consumption rates in μmol O₂ L⁻¹ min⁻¹. **(F)** Respiratory response of freshly isolated mitochondria after sequential addition of mitochondrial protein (500 μg), NADH (1 mM), and ADP (200 μM). **(G)** Respiratory response of mitochondria recovered after the 6-h H₂ assay in **(E)**. The reaction mixture was transferred directly to the electrode chamber without washing or resuspension; the recovered bulk-medium pH was 6.46. **(H)** Effect of temperature on H₂ accumulation after 4 h in EDTA-free Buffer W. **(I)** Effect of bulk-medium pH on H₂ accumulation after 4 h at 35 °C in EDTA-free Buffer W. **(J)** Time courses of H₂ accumulation and headspace O₂ concentration at 35 °C in Buffer H, initially adjusted to pH 6.0. O₂ was quantified as in **(E)**. **(K)** H₂ accumulation by crude mitochondrial fractions from roots of 1-week-old etiolated *Glycine max* seedlings and 2-week-old *Arabidopsis thaliana* seedlings after incubation in Buffer H at 35 °C for the indicated times. *G. max* seedlings were grown in darkness, whereas *A. thaliana* seedlings were grown under a 16-h-light/8-h-dark photoperiod. Quantitative data are mean ± SD; circles represent individual assay vials, n=3. Oxygen-electrode traces in **(F)** and **(G)** are representative. One-way ANOVA followed by Dunnett’s test was used in **(A)** and **(B)**; Brown–Forsythe and Welch ANOVA followed by Dunnett’s T3 test was used in **(C)**; and two-tailed unpaired *t*-tests were used in **(D)**. Exact *P* values are shown in **(B)**. Panels **(E)** and **(J)** show descriptive time courses; **(H)**, **(I)**, and **(K)** show descriptive trends without inferential testing. Statistical significance: \**P* < 0.05; \*\**P* < 0.01; \*\*\**P* < 0.001; \*\*\*\**P* < 0.0001; ns, not significant.

Crude mitochondrial fractions accumulated substantially more H₂ than air or buffer blanks, and boiling at 100 °C for 5 min almost completely eliminated the signal (Figure 3A), indicating a heat-sensitive component. Buffer composition had a pronounced effect: DTT and especially EDTA suppressed the signal, whereas PMSF had little detectable effect (Figure 3B); fractions incubated in EDTA-free Buffer W accumulated robust H₂, whereas signals in EGTA-containing Buffer E or EDTA-containing Buffer W remained close to baseline (Figure 3C). EDTA-free conditions were therefore used for further characterization.

N₂-flushed fractions accumulated more H₂ than air-started fractions at both 6 and 8 h (Figure 3D). At 25 °C, H₂ remained low during the early incubation period and subsequently increased as headspace O₂ remained low, whereas accumulation in air-started vials was markedly weaker (Figure 3E). Fresh crude mitochondrial fractions in EDTA-free Buffer W (pH 7.2) consumed O₂ and responded to NADH and ADP (Figure 3F). Fractions recovered after a 6-h H₂ assay retained measurable respiratory responses, although their activity was lower than that of freshly isolated material (Figure 3G); the external pH had decreased from 7.2 to 6.46 over the 6-h incubation. The low-O₂ dependence of this activity parallels that in seedlings, supporting prediction P9 (Table 1). Residual O₂ remained detectable during H₂ accumulation, consistent with, but not proving, a micro-oxic redox-window requirement; controlled O₂-gradient and strictly anoxic assays will be required to establish the predicted optimum and test whether complete anoxia suppresses H₂ formation. H₂ accumulation was maximal near 35 °C and was favored by mildly acidic to neutral assay conditions. Comparable H₂ levels were observed from pH 5 to pH 7, with the highest mean signal at an initial bulk-medium pH of approximately 6; activity declined markedly at pH 4 and pH 8 (Figure 3H, I). This broad pH range is compatible with prediction P8 (Table 1), which concerns a mildly acidic matrix requirement; matrix pH was not measured directly, so the bulk-pH optimum is taken as a practical assay condition rather than a direct test of the matrix-proton dependence.

Buffer H, the EDTA-free Buffer W adjusted to pH 6.0, was consequently adopted for subsequent experiments. Under these conditions, H₂ increased sharply after an initial lag and approached a plateau by 7-8 h, while headspace O₂ remained low (Figure 3J). Crude mitochondrial fractions from soybean and *Arabidopsis* roots also accumulated H₂, although the magnitude and kinetics differed between species (Figure 3K). Collectively, these experiments define low O₂, the absence of EDTA, mildly acidic pH, and 35 °C as favorable conditions for H₂ accumulation by crude plant mitochondrial fractions.

### Inhibitor and NADH responses are consistent with the branched model

To test the pharmacological predictions P1–P4 (Table 1), which follow directly from the branched electron-transfer framework developed above (Figure 1C), we treated crude mitochondria with a panel of mitochondrial inhibitors or the uncoupler FCCP under optimized conditions (35 °C, pH 6.0, 3 h; Figure 4B). No exogenous respiratory substrates were added.

**Figure 4.**
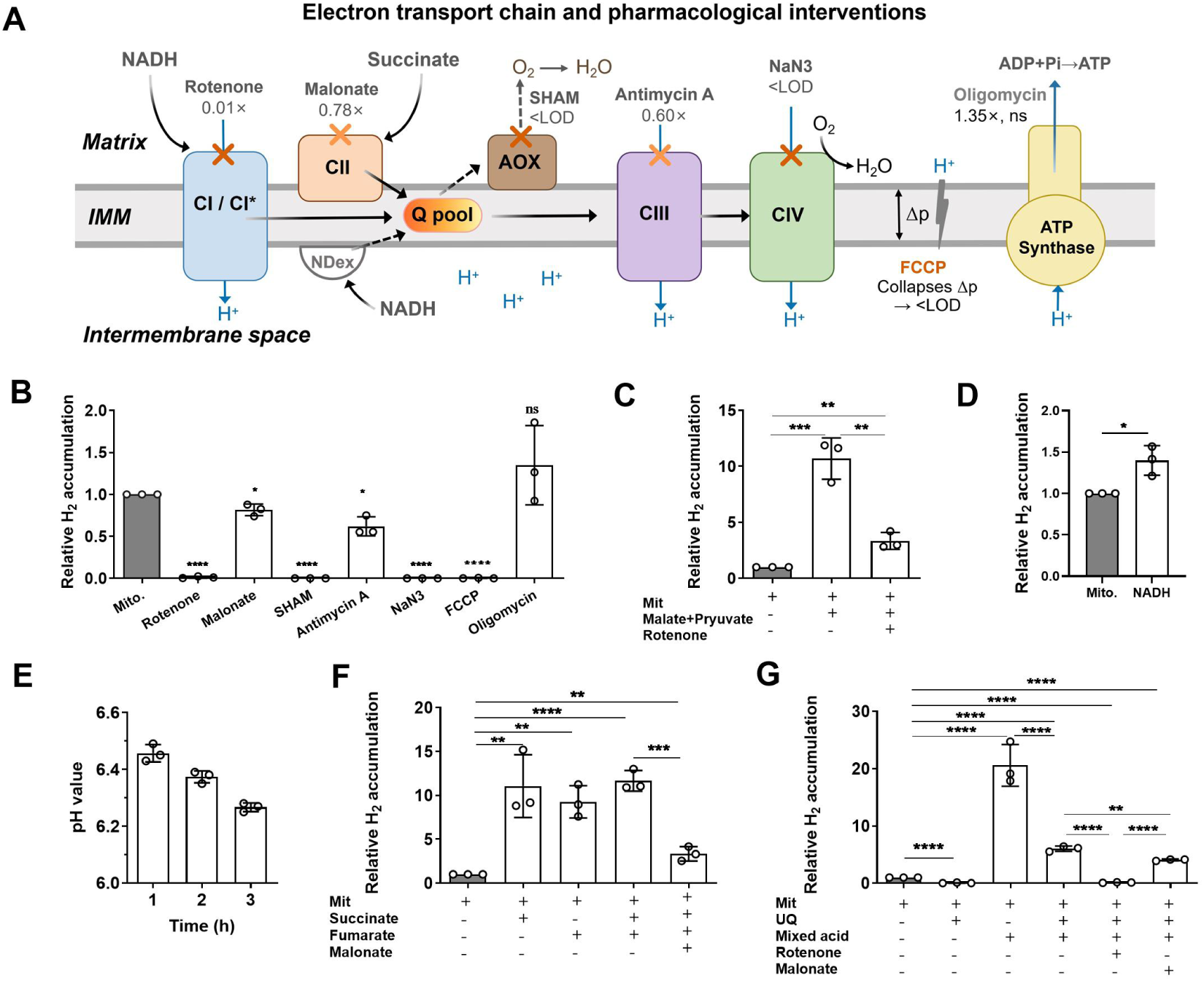
Inhibitor and substrate analyses reveal respiratory-chain dependencies of H₂ production. **(A)** Schematic representation of the mitochondrial electron-transport chain and the sites of pharmacological intervention. CI/CI* denotes coexisting mature Complex I and the proposed H₂-associated assembly intermediate described in Figure 1. Rotenone, malonate, SHAM, antimycin A, NaN₃, and oligomycin inhibit Complex I, Complex II, alternative oxidase, Complex III, Complex IV, and ATP synthase, respectively; FCCP dissipates the proton motive force (Δp). Values indicate mean H₂ accumulation relative to the untreated mitochondrial-only control in **(B)**. NDex, external NADH dehydrogenase; UQ pool, ubiquinone pool; IMM, inner mitochondrial membrane; IMS, intermembrane space; LOD, limit of detection. **(B)** Effects of mitochondrial inhibitors and FCCP on H₂ accumulation without added respiratory substrates. Crude mitochondria (300-500 μg protein) isolated from *Vigna radiata* hypocotyls were incubated under hypoxia for 3 h at 35 °C in Buffer H (pH 6.0, 500 μL), either without inhibitor (Mito.) or with rotenone (5 μM), malonate (10 mM), SHAM (2 mM), antimycin A (10 μM), NaN₃ (5 mM), FCCP (25 μM), or oligomycin (2 μM). **(C–G)** Substrate supplementation and ubiquinone competition experiments. All reactions contained crude mitochondria (500 μg) in Buffer H (pH 6.0, 500 μL), incubated under hypoxia at 35 °C for 3 h. H₂ was quantified by headspace GC–BID. **(C)** Path 1 (TCA-derived NADH): Malate (5 mM) plus pyruvate (5 mM), which generate matrix NADH via malate dehydrogenase and the pyruvate dehydrogenase complex. **(D)** Exogenous NADH (1 mM) supplied through NDex. **(E)** Change in bulk-medium pH during mitochondrial incubation. The reaction medium was initially adjusted to pH 6.5 and measured after 1, 2, and 3 h using a calibrated pH electrode (Oxytherm, Hansatech Instruments). **(F)** Effects of Complex II-linked substrates. Crude mitochondria were incubated alone or supplemented with succinate (10 mM), fumarate (5 mM), or both substrates, with or without malonate (10 mM). **(G)** Effect of exogenous oxidized ubiquinone on substrate-stimulated H₂ accumulation. Crude mitochondria were incubated with oxidized ubiquinone-10 (UQ; 60 μM), alone or in combination with a mixed-substrate solution containing malate (5 mM), pyruvate (5 mM), succinate (10 mM), and fumarate (5 mM), with or without rotenone (5 μM) or malonate (10 mM). Except for the absolute pH values in **(E)**, data were normalized to the corresponding mitochondria-only control. Bars show mean ± SD. See Supplementary Table S2 for complete raw data. Sample sizes were n=3 in **(B–G)**. Statistical significance was assessed by one-way ANOVA followed by Dunnett’s multiple-comparisons test in **(B)**, a two-sided paired test in **(D)**, and one-way ANOVA followed by Šídák’s multiple-comparisons test in **(C, E–G)**. Statistical significance: \**P* < 0.05; \*\**P* < 0.01; \*\*\**P* < 0.001; \*\*\*\**P* < 0.0001; ns, not significant.

Rotenone (5 μM), which blocks the N2-to-ubiquinone step without impeding NADH→FMN hydride transfer, reduced H₂ to 1.32 ± 1.02% of the mitochondria-only control (mean ± SD; *P* < 0.0001), consistent with **P1**: continued Fe–S/quinone-branch electron flow, rather than FMN reduction per se, is required for H₂ formation. SHAM (2 mM) and NaN₃ (5 mM) each reduced H₂ below the limit of detection (LOD, 0.1 nmol vial⁻¹), whereas antimycin A (10 μM) caused only partial inhibition (38.1 ± 11.3%; *P* < 0.01), consistent with **P3–P4**: AOX is the principal terminal acceptor sustaining ubiquinone re-oxidation under micro-oxia. By blocking electron transfer to oxygen, azide additionally removes the cytochrome branch’s contribution to proton pumping; this Δp-side consequence of the same lesion is addressed together with the FCCP result below and developed in the Discussion. FCCP (25 μM) likewise reduced H₂ below the LOD, whereas oligomycin (2 μM) did not significantly alter H₂ accumulation (134.9 ± 47.2% of the mitochondria-only control; P = 0.33). The FCCP response is consistent with a permissive role for Δp (P2), whereas the oligomycin result leaves the contribution of Complex V-dependent ATP synthesis unresolved.

Exogenous NADH significantly increased H₂ accumulation relative to the mitochondria-only control (Figure 4D; ∼1.4-fold; two-sided paired test) and stimulated oxygen consumption 6.3-fold in freshly isolated preparations (5.72 → 36.19 μmol O₂ L⁻¹ min⁻¹; Figure 3F), consistent with the external NADH dehydrogenase (NDex) actively feeding electrons into the ubiquinone pool without direct access to the CI*-bound FMN. The modest ∼1.4-fold enhancement, nearly an order of magnitude smaller than the 10.7-fold increase elicited by matrix NADH-generating substrates (Figure 4C) despite a 6.3-fold stimulation of oxygen consumption by the same NADH addition, indicates that H₂ formation is driven primarily by direct electron supply to the CI* FMN from the matrix NADH pool (**R1**), with NDex-mediated ubiquinone-pool reduction supporting H₂ only indirectly. The bulk-medium pH declined from 6.46 ± 0.03 at 1 h to 6.37 ± 0.02 at 2 h and 6.27 ± 0.02 at 3 h (Figure 4E), indicating progressive acidification of the external medium during the assay; matrix pH was not measured directly under these conditions.

### Substrate supplementation and ubiquinone competition reveal convergent electron supply

To test P6 and P7 (Table 1), we supplemented mitochondria with respiratory substrates and competed for electrons with exogenous ubiquinone (Figure 4C–G). The CII inhibitor malonate (10 mM) caused only modest inhibition (∼18.5%) in the absence of added respiratory substrates (Figure 4B). When succinate and fumarate were supplied together, malonate reduced total H₂ accumulation by 71.3 ± 7.0% relative to the same substrate condition without malonate (P < 0.01; Figure 4F). This malonate sensitivity is consistent with the model proposed in P6, in which succinate oxidation through Complex II supports matrix NADH regeneration via reverse electron transport (RET). Malate plus pyruvate (5 mM each), which generate matrix NADH via matrix dehydrogenases, enhanced H₂ 10.7 ± 1.8-fold (P < 0.001; Figure 4C); rotenone suppressed 68.7% of this enhancement, indicating that most of the supplemented reducing power requires Fe–S/quinone-branch turnover.

Succinate alone enhanced H₂ 11.1 ± 3.6-fold and fumarate alone 9.3 ± 1.8-fold; succinate plus fumarate (10 mM and 5 mM) enhanced H₂ 11.7 ± 1.2-fold (P < 0.001; Figure 4F). The comparable increases with malate plus pyruvate and with succinate are consistent with two routes that converge on the matrix NADH pool in the model (Figure 1C): direct NADH generation by matrix dehydrogenases and RET-dependent NADH regeneration supported by succinate oxidation.

Exogenous oxidized ubiquinone (UQ; 60 μM) suppressed H₂ to 6.9 ± 5.1% of the mitochondria-only control level (*P* < 0.01; Figure 4G), and reduced the mixed-acid stimulation by 70.7%; rotenone eliminated 97.8 ± 2.3% of the residual activity under ubiquinone competition, whereas malonate suppressed only 32.4 ± 2.5% (Figure 4G), consistent with **P7**: both supply routes converge on the same rotenone-sensitive, Complex I-associated step, and the oxidized quinone pool competes with the H₂-forming branch for electrons.

### Exploratory metabolomic screening reveals condition-dependent changes in carbon metabolism

To explore the metabolic processes associated with H₂ production, we performed an initial metabolomic screen of the crude mitochondrial fraction at 25 °C under N₂-flushed and air-started conditions (Figure 5—figure supplement 1). The preparation retained endogenous glycolytic and organic-acid intermediates and malate synthase activity (Figure 5—figure supplement 1B), indicating that glyoxylate-cycle-associated enzymes co-purified with the mitochondrial fraction. Although the six-carbon glycolytic intermediates changed relatively little between the two gas conditions, the three-carbon intermediates accumulated more strongly in the air-started samples. By contrast, N₂ flushing produced a prominent increase in succinate (∼95-fold at 10 h), accompanied by fumarate and malate accumulation, whereas pyruvate, isocitrate, and α-ketoglutarate increased preferentially in the air-started samples. Rotenone treatment substantially altered these patterns, consistent with disruption of redox-dependent carbon metabolism and TCA-cycle activity.

**Figure 5.**
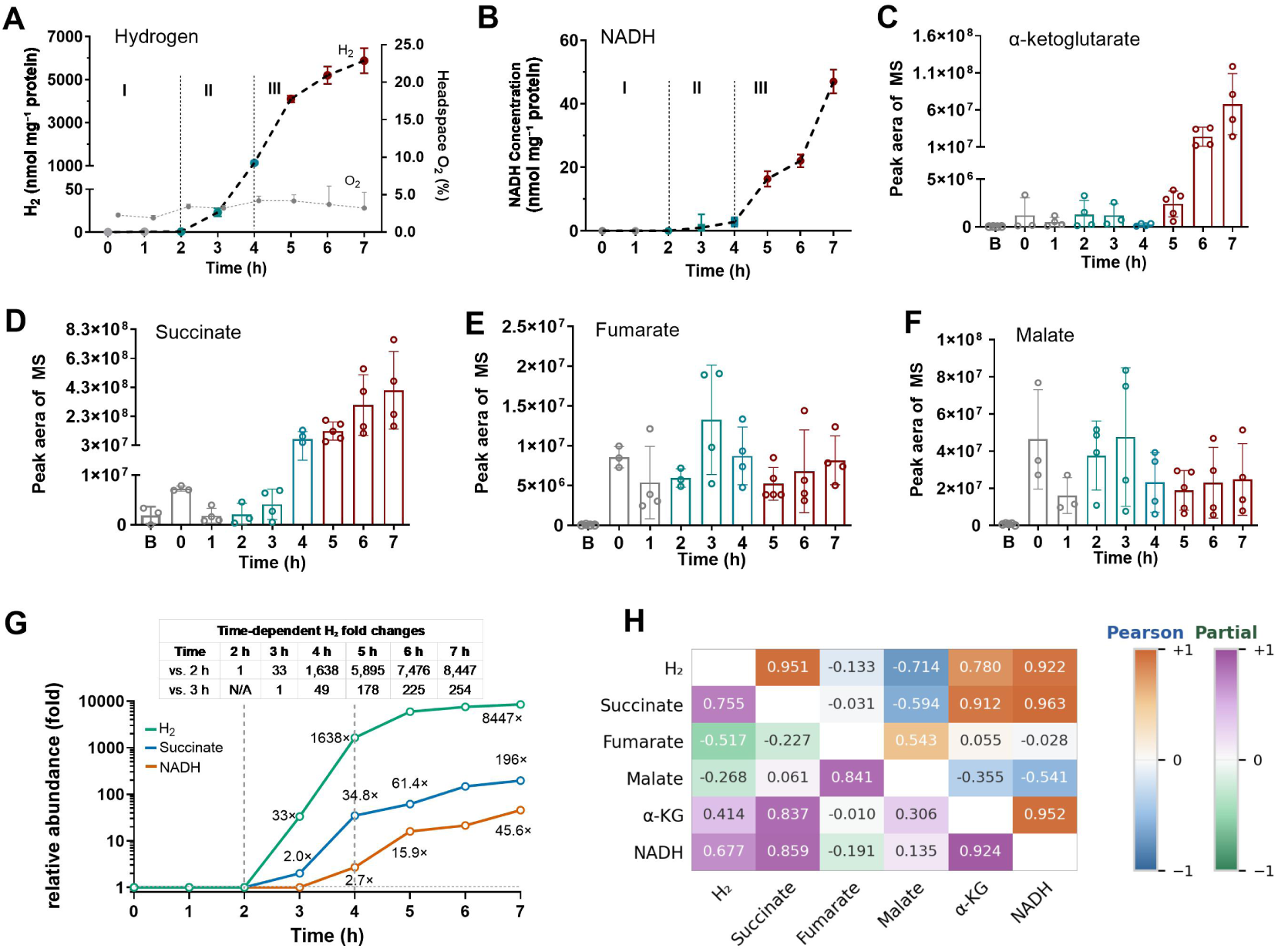
Time-resolved metabolic changes accompanying hypoxic H₂ production. Crude mitochondrial fractions (500 μg protein) isolated from *Vigna radiata* hypocotyls were incubated in Buffer H (500 μL) under hypoxia at 35 °C. Samples were collected hourly from 0 to 7 h for headspace H₂ and O₂ analysis by GC–BID, NADH quantification using the Amplite™ Colorimetric NADH Assay Kit (AAT Bioquest; absorbance at 460 nm), and targeted metabolite analysis by UHPLC–HRMS (Thermo Orbitrap; negative-ionization mode; resolution 30,000). Metabolites were identified by matching MS/MS spectra against the mzCloud database. Representative spectra and detailed analytical procedures are provided in Figure 5—figure supplement 2 and Methods. Dashed lines at 2 and 4 h define Phases I–III. **(A–F)** Time courses, under the same assay conditions, of headspace H₂ accumulation and residual O₂ (A), NADH concentration (B; background-corrected against the 0-h aliquot, see Methods), α-ketoglutarate peak area (C), succinate peak area (D), and fumarate and malate peak areas (E, F). **(G)** Relative time courses of H₂, succinate, and NADH. H₂ and succinate were normalized to their 2-h values, whereas NADH was normalized to its first detectable value at 3 h. Values before normalization are shown at 1× on the logarithmic ordinate. **(H)** Time-resolved correlations among H₂, NADH, and TCA-cycle intermediates. Upper-right triangle, Pearson correlations across the time course; lower-left triangle, partial correlations after controlling for linear time. Complete coefficients are provided in Supplementary Table S4. Data are means ± SD from three to five independent biological replicates derived from separate mitochondrial preparations. Where shown, error bars represent SD. In panels C–F, B denotes the extraction blank used for metabolomic background assessment. Metabolite abundances are expressed as mass-spectrometric peak areas in arbitrary units.

This exploratory dataset, particularly the pronounced succinate-associated response under N₂-flushed conditions, motivated a subsequent time-resolved metabolite quantification by UHPLC-HRMS analysis at 35 °C (Figure 5), designed to evaluate prediction P5 (Table 1).

### A three-phase metabolic cascade accompanies pronounced H₂ escalation

This 7-h time-course analysis resolved three operational phases of metabolic change accompanying hypoxic H₂ production (Figure 5A-F, Supplementary Table S3), evaluating P5. Throughout the assay, headspace O₂ remained low but detectable, demonstrating that H₂ accumulated under hypoxic rather than completely anoxic conditions (Figure 5A).

During Phase I (0-2 h), H₂ remained near baseline, NADH remained below the detection limit (LOD), and succinate remained low. α-Ketoglutarate, fumarate, and malate varied without sustained directional changes (Figure 5A–F).

During Phase II (2-4 h), pronounced H₂ accumulation began, increasing 33-fold at 3 h and 1,638-fold at 4 h relative to the 2-h value (49-fold by 4 h relative to 3 h). Succinate rose sharply at 4 h to approximately 34.8-fold above its 2-h level, coincident with a transient minimum in α-ketoglutarate, whereas NADH first became detectable at 3 h (1.03 nmol mg⁻¹ protein relative to the 0-h background; Figure 5B) and rose only to 2.81 nmol mg⁻¹ protein by 4 h (Figure 5A–D, G).

During Phase III (4-7 h), H₂ and succinate continued to accumulate, reaching 8,447- and approximately 196-fold above their respective 2-h values by 7 h. The 7-h H₂ level was 254-fold higher than that at 3 h. In contrast, NADH increased only from 5 h onward, reaching 47.0 nmol mg⁻¹ protein by 7 h. α-Ketoglutarate accumulated at 6-7 h after its transient minimum, while fumarate and malate showed no sustained directional trend (Figure 5B–G).

Correlation analysis distinguished temporal co-accumulation from time-independent metabolic associations (Figure 5H). H₂ correlated strongly with succinate and NADH across the time course (Pearson *r* = 0.951 and 0.922, respectively), and both associations were attenuated but retained after controlling for linear time (partial *r* = 0.755 and 0.677). Within the succinate–α-ketoglutarate–NADH module, all associations remained positive after time adjustment (partial *r* = 0.837-0.924), the α-ketoglutarate–NADH association being the strongest (partial *r* = 0.924), while the fumarate–malate association increased from Pearson *r* = 0.543 to partial *r* = 0.841 (Figure 5H; Supplementary Table S4).

Together, these results define a three-phase metabolic progression characterized by the onset of pronounced H₂ accumulation in Phase II, substantial succinate accumulation, and a delayed, comparatively modest rise in NADH. The time-independent association between succinate and NADH is consistent with P5 and with succinate-linked NADH regeneration through the quinone pool and reverse electron transport (Path 2, Figure 1C); likewise, the persistence of the H₂–succinate and H₂–NADH associations after time adjustment indicates that H₂ formation is linked to the succinate–NADH axis beyond shared temporal dynamics, consistent with sustained NADH/NAD⁺ cycling during active H₂ production.

## Discussion

Our findings suggest that a longstanding physiological phenomenon (H₂ release during seed germination) and an unresolved structural state of plant mitochondrial Complex I may represent two aspects of the same response to oxygen limitation: when respiratory demand exceeds oxygen availability, CI* may provide a condition-dependent platform for electron partitioning, allowing a fraction of excess reducing equivalents to be released as H₂.

### Localization, constraints, and model

Three levels of evidence support this conclusion. First, the H₂-forming activity co-fractionates with respiration-competent plant organelles, is heat-labile, and displays defined pH, temperature, and oxygen optima. Second, inhibitor and substrate-feeding experiments define the necessary conditions: sustained matrix NADH supply (R1), continued electron flow through the Complex I Fe–S/quinone branch (R2), and mildly acidic pH (R3). Third, these constraints are reproduced by an FMN–N1a branched electron-transfer model operating on the plant-specific Complex I assembly intermediate CI*. We do not claim direct proof of electron bifurcation; rather, we treat the branched pathway as a computation-grounded working model, distinct from the experimental constraints it explains.

### Rotenone and the logic of obligatory coupling

The response to rotenone is the most instructive observation of this study, because it excludes the simplest class of mechanisms. If H₂ arose from spontaneous discharge of accumulated FMNH₂, or from back-transfer of the N1a electron to FMNH· to form FMNH₂, blocking electron exit at the N2→ubiquinone step should leave H₂ unchanged or enhance it by enlarging the reduced-flavin pool. Instead, rotenone reduced H₂ to 1.32 ± 1.02% of control. Productive H₂ formation therefore depends on electron exit through the Fe–S chain. The branched model explains this in two steps. First, electron flow through the quinone branch cannot be interrupted. Rotenone removes the exergonic high-potential branch (FMNH₂ → N3 → UQ; ΔG ≈ −7.8 kJ mol⁻¹) that drives the low-potential transfer to N1a (ΔG ≈ +3.8 kJ mol⁻¹), and without N3 accepting the first electron, FMNH· cannot form; diverting both electrons directly to N1a is strongly unfavorable (ΔE = −129 mV; ΔG ≈ +12.4 kJ mol⁻¹). Both H₂ co-substrates, FMNH· and N1a(red), therefore disappear, freezing the cycle no matter how much FMNH₂ accumulates. Second, electron flow cannot be too rapid. The high-potential branch outcompetes the N1a route by roughly three orders of magnitude: FMN→N3 proceeds at ≈ 1.4 × 10⁹ s⁻¹, whereas FMN→N1a is ≈ 4.6 × 10⁵ s⁻¹, and N1a(red) back-transfers to the flavin at ≈ 1.4 × 10⁶ s⁻¹. The two one-electron donors therefore coexist only if the H₂-forming step captures N1a(red) before back-transfer. This dual requirement for electron flow that is neither interrupted nor uncontrolled mirrors bifurcating [FeFe]-hydrogenases, where blocking the high-potential acceptor abolishes H₂ (Buckel and Thauer 2018).

Inhibitors acting on opposite sides of the quinone pool are equally informative. Rotenone blocks electron entry, leaving the pool more oxidized; antimycin A and NaN₃ block electron exit, leaving it more reduced. If reductive pressure alone drove H₂, downstream blockers should have stimulated it; instead, they inhibited it, so H₂ formation requires continued drainage toward a partially oxidized quinone acceptor, which exogenous oxidized ubiquinone suppresses (Figure 4G). This convergence, in which blocking drainage abolishes H₂ and flooding the acceptor suppresses it, is the signature of obligatory coupling to quinone-branch turnover, and is difficult to reconcile with any uncoupled mechanism. The coupling is an experimental result; its realization through FMN–N1a electron bifurcation is a parsimonious molecular model, not proven chemistry. The azide data additionally remove the cytochrome branch’s proton-pumping contribution, a Δp-side consequence addressed below.

### A micro-oxic window sustained by the alternative oxidase

Continued drainage toward the quinone pool requires a terminal oxidant even when oxygen is scarce. Micro-oxia serves all three primary requirements at once: it preserves a residual oxidized-quinone fraction as the high-potential-branch acceptor (R2), limits NADH reoxidation (R1), and coincides with the progressive acidification that establishes R3; H₂ accumulation indeed proceeds under micro-oxic conditions with detectable residual O₂ (P9; Figures 2 and 3). The model adds a testable prediction: complete anoxia should suppress H₂ formation by starving the acceptor branch, whereas higher oxygen availability should dissipate the reduced state the chemistry requires, so activity should peak at an intermediate oxygen tension. This non-monotonic dependence distinguishes the model from any scheme in which H₂ is a simple endpoint of progressive over-reduction. A hint of this behavior is already visible in intact seedlings: under the same initial low-O₂ treatment, roots accumulated H₂ to a level comparable to that of cotyledons plus leaves while retaining the highest residual O₂ (1.37%, versus 0.82% and 0.63% in cotyledons plus leaves and hypocotyls, respectively) (Figures 2H and 2I). Organ-specific differences in metabolism and gas diffusion preclude inferring an oxygen optimum from this comparison, but the pattern argues against H₂ production as a simple monotonic consequence of oxygen depletion. If an optimum does lie within the micro-oxic range typical of oxygen-limited root tissues, this pathway could serve a physiological role that controlled O₂-gradient experiments can now test.

Under these conditions the alternative oxidase provides the indispensable quinol-oxidizing outlet: SHAM reduced H₂ below the LOD whereas antimycin A inhibited only partially (Figure 4B; P3–P4). This is consistent with the known induction of AOX under low oxygen (Millar et al. 2011; Vanlerberghe 2013), and with micro-electrode recordings showing steep radial O₂ gradients and oxygen-impermeable barriers in wetland plant roots (Armstrong 2000). It follows from the energetics of the hypoxic state. Complex IV retains catalytic competence at the residual oxygen tensions observed here (sub-micromolar Km for O₂; Rich 2003). As a proton-pumping oxidase, however, it is restrained by the Δp it builds (Brown 1992; Kadenbach 2003), whereas the non-pumping AOX escapes this back-pressure and provides a non-coupled outlet when Δp is high(Gupta, Zabalza, and Van Dongen 2009). Flux through the proton-coupled cytochrome branch may thus contribute to the residual Δp that powers reverse electron transfer, while AOX keeps the quinone pool partly oxidized. The partial effect of antimycin A, in contrast to the complete abolition by SHAM and azide, also has precedent in this developmental window (Kennedy, Fox, and Rumpho 1987); the basis of this difference is addressed in the next section.

### NADH cycling and a RET-compatible supply route

Sustained electron exit solves only half the problem; the supply side must keep pace. The results establish that H₂ formation is driven by direct electron delivery from the matrix NADH pool to the CI*-bound FMN rather than by external NADH oxidation (R1; Figures 4C, D and F). Exogenous NADH, even at 1 mM, increased H₂ accumulation only ∼1.4-fold. Oxygen-electrode assays confirmed that this NADH was indeed oxidized, indicating that externally supplied NADH feeds the quinone pool via external NADH dehydrogenases rather than the matrix-facing CI* flavin. A simple stoichiometric consideration shows that this supply must be cyclic: cumulative H₂ production (5879 nmol mg⁻¹ protein) exceeded the maximal measured NADH level (47 nmol mg⁻¹ protein) by roughly two orders of magnitude, meaning the NADH pool could not have acted as a static reservoir if NADH supplies the electrons, so that continuous NADH/NAD⁺ cycling would be required.

The cycle is closed by the coexistence of the two Complex I species in the same mitochondrial preparations (Ligas et al. 2019; Maldonado et al. 2020): In this model, CI* oxidizes matrix NADH at its exposed flavin, releasing reducing equivalents as H₂ and regenerating NAD⁺, which is reused by matrix dehydrogenases and by succinate-driven reverse electron transport (RET) (Murphy 2009; Rustin and Lance 1991). Two NADH-generating routes converge on this shared pool: direct TCA-cycle dehydrogenation, fed in part by succinate exported from the glyoxysomal glyoxylate cycle (Eastmond and Graham 2001; Graham 2008), and RET, which is thought to operate primarily in the mature holoenzyme; whether CI* itself can sustain RET is not established. This division of labour also precludes futile electron oscillation between the flavin and the quinone pool within a single enzyme.

Succinate-driven RET through Complex I is established in mammals, where it underlies ischaemia–reperfusion ROS production (Chouchani et al. 2014; Murphy 2009). In plants, biochemically indicative evidence has long been available (Rustin and Lance 1991; Storey 1971), but its physiological relevance is not established: Huang et al. (2019) list RET at Complex I among the key unanswered questions of the field, noting the parallel between succinate accumulation during waterlogging or anoxia in plants and reperfusion in mammals. The convergent pattern reported here offers a plausible route into this open question. Succinate and fumarate enhanced H₂ accumulation 11.1- and 9.3-fold, respectively, and malonate suppressed the combined stimulation by 71%, tying the effect to succinate oxidation through Complex II (Figure 4F; P6). The metabolite correlations tell the same story: the α-ketoglutarate–NADH and succinate–NADH associations survive statistical control for linear time (partial r = +0.924 and +0.858), and H₂ itself remains positively associated with succinate and NADH after the same control (partial r = +0.755 and +0.677, respectively; Figure 5H; P5). Both supply routes thus converge on the shared matrix NADH pool that feeds the same FMN site (Figures 4C, F and G; P7).

We emphasize that these data establish a RET-compatible supply route, not direct reverse electron flux. They also suggest a possible explanation for why plant RET has been difficult to detect: rather than producing a burst of ROS at Complex I, as in mammals, venting reductive pressure as H₂ could reduce or alter the corresponding ROS signature. This explanation is testable by simultaneously measuring H₂ flux and site-specific ROS production under RET-promoting conditions. Both the presence of RET and its proposed masking by H₂ require direct determination of NADH formation, quinone redox state and Δp dependence.

### Protonmotive force and matrix acidity define the permissive window

In our model, the protonmotive force (Δp) supports H₂ formation indirectly by driving reverse electron transport (RET) through mature Complex I and thereby replenishing matrix NADH (R1). FCCP reduced H₂ accumulation below the LOD, whereas oligomycin did not significantly alter it (134.9 ± 47.2% of control, P = 0.33; Figure 4B; P2). This identifies Δp, rather than ATP synthesis, as the permissive factor. Because dissipating Δp can only suppress a Δp-dependent process, the response also indicates that a protonmotive force was present under the assay conditions. Dissipation of Δp would curtail RET-dependent NADH regeneration while relieving thermodynamic back-pressure on forward electron transfer through Complex I. Both effects would favor depletion of the matrix NADH pool (Supplementary Calculation S2.5).

The oligomycin result bears on a separate question: whether ATP synthase activity contributes to H₂ accumulation under our assay conditions. De Col et al. (2017) reported that cytosolic MgATP²⁻ declined slowly over several hours before a sharp decrease during progressive hypoxia. Those whole-cell measurements do not support the inference that Complex V was already inactive in our mitochondrial preparations. Conversely, the absence of a significant oligomycin effect does not show that Complex V maintained Δp. The differential effect of antimycin A and azide on H₂ formation reflects their distinct impact on Δp maintenance. In mitochondria from anaerobically germinated Echinochloa seedlings, respiration is barely inhibited by antimycin A, indicating a low-flux or structurally altered Complex III under these conditions (Kennedy et al. 1987). If the same applies to our hypoxic system, antimycin A would leave Complex III and IV pumping partially intact; together with site-1 pumping at mature Complex I (retained under alternative-pathway respiration; Millar et al. 2011), Δp would decline but not collapse, attenuating RET-driven NADH regeneration and reducing H₂ by only 38.1 ± 11.3%, to 61.9 ± 11.3% of the control level (Figure 4B), rather than abolishing it. Azide, by contrast, blocks Complex IV completely; at the 3-h measurement window, where the system plausibly sits close to the Δp threshold for RET (Figure 5A), this complete block would collapse RET-driven NADH regeneration and abolish H₂. SHAM and azide thereby act as complementary probes along the same chain: the former compromising quinol oxidation while the cytochrome branch still pumps, the latter compromising cytochrome-branch pumping while AOX-mediated quinol oxidation persists (Bendall and Bonner 1971).

The pH dependence adds a second dimension to this window. A fall in external pH enlarges the transmembrane pH gradient and hence the inward driving force on protons; when respiratory proton pumping cannot sustain Δp, as under hypoxia, this gradient is dissipated by proton influx and the matrix follows external acidification at least partially (Douce, 1985). Consistent with this, cytosolic pH falls to ≈6.8 in hypoxic maize root tips (Roberts et al., 1984) and anoxic sycamore cells (Gout et al., 2001; Felle, 2001), and hypoxia is associated with matrix acidification in plants (Ramírez-Aguilar et al., 2011). As matrix pH was not measured here, these comparisons establish plausibility rather than a direct test. Mechanistically, the low-potential branch crosses thermodynamic neutrality near pH 6.7 (R3; Supplementary Calculation S4). Because the cooperative H₂-forming step consumes a matrix proton whereas the direct FMNH₂ route is pH-neutral, mild acidification favours the cooperative pathway (P8), tempered by any pH dependence of the donor couple (S10). The crossover position also depends on the proton-coupled stoichiometry and the actual H₂ partial pressure (S5, S10). External buffer pH 6.0 is an experimental condition under which H₂ accumulation was maximal; the corresponding matrix pH was not measured. Whether this acidic optimum acts through the Δp/ΔpH it imposes or through bulk-medium protonation remains unresolved; FCCP reduced H₂ accumulation below the LOD at pH 6.0 (Figure 4B), indicating that maintenance of Δp is required under these conditions. The descending limb is provisionally attributed to the combined burden of extreme acidity on protein stability and on Δp maintenance (Figure 3I).

The chemical locus of H–H bond formation remains open between the two variants introduced in the Results: a cooperative FMN-N5/N1a site, with N1a transiently donating the second electron (Model A, favored as the primary working model), and a transiently exposed N1a iron acting through a metal-hydride route (Model B). Both are thermodynamically permissible and consistent with the mildly acidic optimum; they are separated by mutational and spectroscopic tests outlined in Supplementary Calculation S13.

### CI* as the plausible catalytic platform

Mature CI contains the same FMN and N1a cofactors as CI*, yet accumulates no detectable H₂, suggesting that the holoenzyme architecture suppresses rather than supports this reaction. In mature CI, incorporation of the PD module may reduce solvent access to FMN-N5 and matrix proton delivery, while tight coupling to proton pumping may limit semiquinone accumulation. Preferential incorporation into closed I+III₂ supercomplexes may further restrict quinone exchange and access to the Q module (Klusch et al. 2021). By contrast, CI* retains the N and Q peripheral-arm modules, the proximal membrane (PP) module and the plant-specific γ-carbonic-anhydrase domain, but lacks the distal PD module (Ligas et al. 2019; Maldonado et al. 2020; Soufari et al. 2020). Thus, the combined requirements for an exposed FMN, proximity to N1a, the lack of the distal PD module and its associated proton-pumping coupling, and compatibility with acidic, micro-oxic conditions converge on CI* as the most plausible catalytic platform.

The dominant CI*-like particle isolated from etiolated mung bean lacks the assembly factor GLDH, which is incorporated late together with the distal PD module and is not present in mature Complex I (Ligas et al. 2019; Schimmeyer et al. 2016). GLDH absence is therefore equally compatible with an assembly intermediate and with a particle derived from the holoenzyme, and does not by itself discriminate between them. We follow Maldonado et al. (2020) in assigning the particle to the CI* intermediate on the basis of their convergent biochemical and structural evidence. Isolated GLDH-deficient particles retain NADH-oxidation activity both after BN-PAGE (Maldonado et al. 2020; Schertl et al. 2012) and in solution-phase NADH:decylubiquinone assays (Maldonado et al. 2020), indicating that the cofactors and subunits required for NADH oxidation are intact in the isolated complex. This catalytic competence, however, does not establish that CI* oxidizes NADH in vivo; it demonstrates only that the redox machinery necessary for the proposed reaction is preserved in the purified particle. If the proposed activity is validated, H₂ evolution would provide a potential physiological function for this otherwise poorly understood intermediate.

Structural comparisons further support the feasibility of electron transfer within CI*. NDUV1 and NDUV2 superimpose on HydB and HydC of the bifurcating [FeFe]-hydrogenase HydABC with RMSD values of 1.04 and 1.15 Å, respectively, and the HydC C1 cluster involved in bifurcation corresponds to N1a (Furlan et al. 2022). N1a lies approximately 11.8 Å from FMN-N5 (Maldonado et al. 2020, Figure 5A; Supplementary Calculation S8.5), and mutational analyses of bacterial Complex I show that N1a is dispensable for steady-state NADH oxidation yet participates in redox-dependent conformational changes (Dörner et al. 2017). Moser–Dutton analysis predicts electron-transfer rates of approximately 1.4 × 10⁹ s⁻¹ for FMN→N3 and 4.6 × 10⁵ s⁻¹ for FMN→N1a, both substantially faster than the measured H₂ flux (Moser et al. 1992; Supplementary Calculation S8). N1a may also participate in redox-dependent conformational regulation, as suggested by the peptide flip observed near the NADH-binding site in bacterial Complex I (Schulte et al. 2019); here, this feature is proposed as a testable possibility rather than an established mechanism. These structural relationships support the feasibility of electron transfer but do not identify the H–H bond-forming site, which remains unresolved between Models A and B.

Several indirect lines associate H₂ evolution with CI* rather than mature CI. H₂-evolving activity is maximal in etiolated tissues at the developmental stage when plant mitochondria are known to accumulate the CI* intermediate (Ligas et al. 2019; Maldonado et al. 2020; Meyer, Letts, and Maldonado 2022). Sustained hypoxia and acidification also promote a shift from supercomplex-assembled to free forms of plant Complex I (Ramírez-Aguilar et al. 2011), conditions that are not restricted to seedlings. CI* occurs in non-etiolated Arabidopsis thaliana seedlings and leaves, most abundantly in young seedlings (Ligas et al. 2019). Hypoxic tissues of mature plants may therefore support H₂ evolution as well, which remains to be tested. Similar activity detected in other plant species supports a conserved CI*-associated mechanism rather than a mung-bean-specific phenomenon.

No single observation establishes CI* as the catalytic species. However, the convergence of structural, biochemical, developmental and fractionation evidence makes CI* the most parsimonious platform consistent with the available constraints. Direct co-fractionation of H₂-evolving activity with CI* and selective destabilization of the intermediate will provide critical tests of this assignment.

### Self-amplifying dynamics and the Phase I–III transition

Assigning the activity to CI* also explains its most conspicuous kinetic feature: a three-phase temporal profile (Figure 5A–G). Phase I is a long permissive interval of hours-long near-zero accumulation, consistent with the progressive establishment of a micro-oxic, acidic and sufficiently reducing state. Phase II marks the onset of pronounced H₂ accumulation (33-fold at 3 h), when NADH first becomes detectable but remains near baseline (1.03 nmol mg⁻¹ protein; Figure 5B), before succinate rises (4 h). In Phase III, H₂ escalates 254-fold between 3 and 7 h, while the NADH pool builds only from 5 h onward, reaching ≈46-fold relative to its first detectable value. This staggering is informative: NADH accumulation lags far behind H₂ escalation, and its small, late increase against the cumulative output is consistent with rapid turnover rather than static accumulation.

The loop is self-sustaining: H₂ formation consumes NADH, while continued upstream regeneration replenishes the pool and sustains further H₂ formation. If NADH supply depended on a single linear route, inhibiting that route should strongly suppress H₂ evolution; the partial effects instead support parallel contributions from malate-driven dehydrogenase flux and the succinate-driven RET route. The former, being enzymatic rather than Δp-powered, should be less Δp-sensitive than the latter, and the observed additive pattern is consistent with this prediction (Figures 4C and F). Because Δp was not measured directly, this interpretation remains inferential (see Limitations). More generally, each requirement gates the next (succinate oxidation → NADH regeneration → FMNH₂ formation → branched electron transfer → H₂), so no single requirement can compensate fully for the others. The proposed pathway is not directly ATP-coupled; its feasibility is governed primarily by internal redox poise and pH (Supplementary Calculation S10.4). Phase III therefore marks the point at which these conditions stabilize together, allowing modest correlated changes to compound into large increases in H₂ output.

### Physiological implications

Seed germination may represent one of the most physiologically relevant contexts in which this latent pathway is engaged. Upon imbibition, respiratory demand rises abruptly, yet oxygen diffusion remains limited by the hydrated seed matrix, seed coat and densely packed storage tissues, so germinating seeds readily develop steep internal oxygen gradients and transient micro-oxic niches. Rapid mobilization of stored carbon reserves (via β-oxidation and the glyoxylate cycle in lipid-storing seeds, or parallel reserve routes in starch- and protein-storing seeds) simultaneously increases mitochondrial reducing pressure, while incomplete oxidation under low oxygen favours accumulation of organic acids and succinate. These conditions match those predicted to favour CI*-dependent H₂ evolution: elevated NADH, succinate-supported reverse electron transport, low effective O₂ availability, AOX engagement and acidification. Mitochondrial biogenesis and respiratory-complex assembly are also maximal during this window. The burst of mitochondrial biogenesis and membrane proliferation during the first 24-72 h of germination (Law, Narsai, and Whelan 2014; Nawa and Asahi 1971) therefore predicts a transiently high CI*:mature-Complex I ratio at exactly the stage of maximal H₂ evolution, suggesting that the classic H₂ emission of germinating seeds (Renwick et al. 1964) reflects not an oddity of seeds but a developmental window in which the conditions for an alternative mitochondrial electron outlet coincide.

The glyoxylate cycle is highly active during seed germination (Eastmond and Graham 2001; Graham 2008), and glyoxysomes can co-sediment with mitochondria during subcellular fractionation. Consistent with this, our crude mitochondrial preparations retain malate synthase activity, a glyoxylate-cycle marker enzyme, together with endogenous organic-acid pools. The metabolite profiles show condition- and time-dependent changes in malate, succinate and other intermediates (Figure 5—figure supplement 1B), consistent with continued turnover of endogenous redox-active metabolites. These observations support retention of endogenous metabolic capacity, but do not by themselves establish mitochondrial localization of malate synthase. Together with heat lability, defined pH and temperature optima, and coherent responses to mechanistically distinct perturbations, they are difficult to reconcile with a simple non-metabolic source of H₂.

The physiological stakes are sharpest in this same window. Metabolic flux rises before the respiratory chain is fully stabilized, creating redox vulnerability and, upon reoxygenation, a risk of oxidative damage. Diverting excess reducing equivalents to H₂ could provide a benign outlet for reductive pressure, limiting over-reduction of the mitochondrial redox network while preserving metabolic continuity. This contrasts with animal mitochondria, where equivalent reductive pressure during ischaemia is vented as superoxide at the FMN site (Chouchani et al. 2014). The extensively documented ability of exogenous H₂ to enhance antioxidant defences and stress tolerance (Jin et al. 2016; Xie et al. 2012), may partly reflect engagement of an endogenous H₂-generating system under hypoxia; whether CI*-derived H₂ acts as a local signal or protective molecule during hypoxia–reoxygenation remains to be tested. We interpret H₂ evolution as a conditional alternative electron-transfer route that operates within the mitochondrial redox-poise system and competes with conventional quinone reduction for reducing equivalents.

### Evolutionary implications

The N-module of Complex I is widely accepted to derive from an ancestral hydrogenase-related electron-transfer module that predated the acquisition of the membrane proton-pumping modules (Efremov and Sazanov 2012; Friedrich and Scheide 2000). In this light, the fully assembled enzyme represents the derived, constrained state, whereas the plant-specific CI* intermediate retains a less restricted and more ancestral redox architecture. This difference is made concrete by assembly order: plants add the distal membrane module last and therefore accumulate a stable NADH-competent intermediate, whereas animals add the N-module last and never expose the ancestral module as a stable species (Ligas et al. 2019; Maldonado et al. 2020). The structural correspondence between the CI* N-module and the bifurcation module of [FeFe]-hydrogenases further places this intermediate within the same enzyme family that couples a low-potential electron to proton reduction (Furlan et al. 2022). Although CI* lacks an H-cluster, it retains the ancestral electron-partitioning module; under high electron pressure, acidic pH, and increased conformational access, that module may support condition-dependent transfer of its low-potential electron to protons. Given the hydrogen-rich, reducing conditions of the early Earth, proton reduction would have been a chemically accessible electron sink; plant CI* may therefore retain an ancestral electron-partitioning capacity that is conditionally expressed as H₂ evolution, rather than a newly evolved function.

The phylogenetic distribution of this activity may therefore reflect differences in Complex I assembly-state stability, FMN-site regulation, and alternative electron-transfer pathways among lineages. The absence of reported mitochondrial H₂ evolution in animals more likely reflects differences in the stability or abundance of relevant assembly states, respiratory coupling, or the availability of alternative electron sinks. The retention of both AOX and a transient CI* intermediate in plants may thus represent a lineage-specific respiratory strategy suited to recurrent oxygen limitation during seed imbibition, germination in compact or waterlogged soils, and later flooding stress. Alongside the cytochrome pathway, AOX, and ROS-generating electron leak, CI*-dependent H₂ evolution may provide a further electron outlet that buffers mitochondrial over-reduction during hypoxia and reoxygenation.

### Limitations of the study

Several considerations should be kept in mind when interpreting our findings.

First, the evidence linking H₂ production to the CI* assembly intermediate remains indirect. Our assignment is guided by three considerations: (i) CI* is a recurrently detected assembly intermediate that retains the FMN, Fe–S chain and N1a cofactors while lacking the distal PD module (Ligas et al. 2019; Maldonado et al. 2020); (ii) its more open conformation leaves the FMN-N5 region more solvent-exposed, providing a structural rationale for the FMN-centered variant of the model; and (iii) the model proposes that assembly of the distal PD module and incorporation into supercomplexes disfavor this chemistry in mature Complex I. However, we have not directly identified CI* as the H₂-producing species. The present experiments cannot distinguish its proposed contribution from possible contributions by mature Complex I, other partially dissociated Complex I species, or mitochondrial ferredoxin. Attempts to reconstitute H₂ production from isolated Complex I subcomplexes or purified components have not yielded activity under the conditions tested. This is compatible with, but does not establish, the proposal that H₂ formation requires an intact membrane context involving AOX activity, maintained protonmotive force and appropriate quinone-pool redox poise. Plant lines that shift the CI:CI* ratio or selectively destabilize CI*, particularly in an inducible or tissue-specific manner, would provide more direct perturbations. Reconstitution in a membrane environment preserving the relevant interactions would provide an independent test.

Second, metabolite profiling employed relative quantification by UHPLC-HRMS without stable-isotope internal standards. Absolute concentrations are therefore not determined, but the temporal trends, particularly the dissociation of the succinate and H₂ surges from NADH accumulation, derive from identically processed samples normalized to initial time points, ensuring internal comparability.

Third, the pH optimum was determined in externally buffered assays, and matrix pH and Δp were not measured directly; conclusions about pH- and Δp-dependent steps rest on indirect evidence, although the pH-6 optimum is compatible with the matrix acidification documented under hypoxia (Ramírez-Aguilar et al. 2011). Direct measurement of matrix pH during H₂ production would further test our thermodynamic switch hypothesis. The pharmacological toolkit also has known caveats: SHAM can inhibit other flavoproteins at high concentrations (Vanlerberghe 2013), azide may act as a weak-acid proton shuttle, and malonate inhibition of Complex II is competitive and incomplete in intact systems.

Despite these limitations, the inhibitor fingerprint and kinetic constraints established here, particularly the rotenone paradox, which argues against the simplest uncoupled mechanisms considered here, provide a rigorous biochemical framework that any future candidate mechanism must satisfy. We therefore present CI* as the most parsimonious platform consistent with the available biochemical, structural and thermodynamic data, while recognizing that its catalytic assignment remains unresolved.

### Falsifiable predictions

If the CI* branched electron-transfer model is correct, H₂ production should scale with CI* abundance and co-fractionate with CI* in native electrophoresis. Blocking electron exit through the Fe–S/quinone branch should suppress rather than enhance H₂ evolution, and succinate stimulation should depend on a Complex II-dependent, Δp-sensitive supply route. H₂ production should further depend on a micro-oxic, AOX-supported redox state and increase under acidic conditions. At the molecular level, branched electron-transfer chemistry should carry a flavin semiquinone EPR signature under H₂-producing conditions, and isotope tracing with ¹³C-succinate and ¹³C-malate should define the carbon fluxes sustaining NADH regeneration. Selective destabilization of CI* without disrupting mature Complex I should reduce H₂ evolution, and reconstitution of the activity in a membrane context retaining AOX and Δp should be required for catalytic competence. Failure of these predictions would argue against the proposed mechanism.

## Conclusion

Our results establish a mitochondrial, Complex I-associated origin for hypoxic H₂ production in vascular plants and define its gating requirements: Fe–S/quinone-branch turnover, succinate-supported reverse electron transport, AOX activity, protonmotive force, low oxygen, and acidic pH. The rotenone paradox, whereby quinone-site blockade abolishes rather than enhances H₂, eliminates simpler non-bifurcating mechanisms and is the defining signature of this activity. This activity may depend on the intact membrane context, rather than a function of any single isolable enzyme, which may explain why it eluded discovery for six decades. While direct confirmation of CI* as the catalytic entity awaits future genetic or structural studies, our findings reveal a latent hydrogenase-like potential within the plant mitochondrial respiratory chain. Complex I assembly may not merely build the respiratory machinery; under specific physiological conditions, its intermediate states may reshape electron fate, making assembly state a previously unrecognized candidate determinant of electron partitioning under hypoxia.

## Materials and methods

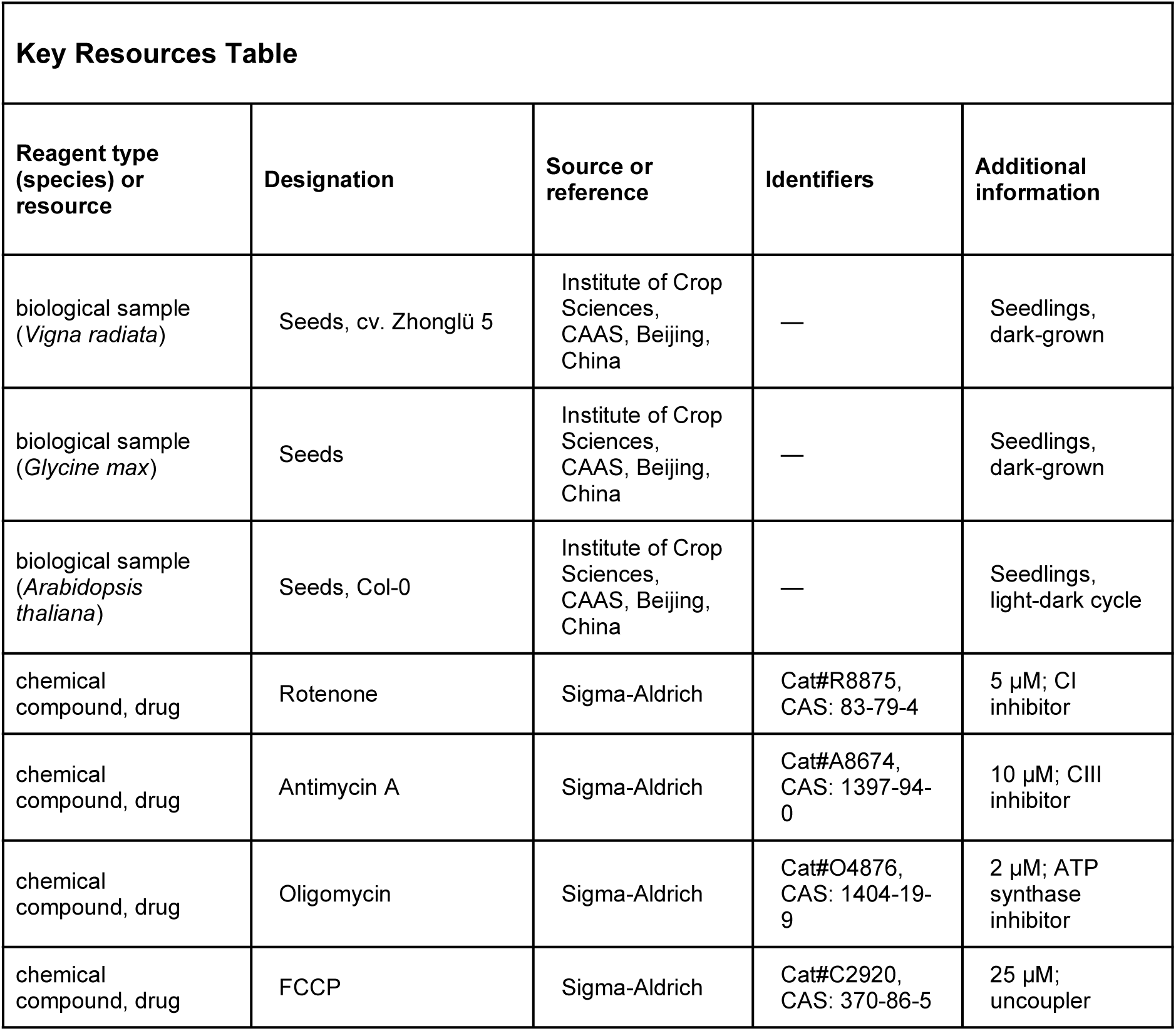
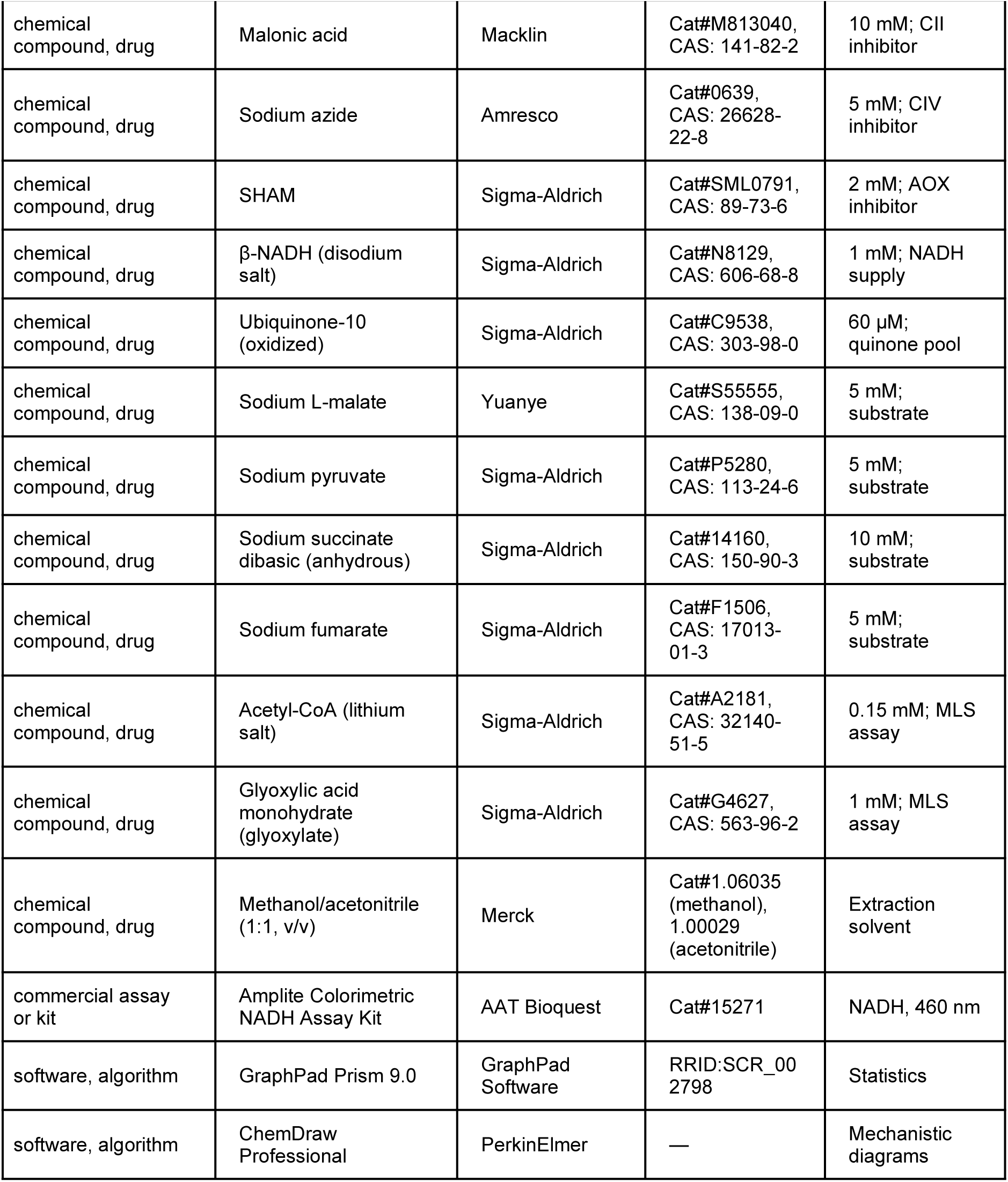
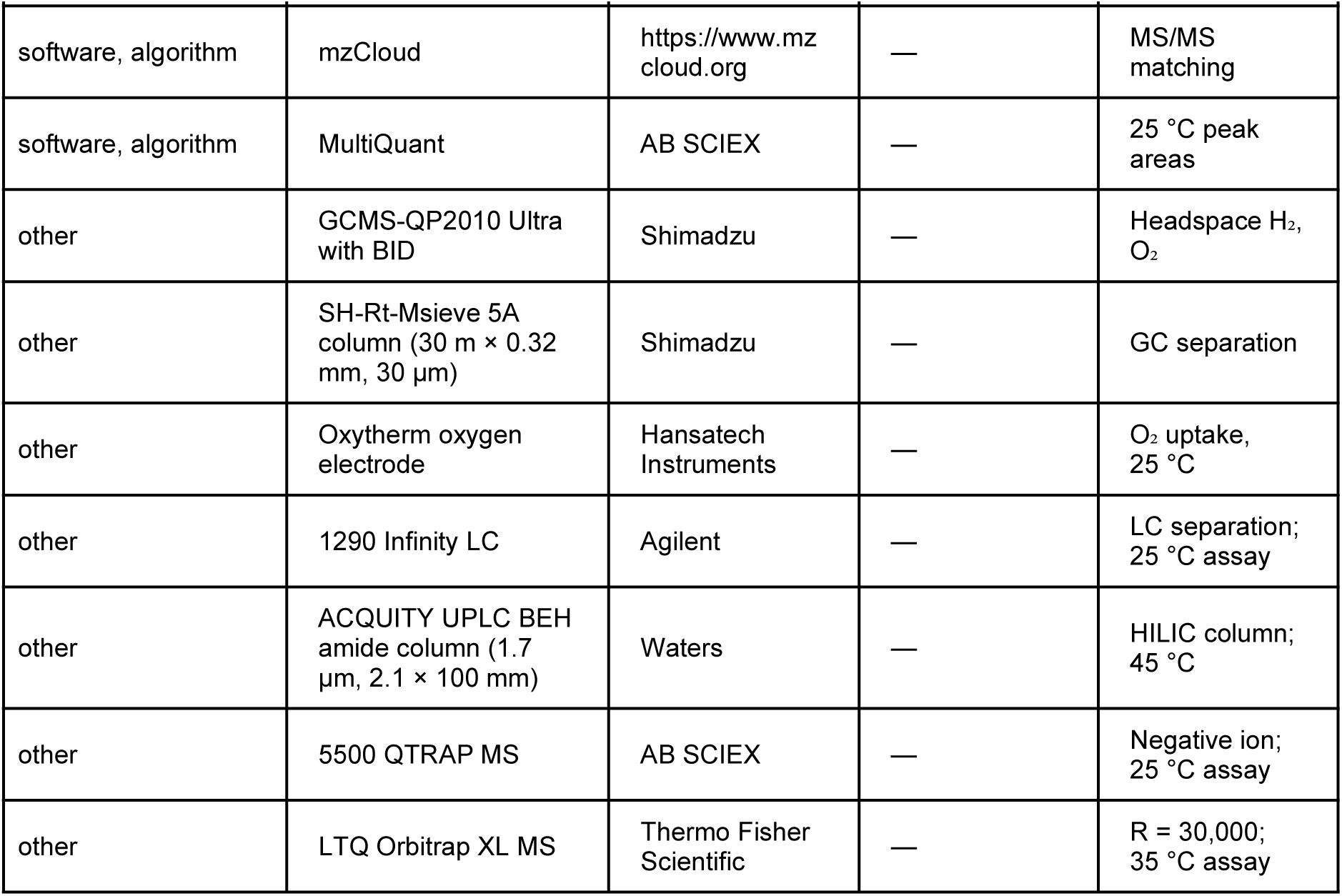

| Key Resources Table |  |  |  |  |
|---|---|---|---|---|
| Reagent type (species) or resource | Designation | Source or reference | Identifiers | Additional information |
| biological sample (*Vigna radiata*) | Seeds, cv. Zhonglü 5 | Institute of Crop Sciences, CAAS, Beijing, China | — | Seedlings, dark-grown |
| biological sample (*Glycine max*) | Seeds | Institute of Crop Sciences, CAAS, Beijing, China | — | Seedlings, dark-grown |
| biological sample (*Arabidopsis thaliana*) | Seeds, Col-0 | Institute of Crop Sciences, CAAS, Beijing, China | — | Seedlings, light-dark cycle |
| chemical compound, drug | Rotenone | Sigma-Aldrich | Cat#R8875, CAS: 83-79-4 | 5 µM; CI inhibitor |
| chemical compound, drug | Antimycin A | Sigma-Aldrich | Cat#A8674, CAS: 1397-94-0 | 10 µM; CIII inhibitor |
| chemical compound, drug | Oligomycin | Sigma-Aldrich | Cat#O4876, CAS: 1404-19-9 | 2 µM; ATP synthase inhibitor |
| chemical compound, drug | FCCP | Sigma-Aldrich | Cat#C2920, CAS: 370-86-5 | 25 µM; uncoupler |
| chemical compound, drug | Malonic acid | Macklin | Cat#M813040, CAS: 141-82-2 | 10 mM; CII inhibitor |
| chemical compound, drug | Sodium azide | Amresco | Cat#0639, CAS: 26628-22-8 | 5 mM; CIV inhibitor |
| chemical compound, drug | SHAM | Sigma-Aldrich | Cat#SML0791, CAS: 89-73-6 | 2 mM; AOX inhibitor |
| chemical compound, drug | β-NADH (disodium salt) | Sigma-Aldrich | Cat#N8129, CAS: 606-68-8 | 1 mM; NADH supply |
| chemical compound, drug | Ubiquinone-10 (oxidized) | Sigma-Aldrich | Cat#C9538, CAS: 303-98-0 | 60 µM; quinone pool |
| chemical compound, drug | Sodium L-malate | Yuanye | Cat#S55555, CAS: 138-09-0 | 5 mM; substrate |
| chemical compound, drug | Sodium pyruvate | Sigma-Aldrich | Cat#P5280, CAS: 113-24-6 | 5 mM; substrate |
| chemical compound, drug | Sodium succinate dibasic (anhydrous) | Sigma-Aldrich | Cat#14160, CAS: 150-90-3 | 10 mM; substrate |
| chemical compound, drug | Sodium fumarate | Sigma-Aldrich | Cat#F1506, CAS: 17013-01-3 | 5 mM; substrate |
| chemical compound, drug | Acetyl-CoA (lithium salt) | Sigma-Aldrich | Cat#A2181, CAS: 32140-51-5 | 0.15 mM; MLS assay |
| chemical compound, drug | Glyoxylic acid monohydrate (glyoxylate) | Sigma-Aldrich | Cat#G4627, CAS: 563-96-2 | 1 mM; MLS assay |
| chemical compound, drug | Methanol/acetonitrile (1:1, v/v) | Merck | Cat#1.06035 (methanol), 1.00029 (acetonitrile) | Extraction solvent |
| commercial assay or kit | Amplite Colorimetric NADH Assay Kit | AAT Bioquest | Cat#15271 | NADH, 460 nm |
| software, algorithm | GraphPad Prism 9.0 | GraphPad Software | RRID:SCR_002798 | Statistics |
| software, algorithm | ChemDraw Professional | PerkinElmer | — | Mechanistic diagrams |
| software, algorithm | mzCloud | https://www.mzcloud.org | — | MS/MS matching |
| software, algorithm | MultiQuant | AB SCIEX | — | 25 °C peak areas |
| other | GCMS-QP2010 Ultra with BID | Shimadzu | — | Headspace H<sub>2</sub>, O<sub>2</sub> |
| other | SH-Rt-Msieve 5A column (30 m × 0.32 mm, 30 µm) | Shimadzu | — | GC separation |
| other | Oxytherm oxygen electrode | Hansatech Instruments | — | O<sub>2</sub> uptake, 25 °C |
| other | 1290 Infinity LC | Agilent | — | LC separation; 25 °C assay |
| other | ACQUITY UPLC BEH amide column (1.7 µm, 2.1 × 100 mm) | Waters | — | HILIC column; 45 °C |
| other | 5500 QTRAP MS | AB SCIEX | — | Negative ion; 25 °C assay |
| other | LTQ Orbitrap XL MS | Thermo Fisher Scientific | — | R = 30,000; 35 °C assay |

### Plant materials and growth conditions

Mung bean (*Vigna radiata*) and soybean (*Glycine max*) seeds were surface-sterilized in 2% (v/v) sodium hypochlorite for 15 min, followed by 75% (v/v) ethanol for 5 min, and then rinsed three times with sterile distilled water. Seeds were incubated in sterile distilled water at 55 °C for 20 min and subsequently soaked in darkness at 25 ± 1 °C for 8 h. After soaking, the seeds were placed on two layers of sterile gauze in seedling trays and germinated in darkness at 25 ± 1 °C.

Unless otherwise specified, mung bean hypocotyls were collected from 72-96-h-old etiolated seedlings. Roots, hypocotyls, and cotyledons together with true leaves were excised from 72-h-old seedlings for organ-specific H₂ assays. Mung bean was selected for biochemical characterization because its relatively high H₂ signal provided a sufficient signal-to-background ratio for inhibitor, kinetic, and assay-optimization experiments.

*Arabidopsis thaliana* Col-0 seeds were surface-sterilized in 75% (v/v) ethanol for 5 min and washed three times with sterile distilled water. Seedlings were grown aseptically on half-strength Murashige and Skoog medium under a 16-h-light/8-h-dark photoperiod at 23 °C during the light period and 18 °C during the dark period. Roots from 1-week-old soybean seedlings and 2-week-old *A. thaliana* seedlings were collected for preparation of crude mitochondrial fractions.

### Isolation of subcellular fractions

All fractionation procedures were performed at 4 °C. Hypocotyls (100 g) from 72-96-h-old mung bean seedlings were homogenized in 300 mL ice-cold extraction buffer (Buffer E) containing 25 mM Tris-MES, pH 7.2, 5 mM EGTA, 250 mM sucrose, 1 mM MgSO₄, 0.5% (w/v) BSA, 0.5% (w/v) polyvinylpyrrolidone, 10% (v/v) glycerol, 1 mM phenylmethylsulfonyl fluoride, and 1 mM DTT. Phenylmethylsulfonyl fluoride (PMSF) and DTT were added immediately before use.

The homogenate was filtered through four layers of gauze and centrifuged at 1,500 × *g* for 20 min. The pellet, containing tissue debris, nuclei, unbroken cells, and other dense material, was discarded. The supernatant was centrifuged at 12,000 × *g* for 20 min to obtain a crude organelle-enriched pellet and a corresponding supernatant.

The 12,000 × *g* pellet was resuspended in 1 mL suspension buffer (Buffer S) containing 5 mM potassium phosphate, pH 7.2, 5 mM KCl, 250 mM sucrose, 0.5% (w/v) BSA, and 1 mM freshly added DTT. A portion of the 12,000 × *g* supernatant was centrifuged at 80,000 × *g* for 30 min in a Beckman Coulter ultracentrifuge. The resulting microsome-enriched pellet was resuspended in 1 mL Buffer S.

For H₂ measurements, 500 μg protein from each fraction was incubated in 500 μL Buffer S in a sealed 1.9-mL vial. The vial headspace was flushed with N₂ for 2 min, and samples were incubated for 14 h at 25 °C. H₂ values were corrected against a Buffer S blank and normalized to protein content. Protein concentrations were determined using the Bradford protein assay kit (Tiangen Biotech, Beijing Co., LTD, CN).

### Preparation of crude mitochondrial fractions

Crude mitochondrial fractions were prepared at 4 °C by differential centrifugation, based on Douce et al. (1987), with minor modifications. Hypocotyls (300 g) from 72-96-h-old etiolated mung bean seedlings were homogenized in 900 mL extraction medium containing 0.3 M D-mannitol, 4 mM L-cysteine, 1 mM EDTA-Na₂, 0.1% (w/v) BSA, 0.6% (w/v) polyvinylpyrrolidone, and 30 mM MOPS adjusted to pH 7.5 with NaOH. Homogenization was performed with a Waring blender at low speed for three 5-s bursts.

The homogenate was filtered through four layers of gauze, readjusted to pH 7.5 when necessary, and centrifuged at 1,500 × *g* for 20 min. The pellet was discarded, and the supernatant was centrifuged at 12,000 × *g* for 20 min. The resulting pellet was resuspended in 4 mL washing medium (Buffer W) containing 0.3 M D-mannitol, 0.1% (w/v) BSA, 1 mM EDTA-Na₂, and 10 mM MOPS adjusted to pH 7.2 with NaOH.

The resuspended material was centrifuged at 1,500 × *g* for 10 min, and the resulting supernatant was centrifuged at 12,000 × *g* for 20 min. The final pellet was gently resuspended in a small volume of Buffer W containing or lacking 1 mM EDTA-Na₂, as required for the subsequent experiment. Protein concentration was determined using the Bradford assay. Mitochondrial integrity was assessed by oxygen-electrode measurements of respiratory activity (Jacoby et al., 2015), as described below.

For clarity, the buffers used in the H₂ experiments are designated as follows:

**Buffer E:** 25 mM Tris-MES, pH 7.2, 5 mM EGTA, 250 mM sucrose, 1 mM MgSO₄, 0.5% (w/v) BSA, 0.5% (w/v) polyvinylpyrrolidone, 10% (v/v) glycerol, 1 mM PMSF, and 1 mM DTT.
**Buffer S:** 5 mM potassium phosphate, pH 7.2, 5 mM KCl, 250 mM sucrose, 0.5% (w/v) BSA, and 1 mM DTT.
**Buffer W:** 0.3 M D-mannitol, 0.1% (w/v) BSA, 10 mM MOPS, pH 7.2, with or without 1 mM EDTA-Na₂ as indicated.
**Buffer H:** 0.3 M D-mannitol, 0.1% (w/v) BSA, and 10 mM MOPS, adjusted to pH 6.0 without EDTA. Buffer H is identical in composition to EDTA-free Buffer W except for the pH; it was adopted for the temperature, cross-species, and all subsequent H₂ assays, including those presented in Figure 4, after pH profiling identified pH 6 as optimal (Figure 3I).

### Sealed-vial assays of H₂ accumulation

H₂ accumulation by crude mitochondrial fractions or subcellular fractions was measured in sealed 1.9-mL GC vials with a total liquid volume of 500 μL. Unless otherwise indicated, each vial contained 500 μg protein. No exogenous respiratory substrates were added. For low-O₂ assays, the vial headspace was flushed with N₂ for 2 min before sealing. Air-started controls were sealed under ambient atmosphere without N₂ flushing. For boiled controls, fractions were heated at 100 °C for 5 min before sealing.

Unless otherwise indicated, vials were incubated in darkness at the specified temperature and sampled once at the assay endpoint. Independent vials were prepared for each time point in kinetic experiments. To release dissolved H₂, selected samples were shaken at 400 rpm and 40 °C for 10 min before GC measurement; unshaken samples served as controls. A 1-mL headspace sample was collected for GC–BID analysis. H₂ values from mitochondrial and subcellular-fraction assays were corrected against the corresponding buffer-only controls and normalized to protein content.

For the buffer-additive experiment (Figure 3B), the S (−DTT) formulation, Buffer S without DTT, PMSF, and EDTA, served as the basal medium, to which 1 mM PMSF, 1 mM DTT, or 1 mM EDTA-Na₂ was added separately. For chelator comparison (Figure 3C), crude mitochondrial fractions were incubated in Buffer E (5 mM EGTA), Buffer W (1 mM EDTA-Na₂), or EDTA-free Buffer W.

For the pH-response experiment, EDTA-free Buffer W was adjusted to initial pH values of 4.0-8.0, and samples were incubated at 35 °C for 4 h. For the temperature-response experiment, samples were incubated in Buffer W at 25-45 °C for 4 h.

For cross-species assays, crude mitochondrial fractions prepared from roots of soybean or *A. thaliana* were incubated in Buffer H at 35 °C. Soybean samples were incubated for 6 h, and *A. thaliana* samples were incubated for 6 or 17 h.

### Whole-seedling and excised-organ H₂ assays

For whole-seedling assays, 10 seedlings were placed in a 40-mL headspace vial containing 2 mL sterile distilled water. For organ assays, cotyledons together with true leaves, hypocotyls, or roots were excised from 72-h-old mung bean seedlings. Organs from multiple seedlings were pooled to obtain 4-5 g fresh weight per 22.5-mL vial containing 2 mL sterile distilled water.

For N₂-flushed treatments, vial headspaces were flushed with N₂ for 2 min before sealing. Air-started vials were sealed under ambient atmosphere. Samples were incubated in darkness at 25 °C for 16 h. For the developmental time-course (Figure 2F), seedlings germinated for 0-96 h were collected at the indicated germination times and incubated for an additional 16 h in independently prepared air-started or N₂-flushed vials; each time point comprised an independent set of biological samples (n = 5) rather than repeated measurements of the same vial. Each vial was sampled only once at the endpoint because repeated removal of headspace gas reduced the measured signal. H₂ peak areas were normalized to seedling or organ fresh weight. Unless otherwise stated, data from whole-seedling and organ assays were presented without background subtraction.

### GC–BID analysis of H₂, O₂, and N₂

Headspace gases were analyzed using a Shimadzu GCMS-QP2010 Ultra equipped with a barrier discharge ionization detector and an SH-Rt-Msieve 5A column (30 m × 0.32 mm internal diameter, 30 μm film thickness), following a validated method (Zhang, Z., et al., 2020). Helium was used as the carrier gas at 60 cm s⁻¹. The column and injector temperatures were 50 and 250 °C, respectively; the split ratio was 5:1, the inlet pressure was 200.2 kPa, and the flow rate was 3.85 mL min⁻¹. Headspace samples (1 mL) were injected. Peaks were identified using authentic H₂, O₂, and N₂ standards, with retention times of 1.0 ± 0.1, 1.4 ± 0.1, and 2.1 ± 0.1 min, respectively. Headspace O₂ was estimated from integrated peak areas relative to an air-started control (21% O₂) and was not interpreted as dissolved O₂.

H₂ was quantified using external standards prepared in identical vials with the same buffer and equilibrated under the same conditions as the samples. Calibration models used integrated peak area as the predictor and H₂ amount per vial as the response (Figure 2—figure supplement 1A–D). Linear calibration curves (R² ≥ 0.995) covered low (0-50), middle (0-400), and high (0-2449 nmol vial⁻¹) ranges. Low- and middle-range standards were prepared by diluting saturated hydrogen-rich water (800 μmol L⁻¹ at 25 °C and 1 atm) (Wilhelm, Battino, and Wilcock 1977). Additional middle- and high-range standards were prepared by injecting 2.5-10 and 10-60 μL, respectively, of pure H₂ gas (99.999%). Gas volumes were converted to amounts using the ideal-gas molar volume at 25 °C (24.47 L mol⁻¹). Samples within overlapping ranges were quantified using the higher-range curve.

The limit of detection (S/N = 3) was 0.1 nmol H₂ vial⁻¹, and the limit of quantification (S/N = 10) was 0.3 nmol H₂ vial⁻¹. Values below the LOD were classified as non-detectable and replaced with half the detection limit (0.05 nmol vial⁻¹).

Using identical vial configurations and equilibration conditions allowed the calibration to account for gas–liquid partitioning (Wilhelm et al. 1977; Zhang, Z., et al., 2020); the gas-to-liquid volume ratio exceeded 2, minimizing the fraction of H₂ remaining dissolved. Cross-validation of the middle-range HRW- and gas-derived curves showed that eight of nine back-calculated values were within ±10% of their nominal amounts (mean signed deviation, +0.2%). Bland–Altman analysis gave a mean bias of +0.19 nmol vial⁻¹ and 95% limits of agreement of −45.10 to +45.48 nmol vial⁻¹ (Figure 2—figure supplement 1E, F).

### Oxygen-electrode measurements

Oxygen consumption was measured at 25 °C using a Clark-type oxygen electrode (Hansatech Oxytherm, Hansatech Instruments Ltd., UK) in a total reaction volume of 500 μL (Jacoby et al., 2015). Freshly isolated crude mitochondrial fractions equivalent to 500 μg protein were analyzed in EDTA-free Buffer W at pH 7.2. NADH and ADP were added at the indicated times to final concentrations of 1 mM and 200 μM, respectively.

For measurements after H₂ accumulation, crude mitochondrial fractions were first incubated for 6 h at 25 °C in EDTA-free Buffer W under the standard sealed-vial conditions. The reaction mixture was then transferred directly to the electrode chamber without washing, centrifugation, or resuspension. After the 6-h incubation, the pH of the reaction mixture was measured with the built-in pH electrode of the Oxytherm electrode chamber (final pH 6.46). Oxygen-consumption rates were calculated from the linear portions of the electrode traces and are expressed as μmol O₂ L⁻¹ min⁻¹.

### Changes in pH during H₂ accumulation

For each independent mitochondrial preparation, parallel reaction aliquots were incubated separately at 35 °C for 1, 2, or 3 h under the conditions described in Buffer H, except that the initial medium pH was 6.5. At the designated endpoint, each aliquot was transferred to the small-volume chamber of a Hansatech Oxytherm liquid-phase oxygen-electrode system, and its pH was measured once using a calibrated pH sensor. The Oxytherm chamber was used to permit accurate measurement of the small reaction volume; pH was measured after incubation rather than monitored continuously during the reaction.

### Effects of oxygen on mitochondrial H₂ accumulation

The mitochondrial H₂ evolution assay was performed as described in the previous section. To establish hypoxia, the headspace of reaction vials was flushed with nitrogen gas (N₂) for 2 min; control samples were maintained under ambient atmospheric oxygen conditions. All samples were incubated in darkness at either 25 °C for 2, 4, 6, 8, 10, and 12 h, or at 35 °C for 1, 2, 3, 4, 5, 6, 7, and 8 h. H₂ accumulation was quantified using headspace gas chromatography under the conditions detailed earlier.

### Effect of mitochondrial electron transport inhibitors on H₂ accumulation

To assess the contribution of specific respiratory chain complexes to H₂ evolution, inhibitors targeting different steps of oxidative phosphorylation were applied. The inhibitors included rotenone (complex I), malonic acid (complex II), antimycin A (complex III), sodium azide (NaN₃, complex IV), oligomycin (ATP synthase), SHAM (alternative oxidase, AOX) and carbonyl cyanide p-trifluoromethoxyphenylhydrazone (FCCP, an uncoupler).

Rotenone, antimycin A, oligomycin, SHAM and FCCP were dissolved in dimethyl sulfoxide (DMSO); malonic acid and NaN₃ were prepared directly in H₂ evolution buffer (Buffer H). Reaction mixtures (500 µL final volume in Buffer H) contained 300-500 µg crude mitochondria and one of the following inhibitors: 5 µM rotenone, 10 mM malonic acid, 2 mM SHAM, 10 µM antimycin A, 5 mM NaN₃, 2 µM oligomycin, or 25 µM FCCP. Control samples contained mitochondria without inhibitor but with an equal volume of solvent (DMSO or Buffer H). Hypoxic conditions were established as described previously. After 3 h of incubation at 35 °C in darkness, H₂ accumulation was quantified by headspace gas chromatography.

### Effect of NADH on H₂ accumulation

To test whether exogenous NADH can directly support H₂ production, crude mitochondria (300-500 μg) were incubated in Buffer H (pH 6.0, 500 μL final volume) with or without 1 mM NADH. Hypoxic conditions were established by flushing the headspace with N₂ for 2 min. After 3 h of incubation at 35 °C in darkness, H₂ accumulation was quantified by headspace GC-BID as described above.

### Effect of metabolic intermediates and ubiquinone on mitochondrial H₂ accumulation

To investigate the role of mitochondrial electron transport chain components in H₂ evolution, various substrates and inhibitors were applied. Malate and pyruvate were used to supply NADH to complex I; succinate and fumarate served as substrate and product of complex II, respectively. Rotenone was used to inhibit complex I by blocking electron transfer from NADH to ubiquinone. Malonic acid, a competitive inhibitor of complex II, was applied to suppress succinate dehydrogenase activity.

For substrate experiments, 500 µg of crude mitochondria was incubated in Buffer H (pH 6.0, 500 µL final volume) with either malate (5 mM) plus pyruvate (5 mM), or succinate (10 mM), fumarate (5 mM), or succinate plus fumarate (10 mM and 5 mM), in the absence or presence of rotenone (5 µM) or malonate (10 mM). Control reactions contained mitochondria without added substrate or inhibitor.

For ubiquinone competition experiments, mitochondria (500 µg) were incubated in Buffer H (pH 6.0, 500 µL) with mixed substrates (MS; 5 mM malate, 5 mM pyruvate, 10 mM succinate, 5 mM fumarate) in the absence or presence of oxidized ubiquinone-10 (UQ; 60 µM), UQ + rotenone (5 µM), or UQ + malonate (10 mM).

Hypoxic conditions were established by flushing the headspace with N₂ as previously described. After 3 h of incubation at 35 °C in darkness, H₂ accumulation was quantified by headspace gas chromatography.

### Measurement of NADH content during H₂ accumulation

NADH levels were quantified colorimetrically using the Amplite™ Colorimetric NADH Assay Kit (AAT Bioquest) according to the manufacturer’s instructions. This assay employs a chromogenic NADH sensor with maximum absorbance at 460 nm upon reduction. Aliquots (50 µL) of the mitochondrial H₂ evolution reaction mixture were collected at intervals over 7 h and mixed with 50 µL of working solution. After incubation in the dark at room temperature for 1 h, absorbance was measured at 460 nm using a microplate reader. A standard curve was generated using known concentrations of NADH. Because crude mitochondrial suspensions themselves absorb and scatter light at 460 nm, the reading of the 0-h aliquot, measured immediately after mitochondrial preparation, was subtracted from all subsequent readings as the background reference; NADH concentrations are therefore reported as nmol mg⁻¹ protein relative to time zero.

### Malate synthase activity assay

MLS activity was measured at 232 nm as the acetyl-CoA-dependent decrease in absorbance (ε₂₃₂ = 4.5 × 10³ M⁻¹ cm⁻¹) in 100 μL containing 100 mM Tris-HCl (pH 8.0), 5 mM MgCl₂, 1 mM glyoxylate, 0.15 mM acetyl-CoA and 10 μg crude mitochondrial protein. Acetyl-CoA-free and protein-free reactions served as controls. A₂₃₂ was read over 12 min at 25 °C; activity was calculated from the initial rate and expressed as apparent specific activity (nmol min⁻¹ mg⁻¹ protein).

### Analysis of mitochondrial metabolites

Isolated mitochondria (500 µg) were aliquoted into 1.9 mL sealed vials, and the reaction volume was adjusted to 500 µL with Buffer H. The headspace was flushed with N₂ for 2 min to remove O₂. Samples were then incubated under normoxic or hypoxic conditions at 35 °C for 0-7 h. After incubation, samples were immediately flash-frozen in liquid nitrogen and stored at −80 °C until analysis. Detailed MS analytical conditions are described below. Metabolites were extracted as follows. Thawed samples were vortexed thoroughly, and 200 µL of each sample was added to 800 µL of a pre-cooled 1:1 (v/v) methanol/acetonitrile solution (Merck). After vigorous vortexing, the mixture was sonicated for 20 min in an ice bath and incubated at −20 °C for 1 h to precipitate proteins. The sample was then centrifuged at 14,000×g for 20 min at 4 °C. The supernatant was collected and dried under vacuum. The resulting residue was reconstituted in 100 µL of acetonitrile/water (1:1, v/v), followed by centrifugation at 14,000×g for 20 min at 4 °C. The final supernatant was transferred to LC vials for subsequent analysis.

No internal standard was added. All samples were processed identically, and metabolite levels are reported as relative abundances (peak areas normalized to the initial time point or total ion current). The extraction and LC/MS analysis were performed under identical conditions for all time points, allowing reliable comparison of temporal trends. Absolute quantification was not performed, but this does not affect the relative comparisons across the time course.

### HPLC–MS analysis of mitochondria treated at 25 °C

The samples were separated by liquid chromatography (LC) using Agilent 1290 Infinity LC HPLC (Agilent). ACQUITY UPLC BEH amide column (1.7μm, 2.1 mm×100 mm) from Waters was used. Mobile phase. A: 10 mM ammonium acetate aqueous solution. B: acetonitrile. Samples were placed in the automatic sampler at 4 °C. Injection volume: 6 µL. Column temperature: 45 °C. Flow rate: 300 µL /min. Gradient of the related liquid phase: 0-18 min 90% to 40% B, 18-18.1 min, 40% to 90% B, 18.1-23 min, 90% B. The mixture of standard metabolites was set up in the sample queue for correction of chromatographic retention time. A 5500 QTRAP MS (AB SCIEX) was used for MS analysis under negative ion mode. 5500 QTRAP ESI source conditions were as follows: source temperature: 450 °C, ion Source Gas1(Gas1): 45, ion Source Gas2 (Gas2): 45, Curtain gas (CUR): 30, ionSpray Voltage Floating (ISVF): -4500 V. Peak areas were extracted with MultiQuant and identified against authentic standards.

### Targeted metabolite analysis at 35 °C

Metabolic changes in samples incubated at 35 °C were analyzed using a Thermo Orbitrap UHPLC–MS system. Chromatographic separation was performed with a mobile phase consisting of acetonitrile and water (80:20, v/v) at a flow rate of 200 µL/min; the injection volume was 10 µL. Mass spectrometric detection was carried out on a Thermo LTQ Orbitrap XL MS operating in negative ionization mode. Key MS parameters were set as follows: capillary temperature, 320 °C; resolution, 30,000; AGC target, 2 × 10⁵; maximum injection time, 500 ms. The instrument was calibrated before analysis using the manufacturer’s standard calibration solution.

For metabolite identification, MS/MS analysis was performed using collision-induced dissociation (CID). Product ion spectra were compared against both the Thermo compound database and the mzCloud advanced mass spectral database (https://www.mzcloud.org) to match characteristic fragment ions for metabolite annotation. For targeted quantification of key organic acids, the following precursor ions and collision energies were applied: m/z 145.01 (α-ketoglutarate, 22 eV), m/z 117.01 (succinate, 30 eV), m/z 115.00 (fumarate, 25 eV), and m/z 133.01 (malate, 26 eV). Representative MS/MS spectra are shown in Figure 5—figure supplement 2.

### Statistical analysis

Data analysis and graph generation were performed using GraphPad Prism version 9.0 (GraphPad Software, USA). Proposed mechanistic diagrams were prepared with ChemDraw Professional (PerkinElmer, USA).

The replicate structure and exact statistical test used for each experiment are specified in the corresponding figure legend. Unless otherwise indicated, pairwise comparisons were evaluated using a two-tailed unpaired Student’s *t*-test, and multiple comparisons were analyzed by ordinary one-way ANOVA followed by Dunnett’s post hoc test. Where variances were unequal or multiple comparisons were not against a single control, alternative tests (Welch ANOVA, Šídák) were used as specified in the figure legends. Data are presented as mean ± SD. Significance levels: \**P* < 0.05, \*\**P < 0.01, ***P < 0.001,* \*\*\*\**P* < 0.0001.

### Theoretical calculations

All thermodynamic and kinetic calculations were performed as described in Supplementary Calculations S1–S13. Key parameters and equations included: Standard redox potentials (Supplementary Calculation S1); Nernst corrections for low H₂ partial pressure and high FMNH₂/FMN ratio (S2); Crossed-potential estimation for CI-bound FMN (S3); Electron bifurcation energetics (S4); H₂ formation step thermodynamics (S5); Overall energy balance per NADH (S6); Quantitative resolution of the rotenone paradox (S7); Moser–Dutton electron transfer rates using CI* distances from Maldonado et al. (2020) (S8); Sensitivity analysis (S9); Effect of pH on H₂ formation (S10); The Moser–Dutton ruler was applied with a reorganization energy λ = 0.7 eV (Moser & Dutton, 1992).

## Acknowledgements

We thank Yanan Wei and Muhan Li for their assistance with plant seedling cultivation and mitochondrial extraction. We also thank Yao Mawulikplimi Adzavon and Xiaokang Zhang for assisting with experiments and data analysis, as well as Andong Wang, Jihong Sun, Yunlong Shao, and Xiayan Wang for their support with gas chromatography and UHPLC-HRMS experiments.

## Author contributions

M.X. conceptualized the study and proposed the hypothesis; Z.X., Z.Z., L.M., and X.Z. developed the methodology; Z.X., Z.Z., L.M., X.Z., Z.P., and X.F. performed the formal analysis and investigation; M.X. performed the theoretical calculations; M.X., Z.X., and Z.Z. wrote the original draft; M.X. reviewed and edited the manuscript; Z.X., Z.Z., L.Y., and M.X. visualized the data; M.X. supervised the project and managed project administration; M.X. and Z.X. acquired the research funding.

## Data availability

All data and materials supporting the conclusions of this study are included in the article or provided as supplemental information.

## Code availability

No custom code was used in this study.

## Artificial Intelligence Use Statement

During the preparation of this work, the authors used Claude (Anthropic) and DeepSeek for language editing. Nano Banana2 assisted in designing Figure 1A. All AI-assisted content was reviewed and edited by the authors, who take full responsibility for the final publication.

## Supplementary Information

Detailed thermodynamic and kinetic calculations are provided in Supplementary Calculations S1–S13. Additional supporting data, including GC calibration curves, subcellular fractionation, exploratory metabolic profiling, MS/MS spectra, partial correlation analysis, and raw data summaries (mean ± SD) for inhibitor and substrate experiments, are presented in the figure supplements to Figures 2, 3, and 5 and Supplementary Tables S1–S4.

## Supplementary Information

### 1. Supplementary Figures

**Figure 2—figure supplement 1.**
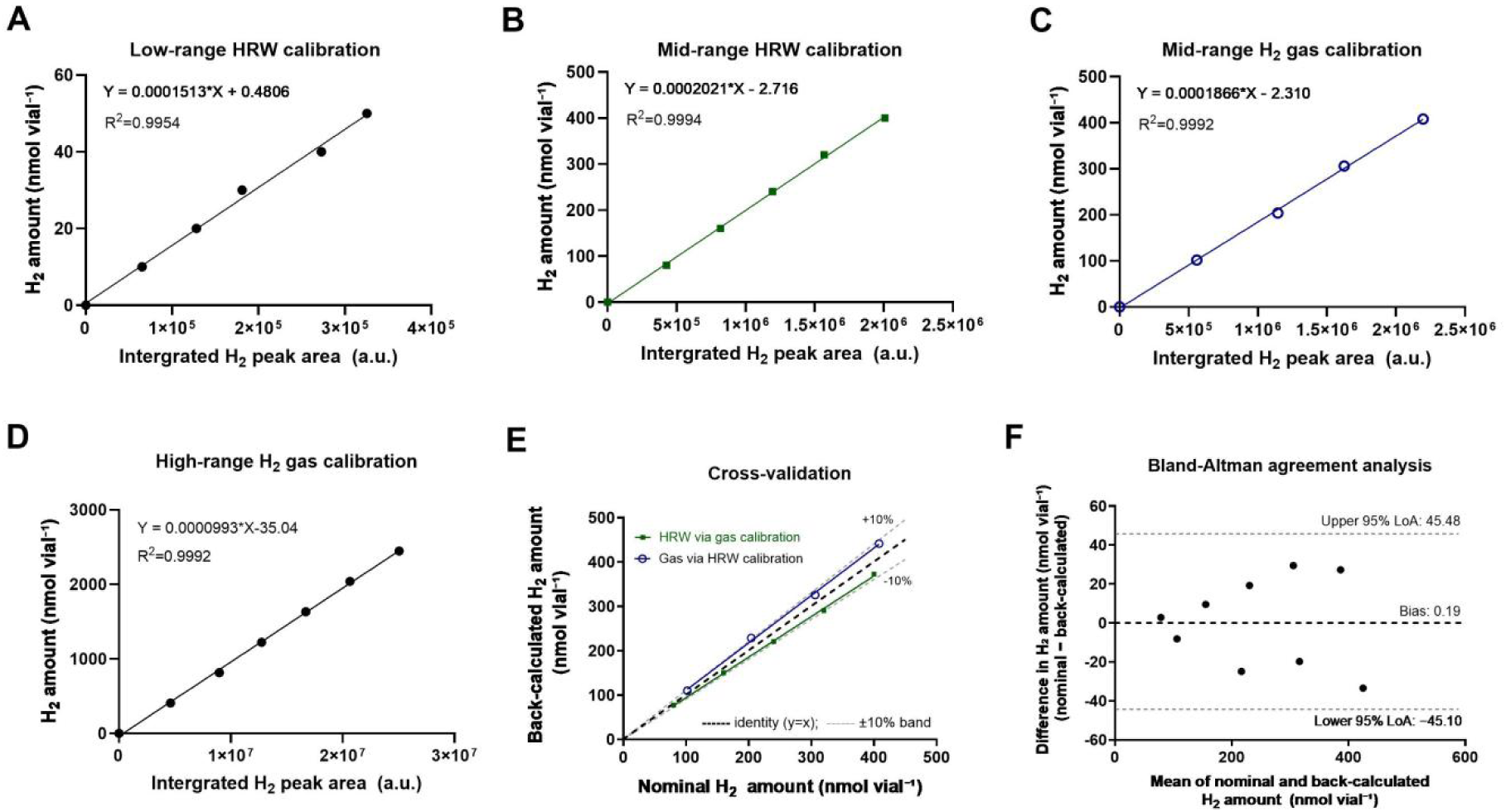
H₂ calibration curves and cross-validation of standard-preparation methods. **(A–D)** Calibration curves generated using standards prepared from saturated hydrogen-rich water (HRW) or pure H₂ gas (99.999%). H₂ amount was plotted against integrated H₂ peak area for **(A)** low-range HRW (0–50 nmol vial⁻¹), **(B)** middle-range HRW (0–400 nmol vial⁻¹), **(C)** middle-range gas (0–400 nmol vial⁻¹; 2.5–10 μL injected), and **(D)** high-range gas standards (0–2449 nmol vial⁻¹; 10–60 μL injected). Gas volumes were converted to amounts using the ideal-gas molar volume at 25 °C (24.47 L mol⁻¹). Each point represents the mean of three replicate injections. Solid lines show linear fits, with equations and (R^2^) values indicated. **(E)** Cross-validation of the two middle-range calibration curves. HRW standards were back-calculated using the gas-derived curve in panel C, and gas standards using the HRW-derived curve in panel B. Dashed lines indicate identity (y=x) and ±10% deviation. Eight of nine values were within ±10% of nominal amounts, with a mean signed deviation of +0.2%. **(F)** Bland–Altman analysis of nominal and back-calculated H₂ amounts. Differences (nominal minus back-calculated) are plotted against their means. The mean bias was +0.19 nmol vial⁻¹, with 95% limits of agreement of −45.10 and +45.48 nmol vial⁻¹. The limits of detection and quantification were 0.1 and 0.3 nmol H₂ vial⁻¹, respectively, at signal-to-noise ratios of 3 and 10. Calibration, cross-validation, and quantification procedures are described in Methods.

**Figure 3—figure supplement 1.**
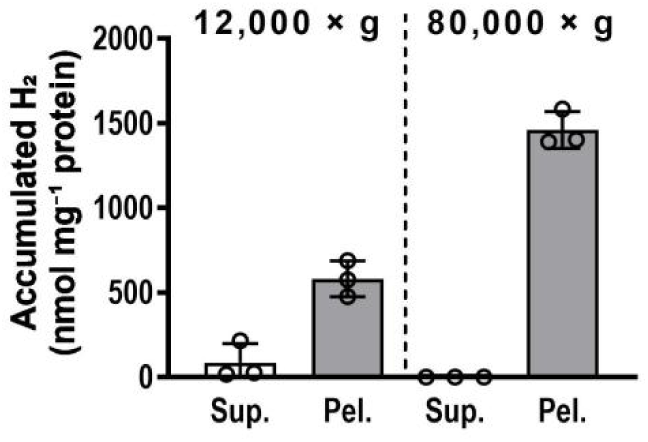
H₂-accumulating activity is enriched in sedimentable fractions of *Vigna radiata* hypocotyl homogenates. Hypocotyl homogenates from 72–96-h-old etiolated *Vigna radiata* seedlings were clarified at 1,500 × *g*. The resulting supernatant was centrifuged at 12,000 × *g* for 20 min to generate a crude organelle-enriched pellet and the corresponding supernatant. A portion of the 12,000 × *g* supernatant was subsequently centrifuged at 80,000 × *g* for 30 min to obtain a microsome-enriched pellet and the corresponding final supernatant. Pellets were resuspended in Buffer S. Each H₂ assay contained 500 μg protein in 500 μL Buffer S in a sealed 1.9-mL vial. Vial headspaces were flushed with N₂ for 2 min, and samples were incubated for 14 h at 25°C. H₂ values were corrected against the Buffer S blank and normalized to protein content. Bars show mean ± s.d.; circles represent individual assay measurements (n=3)(n=3). No inferential statistical test was performed. Sup., supernatant; Pel., pellet.

**Figure 5—figure supplement 1.**
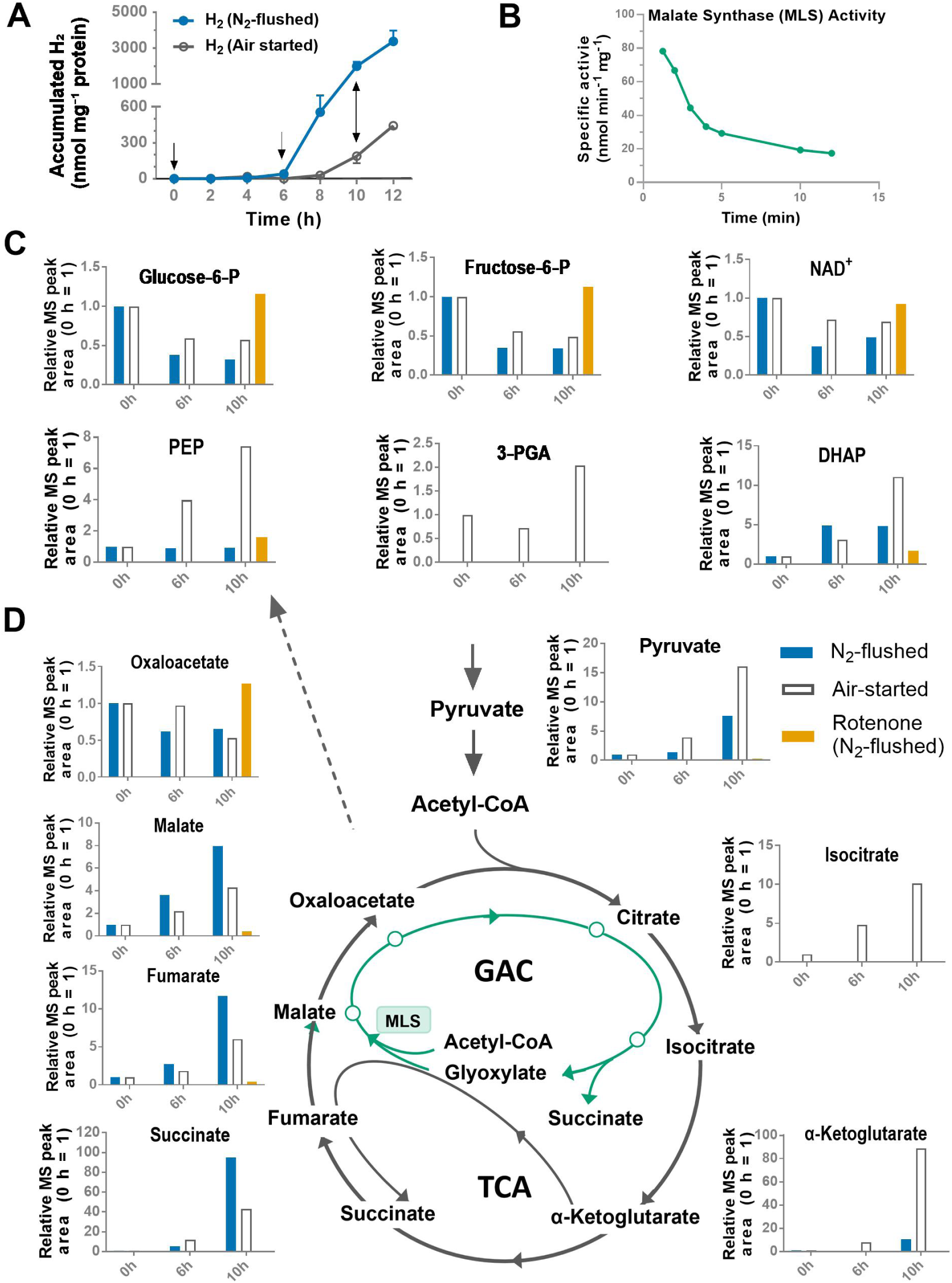
Exploratory metabolic profiling of the crude mitochondrial fraction under N₂-flushed, air-started, and rotenone-treated conditions. A crude mitochondrial fraction (500 μg protein) isolated from *Vigna radiata* hypocotyls was incubated in Buffer H (500 μL) at 25 °C under N₂-flushed or air-started conditions. A separate N₂-flushed sample containing rotenone (5 μM) was analyzed at 10 h only. Metabolomic samples were collected at 0, 6, and 10 h; the common 0-h sample was frozen immediately after preparation and used for normalization. Each metabolomic condition and time point was analyzed once (*n* = 1). The dataset therefore represents an exploratory screen that identified condition-dependent changes in TCA-cycle-associated metabolites and motivated the subsequent time-resolved experiment at 35 °C (Figure 5). Peak areas can be compared across conditions and time points within each metabolite, but not between different metabolites. **(A)** Accumulated headspace H₂ under N₂-flushed and air-started conditions. Arrows indicate metabolomic sampling at 0, 6, and 10 h. H₂ accumulation was more rapid under N₂-flushed conditions, reaching ∼3,000 nmol mg⁻¹ protein at 12 h. Data are replotted from Figure 3E. **(B)** Malate synthase (MLS) activity detected in the crude organellar fraction (containing mitochondria and glyoxysomes). Specific activity was monitored over 12 min. The detectable activity indicates that glyoxylate-cycle-associated enzymatic capacity was retained in the preparation. **(C)** Relative abundances of glycolytic intermediates and NAD⁺, normalized to the common 0-h value. The six-carbon intermediates glucose-6-phosphate and fructose-6-phosphate differed only modestly between the N₂-flushed and air-started conditions. In contrast, the three-carbon intermediates PEP, 3-PGA, and DHAP showed larger condition-dependent changes and accumulated preferentially in the air-started samples. At 10 h, rotenone treatment was associated with higher glucose-6-phosphate, fructose-6-phosphate, and NAD⁺, but lower levels of the three-carbon intermediates than in the air-started sample, suggesting that complex I inhibition might have stalled glycolytic carbon flux downstream of the hexose-phosphate pool. **(D)** Relative abundances of TCA-cycle- and glyoxylate-cycle-associated metabolites. Under N₂-flushed conditions, succinate showed the largest increase (∼95-fold at 10 h), accompanied by fumarate and malate accumulation, consistent with operation of the reductive branch of the TCA cycle and glyoxylate-cycle-associated carbon input. In contrast, air-started samples displayed a distinct metabolic signature dominated by pyruvate, isocitrate, and α-ketoglutarate (∼16-, ∼10-, and ∼89-fold at 10 h, respectively), reflecting accumulation at the oxidative (NAD⁺-dependent) decarboxylation steps of the TCA cycle. Rotenone treatment strongly attenuated the 10-h accumulation of pyruvate, malate, fumarate, succinate, and α-ketoglutarate, consistent with inhibition of complex I-linked redox cycling and TCA-cycle turnover. Gray and orange lines denote the TCA and glyoxylate cycles, respectively. Gray and green lines denote the TCA and glyoxylate cycles, respectively. Green filled bars, open bars, and yellow filled bars indicate N₂-flushed, air-started, and rotenone-treated N₂-flushed samples, respectively. Metabolite abundances are expressed as relative MS peak areas in arbitrary units.

**Figure 5—figure supplement 2.**
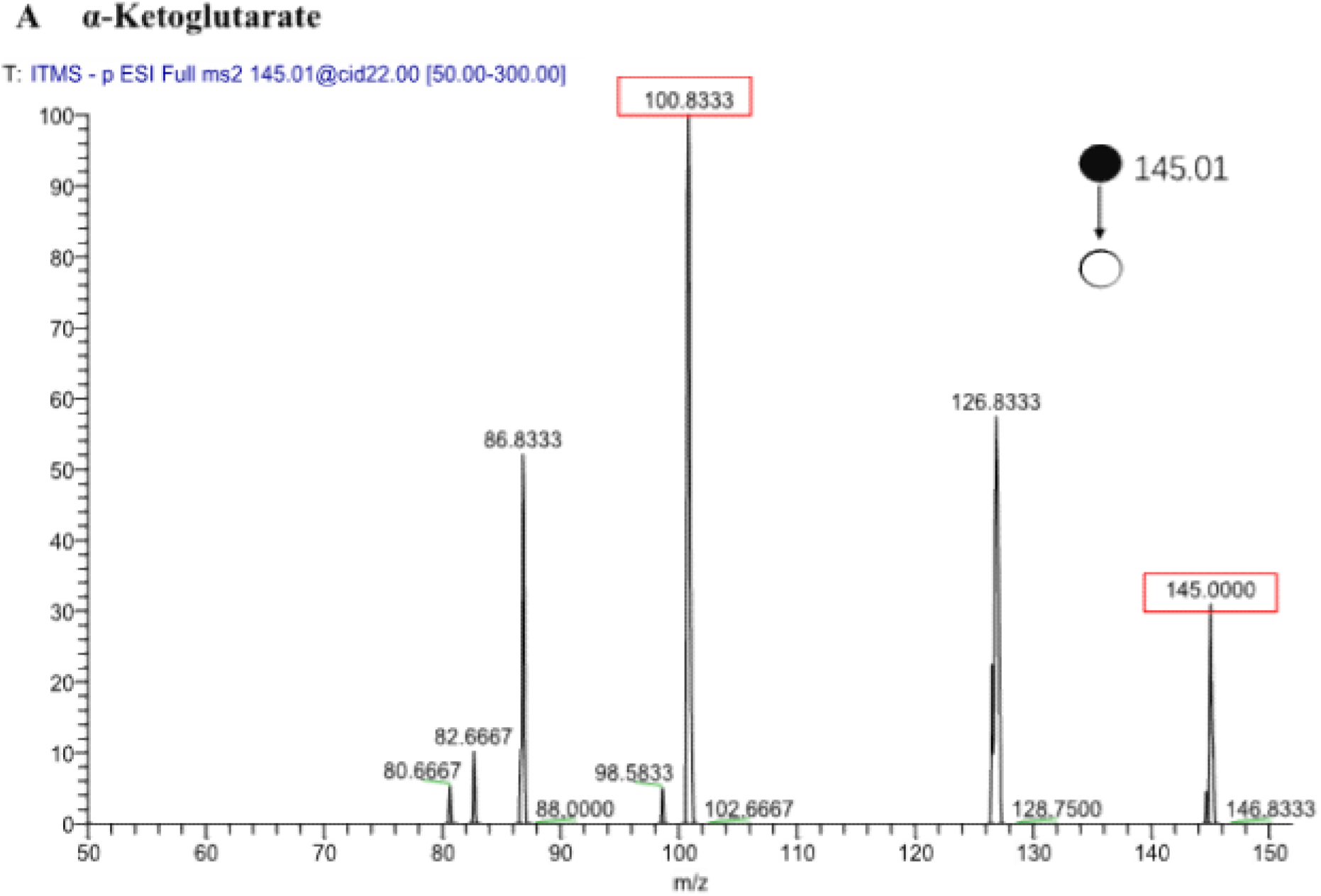
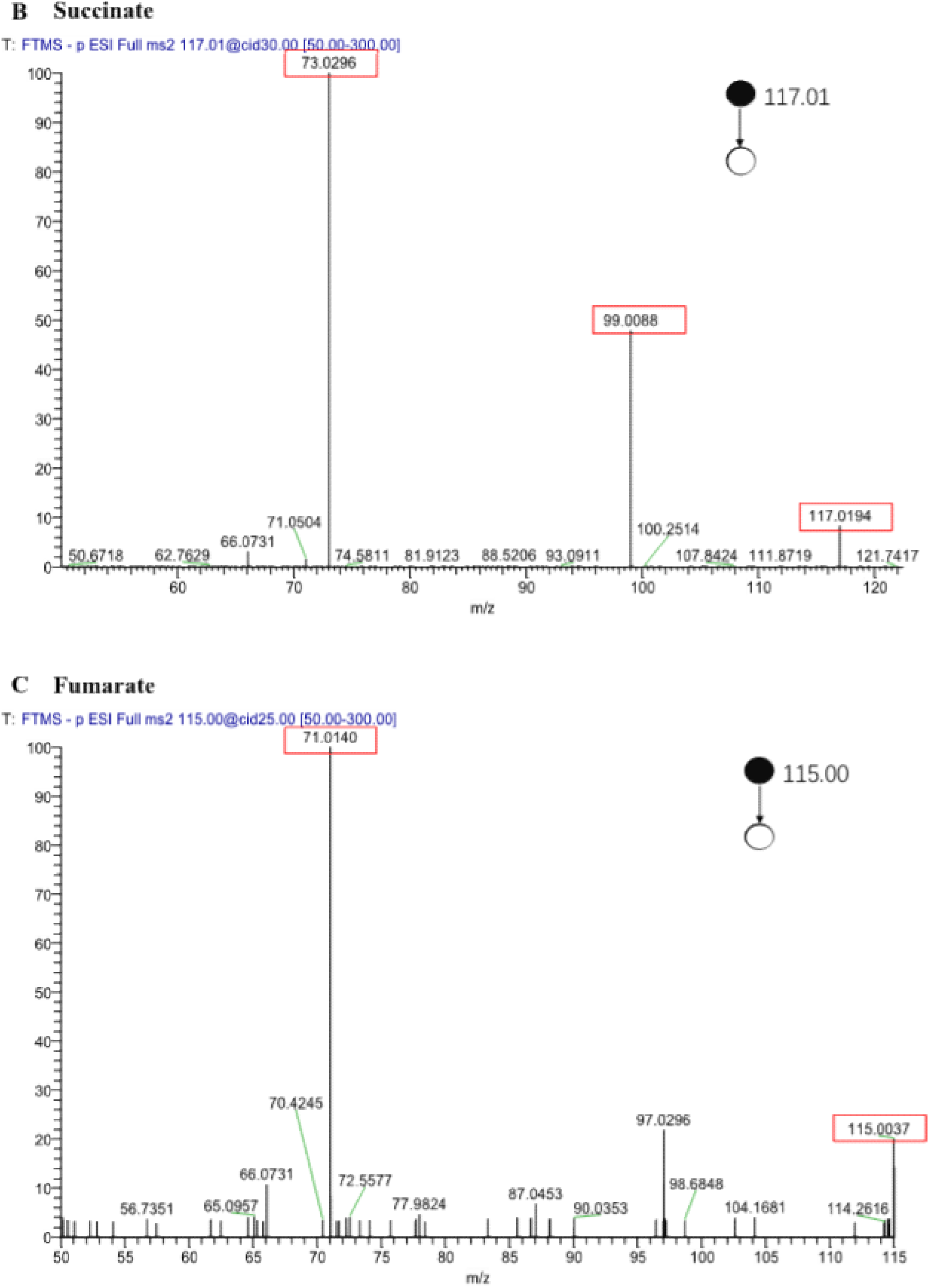
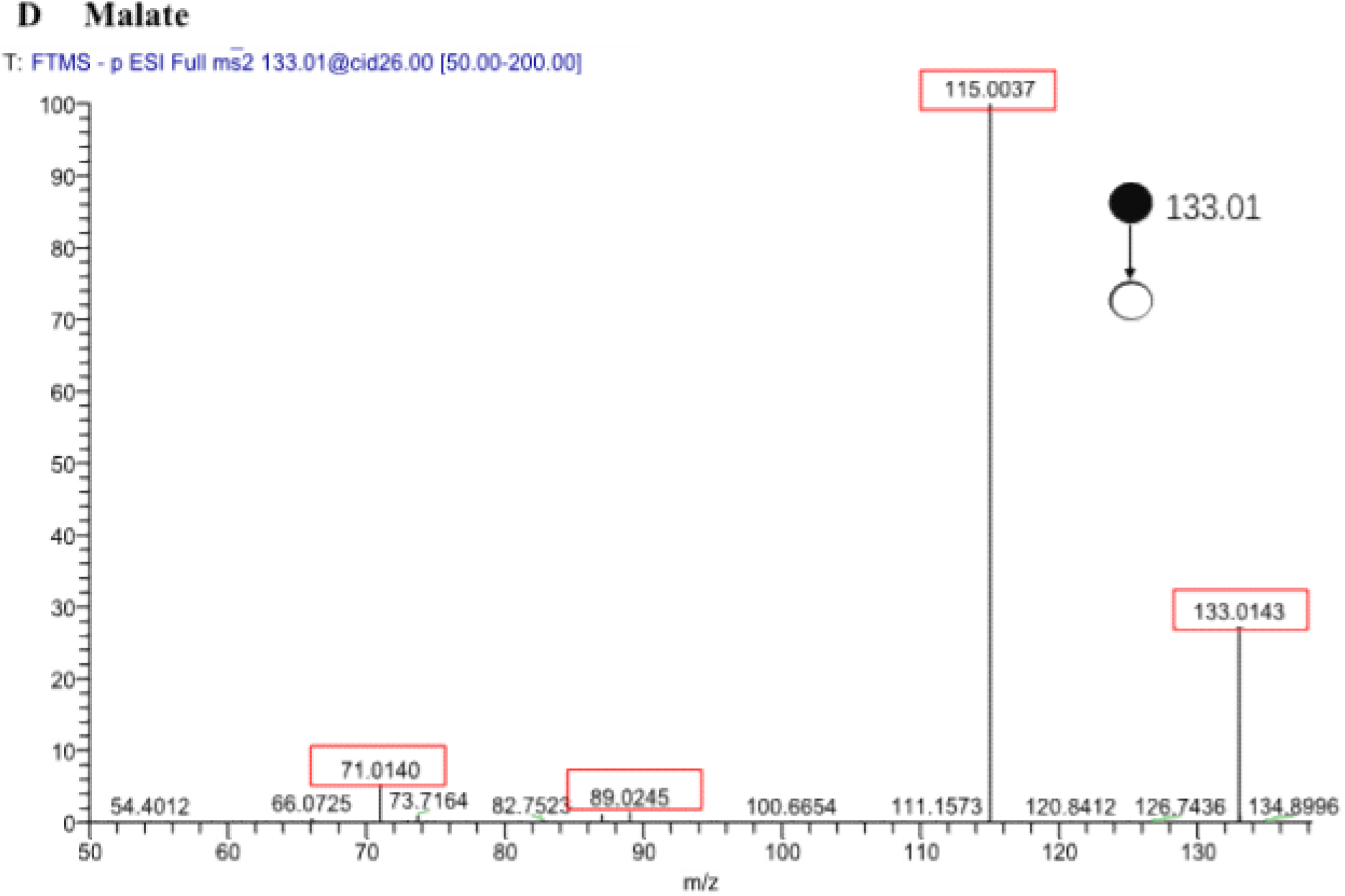
Representative MS/MS spectra of TCA cycle intermediates. Tandem mass spectra (collision-induced dissociation, negative ion mode) of the following metabolites, identified by matching against the mzCloud database: (A) α-Ketoglutarate (precursor m/z = 145.01, collision energy = 22 eV). (B) Succinate (precursor m/z = 117.01, CE = 30 eV). (C) Fumarate (precursor m/z = 115.00, CE = 25 eV). (D) Malate (precursor m/z = 133.01, CE = 26 eV)

Figure 5—figure supplement 2. The secondary MS of mitochondrial metabolites.

### 2. Supplementary Tables

**Supplementary Table S1.** Summary of inhibitor effects on mitochondrial H₂ accumulation.

| Treatment | Concentration | Relative H<sub>2</sub> production (mean ± SD, n=3) | Inhibition (%) | Prediction tested |
|---|---|---|---|---|
| Control (Mit only) | — | 1.000 ± 0.000 | — | — |
| Rotenone (CI) | 5 μM | 0.0132 ± 0.0102 | 98.7 ± 1.0 | P1 |
| Malonate (CII) | 10 mM | 0.815 ± 0.070 | 18.5 ± 7.0 | P6 |
| Antimycin A (CIII) | 10 μM | 0.619 ± 0.113 | 38.1 ± 11.3 | P4 |
| Sodium azide (CIV) | 5 mM | below LOD | not quantifiable | P2 |
| Oligomycin (ATP synthase) | 2 μM | 1.348 ± 0.472 | —<br>(enhancement) | P2 |
| SHAM (AOX) | 2 mM | below LOD | not quantifiable | P3 |
| FCCP (uncoupler) | 25 μM | below LOD | not quantifiable | P2 |

**Note:**
● Relative H<sub>2</sub> production is normalized to the control (Mit only). Data are mean ± SD from three independent biological replicates.
● Inhibition (%) = (1 – Relative H<sub>2</sub>) × 100. For Oligomycin, the value >1 indicates enhanced H<sub>2</sub> production (135% of control).
● Predictions refer to Table 1 in the main text.
● The small negative values (e.g., for NaN<sub>3</sub> and SHAM) were treated as zero for inhibition calculation; the reported values are based on raw data.

**Supplementary Table S2.** Summary of substrate effects on mitochondrial H₂ accumulation.

| **Treatment** | **Concentration** | **Relative H<sub>2</sub> production (mean ± SD, n=3)** | **Change (fold / % inhibition)** | **Prediction tested** |
|---|---|---|---|---|
| **A. Path 1 (Direct NADH)** |  |  |  |  |
| Control (Mit only) | — | 1.000 ± 0.000 | — | — |
| Malate + Pyruvate (M+P) | 5 mM each | 10.7 ± 1.8 | 10.7-fold | P7 |
| M+P + Rotenone | 5 µM | 3.3 ± 0.7 | 68.7% inhibition | — |
| **B. Path 2 (Succinate-driven RET)** |  |  |  |  |
| Succinate | 10 mM | 11.1 ± 3.6 | 11.1-fold | — |
| Fumarate | 5 mM | 9.3 ± 1.8 | 9.3-fold | — |
| Succinate + Fumarate (S+F) | 10 + 5 mM | 11.7 ± 1.2 | 11.7-fold | P6/P7 |
| S+F + Malonate | 10 mM | 3.3 ± 0.8 | 71.3 ± 7.0% inhibition | P6 |
| **C. Ubiquinone competition** |  |  |  |  |
| Control (Mit only) | — | 1.000 ± 0.000 | — | — |
| UQ (60 µM) | 60 µM | 0.069 ± 0.051 | 93.1% inhibition | — |
| Mixed substrates (MS)<sup>1</sup> | 5 mM each | 20.6 ± 3.6 | 20.6-fold | P7 |
| MS + UQ | 60 µM | 6.0 ± 0.5 | 70.7% inhibition | — |
| MS + UQ + Rotenone | 60 µM + 5 µM | 0.135 ± 0.057 | 97.8 ± 2.3% inhibition | P1/P7 |
| MS + UQ + Malonate | 60 µM + 10 mM | 4.08 ± 0.38 | 32.4 ± 2.5% inhibition | P6 |

<sup>1</sup>Mixed substrates (MS) = 5 mM malate, 5 mM pyruvate, 10 mM succinate, 5 mM fumarate.

**Supplementary Table S3. Time-course analysis of H_2_ production, NADH, and TCA-cycle intermediates**

**Table S3-1.**
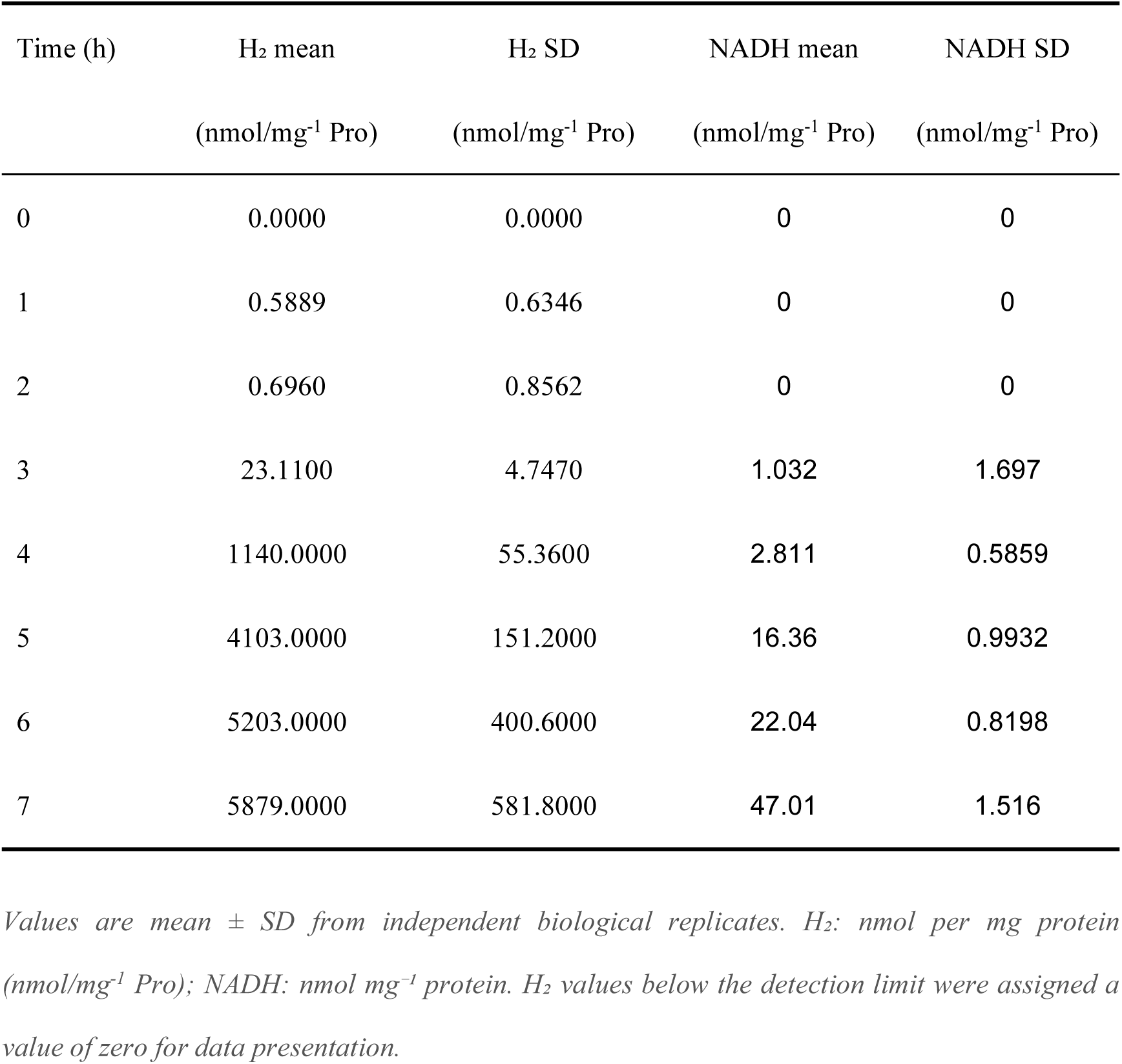
Time-course measurements of H₂ production and NADH concentration.

| Time (h) | H<sub>2</sub> mean<br>(nmol/mg<sup>-1</sup> Pro) | H<sub>2</sub> SD<br>(nmol/mg<sup>-1</sup> Pro) | NADH mean<br>(nmol/mg<sup>-1</sup> Pro) | NADH SD<br>(nmol/mg<sup>-1</sup> Pro) |
|---|---|---|---|---|
| 0 | 0.0000 | 0.0000 | 0 | 0 |
| 1 | 0.5889 | 0.6346 | 0 | 0 |
| 2 | 0.6960 | 0.8562 | 0 | 0 |
| 3 | 23.1100 | 4.7470 | 1.032 | 1.697 |
| 4 | 1140.0000 | 55.3600 | 2.811 | 0.5859 |
| 5 | 4103.0000 | 151.2000 | 16.36 | 0.9932 |
| 6 | 5203.0000 | 400.6000 | 22.04 | 0.8198 |
| 7 | 5879.0000 | 581.8000 | 47.01 | 1.516 |

*Values are mean ± SD from independent biological replicates. H<sub>2</sub>: nmol per mg protein (nmol/mg<sup>-1</sup> Pro); NADH: nmol mg<sup>−1</sup> protein. H<sub>2</sub> values below the detection limit were assigned a value of zero for data presentation.*

**Table S3-2.** Mass-spectrometric peak areas of TCA-cycle intermediates during the hypoxic time course.

| Sample | Succinate Mean | Succinate SD | α-KG Mean | α-KG SD | Malate Mean | Malate SD | Fumarate Mean | Fumarate SD |
|---|---|---|---|---|---|---|---|---|
| B | 1,931,598 | 1,724,535 | 30,090 | 11,399 | 706,065 | 456,210 | 50,524 | 50,592 |
| 0 h | 7,298,694 | 514,434 | 1,204,214 | 1,862,316 | 46,356,303 | 26,736,533 | 8,569,850 | 1,336,497 |
| 1 h | 1,838,875 | 1,473,577 | 510,860 | 480,736 | 16,118,201 | 9,617,573 | 5,378,417 | 4,546,787 |
| 2 h | 2,088,968 | 2,228,466 | 1,289,528 | 1,470,369 | 37,615,338 | 18,552,506 | 5,940,326 | 1,163,536 |
| 3 h | 4,150,569 | 3,049,960 | 1,200,516 | 1,217,514 | 47,640,014 | 37,289,855 | 13,256,818 | 6,890,940 |
| 4 h | 72,617,030 | 51,613,527 | 223,973 | 152,741 | 23,231,986 | 16,120,697 | 8,710,499 | 3,626,195 |
| 5 h | 128,315,914 | 62,366,568 | 2,401,762 | 1,363,971 | 18,905,823 | 10,693,191 | 5,198,432 | 2,055,342 |
| 6 h | 307,969,575 | 209,010,076 | 23,853,400 | 12,753,022 | 22,956,346 | 19,087,422 | 6,798,474 | 5,189,855 |
| 7 h | 409,415,300 | 267,238,232 | 67,682,885 | 41,241,553 | 24,791,711 | 19,270,375 | 8,159,486 | 3,077,394 |

*B = Blank (background control). All values are peak areas measured by mass spectrometry (arbitrary units). Data presented as mean ± SD from independent biological replicates. α-KG = α-ketoglutarate.*

**Supplementary Table S4. Pearson and time-controlled partial correlation analyses**

**Partial correlation analysis controlling for time.** To distinguish pairwise metabolic associations from correlations attributable to shared temporal trends, Pearson correlation coefficients were first calculated using the mean values at each of the eight sampling times from 0 to 7 h. First-order partial correlation coefficients controlling for linear time were then calculated as:

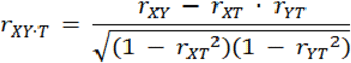

where *r*_*XY*_ is the Pearson correlation between metabolites *X* and *Y*, and *r*_*XT*_ and *r*_*YT*_ are their respective correlations with time *T*. This calculation describes the residual association between *X* and *Y* after adjustment for linear temporal trends. Because only eight time points were available, these analyses should be regarded as exploratory.

The background (blank) sample reported in Table S3-2 was excluded from the correlation analysis, which therefore rests on the eight incubation time points (0–7 h); it is shown alongside the time course to document baseline (pre-incubation) levels of the measured metabolites.

**Table S4-1.**
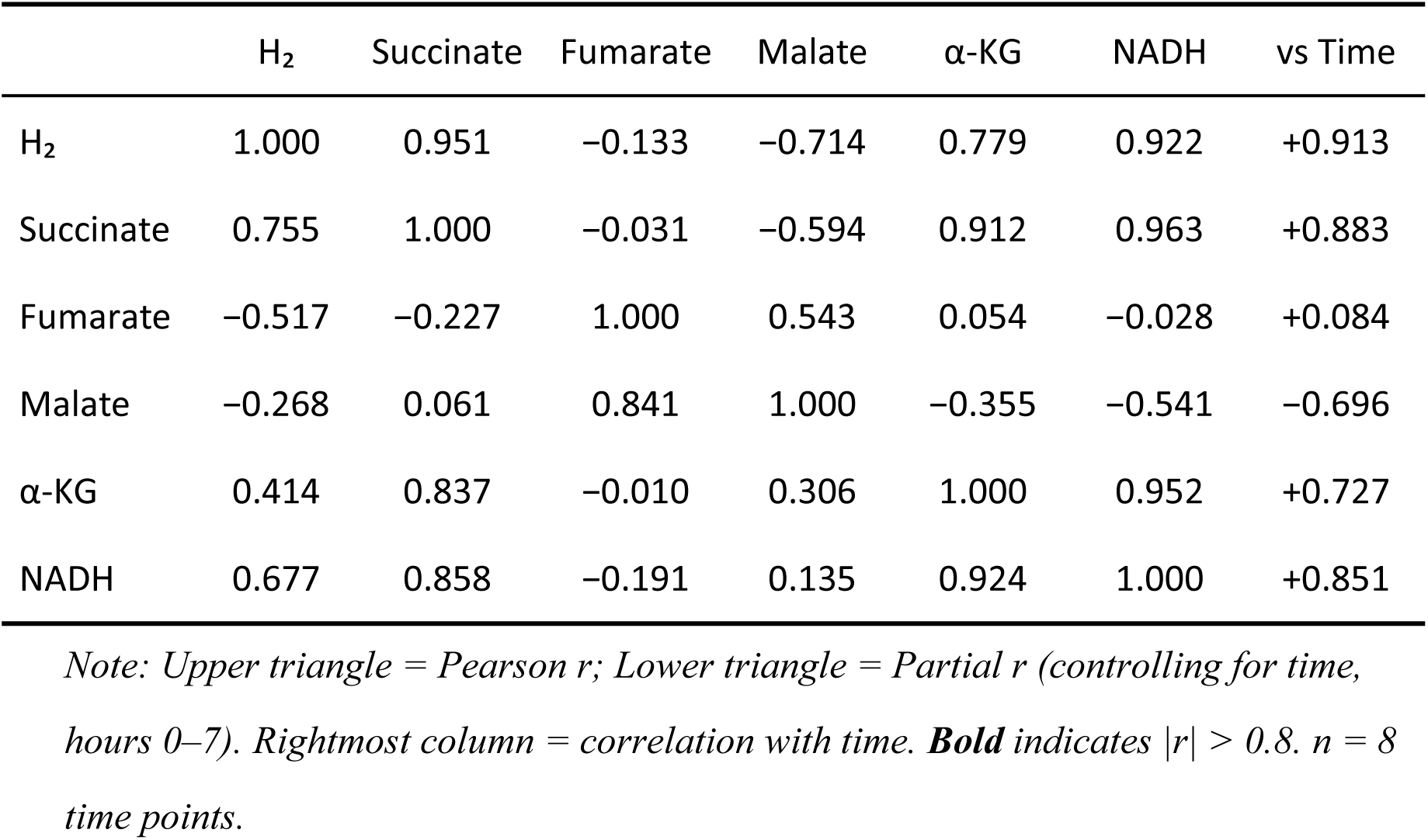
Pearson (upper triangle) and partial (lower triangle, time-controlled) correlation matrices.

|  | H<sub>2</sub> | Succinate | Fumarate | Malate | α-KG | NADH | vs Time |
|---|---|---|---|---|---|---|---|
| H<sub>2</sub> | 1.000 | 0.951 | −0.133 | −0.714 | 0.779 | 0.922 | +0.913 |
| Succinate | 0.755 | 1.000 | −0.031 | −0.594 | 0.912 | 0.963 | +0.883 |
| Fumarate | −0.517 | −0.227 | 1.000 | 0.543 | 0.054 | −0.028 | +0.084 |
| Malate | −0.268 | 0.061 | 0.841 | 1.000 | −0.355 | −0.541 | −0.696 |
| α-KG | 0.414 | 0.837 | −0.010 | 0.306 | 1.000 | 0.952 | +0.727 |
| NADH | 0.677 | 0.858 | −0.191 | 0.135 | 0.924 | 1.000 | +0.851 |

*Note: Upper triangle = Pearson r; Lower triangle = Partial r (controlling for time, hours 0–7). Rightmost column = correlation with time.* ***Bold*** *indicates |r| > 0.8. n = 8 time points.*

**Table S4-2.**
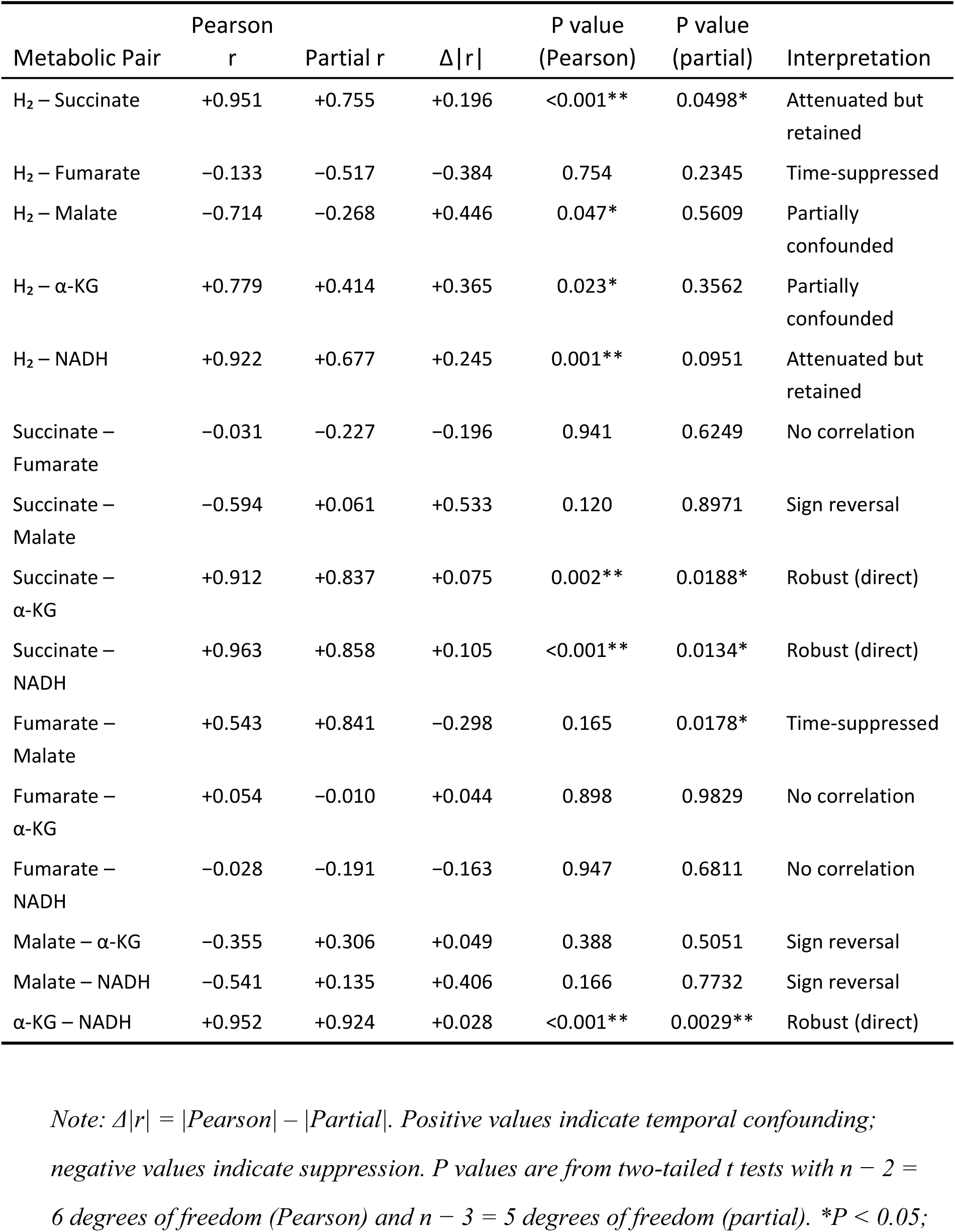
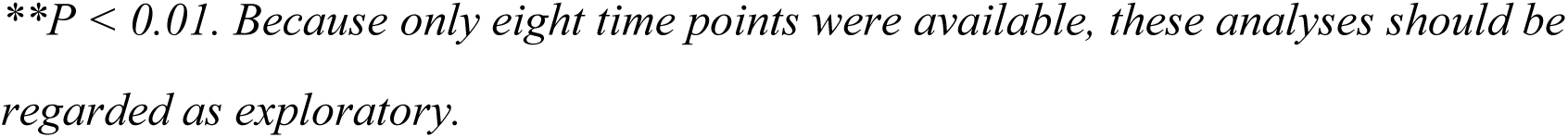
Detailed Pairwise Comparison: effect of temporal confounding on metabolic correlations.

**Table S4-3.** Individual Temporal Trends: correlation of each metabolite with time.

| Variable | r vs Time | 95% CI | P value | Trend |
|---|---|---|---|---|
| H<sub>2</sub> | +0.913 | +0.63 to +0.98 | **0.0015** | Strong accumulation |
| Succinate | +0.883 | +0.53 to +0.98 | **0.0037** | Strong accumulation |
| Fumarate | +0.084 | −0.61 to +0.71 | 0.8424 | No trend |
| Malate | −0.696 | −0.93 to −0.06 | 0.0552 | Depletion (marginal) |
| α-KG | +0.727 | +0.12 to +0.94 | **0.0411** | Accumulation |
| NADH | +0.851 | +0.43 to +0.97 | **0.0074** | Strong accumulation |

*Note: CI = confidence interval by Fisher z-transformation. Bold P < 0.05.*

Time-controlled partial correlation analysis separated shared temporal progression from time-independent metabolic coupling. The associations of H₂ with succinate and NADH were attenuated but retained after adjustment for linear time (H₂–succinate: Pearson r = 0.951, partial r = 0.755; H₂–NADH: Pearson r = 0.922, partial r = 0.677), indicating that co-accumulation over the time course contributes to these relationships without accounting for them.

In contrast, associations linking succinate, α -ketoglutarate and NADH were almost unaffected by time adjustment ( α -ketoglutarate – NADH: Pearson r = 0.952, partial r = 0.924; succinate – NADH: Pearson r = 0.963, partial r = 0.858; succinate – α -ketoglutarate: Pearson r = 0.912, partial r = 0.837). The fumarate – malate association increased from Pearson r = 0.543 to partial r = 0.841, suggesting that their different overall temporal trends obscured their positive covariation in the unadjusted analysis.

### Supplementary Calculations

#### Thermodynamic and Kinetic Calculations

##### Model status note

The calculations below constitute a *working model* — a thermodynamically feasible and internally consistent framework — rather than an established mechanism. Direct spectroscopic or kinetic evidence for branched electron transfer within CI* is not yet available, and the identity of the H₂-forming site remains unresolved. Among the mechanisms considered, an FMN-centered branched electron-transfer mechanism provides the most internally consistent explanation of the current observations. The specific catalytic chemistry (FMN-N5 vs N1a) is therefore presented as two testable hypotheses (S5): Model A (FMN–N1a cooperative) is favored as the primary working model, with Model B (N1a iron direct) retained as a less-supported alternative; discriminating experiments are outlined in S13.

##### Temperature note

All calculations are performed at 25°C (298.15 K) using RT/F = 25.69 mV, unless otherwise stated. At the experimental temperature of 35°C (308.15 K), RT/F = 26.55 mV — a 3.3% difference that marginally increases all Nernst corrections but does not alter any qualitative conclusions or change the sign of any ΔG value. Where relevant, 35°C values are provided in parentheses.

#### S1. Standard Redox Potentials of Relevant Couples

| **Redox couple** | ***E′*<sub>m,7</sub> (mV)** | **Reference** |
|---|---|---|
| NAD<sup>+</sup>/NADH (2e<sup>−</sup>/1H<sup>+</sup>) | −320 | Ohnishi et al., 2018 |
| FMN/FMNH<sub>2</sub> (2e<sup>−</sup>, CI-bound) | −340 | Sled et al., 1994 |
| FMN/FMNH· (1st reduction, *E*<sub>1</sub>) | −390 | Sazanov & Hinchliffe, 2006 |
| FMNH·/FMNH<sub>2</sub> (2nd reduction, *E*<sub>2</sub>) | −300 | Sazanov & Hinchliffe, 2006 |
| N3 [4Fe-4S] | −250 | Ohnishi et al., 2018; Hirst, 2013 |
| N1a [2Fe-2S] | −370 | Ohnishi, 1998; Ohnishi et al., 2018 |
| N2 [4Fe-4S] | −100 | Estimated (range −80 to −220 depending on pH and species; Hirst, 2013; Zwicker et al., 2006). Not used in the S4 bifurcation balance; referenced in S7 (rotenone blocks N2→UQ) |
| UQ/UQH<sub>2</sub> (2e<sup>−</sup>/2H<sup>+</sup>, CI-bound) | +90 | Estimated (90 ± 10 mV; Ohnishi et al., 2018; Mitchell, 1976) |
| UQ/UQ•<sup>−</sup> (1e<sup>−</sup>, CI-bound) | ≈ −160 | Estimated (membrane milieu; range −150 to −200) |
| H<sup>+</sup>/H<sub>2</sub> (pH 7) | −414 | Standard |

**Note:** All standard redox potentials are given relative to the standard hydrogen electrode (SHE) at pH 7.0 and 25°C. Where plant-specific midpoint potentials are unavailable, values determined for bovine heart complex I were used; this is justified by the strong conservation of the cofactor environment and is treated as an assumption in the sensitivity analysis (S9).

##### S1.1 pH 6.0 corrections (experimental condition)

H₂ accumulation experiments were performed at pH 6.0. For proton-coupled couples, the formal potential shifts with pH:

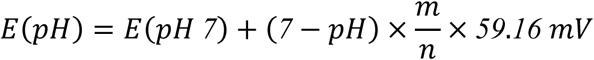

where *n* = number of electrons transferred and *m* = number of protons transferred.

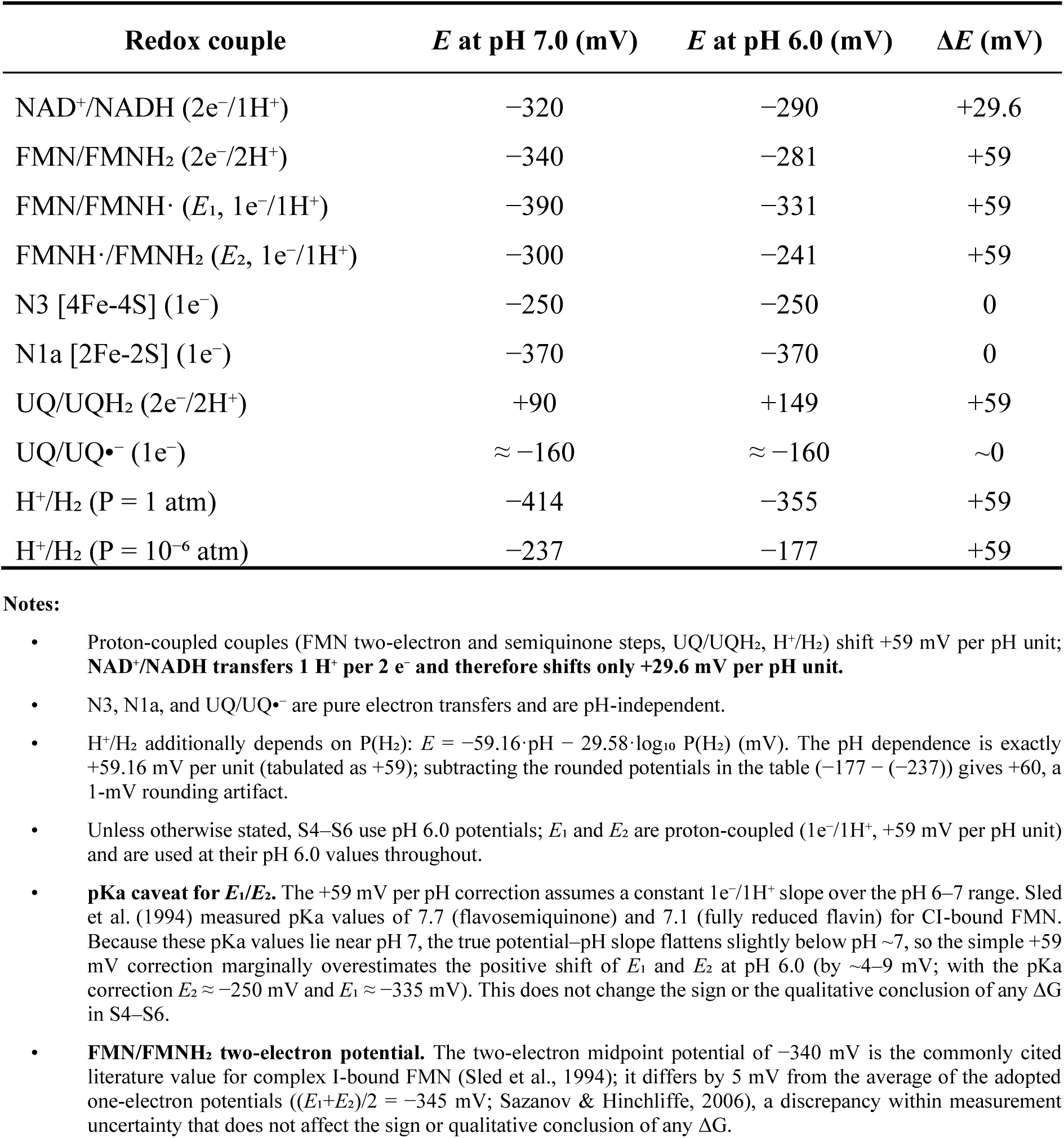

| **Redox couple** | ***E* at pH 7.0 (mV)** | ***E* at pH 6.0 (mV)** | **Δ*E* (mV)** |
|---|---|---|---|
| NAD<sup>+</sup>/NADH (2e<sup>−</sup>/1H<sup>+</sup>) | −320 | −290 | +29.6 |
| FMN/FMNH<sub>2</sub> (2e<sup>−</sup>/2H<sup>+</sup>) | −340 | −281 | +59 |
| FMN/FMNH· (*E*<sub>1</sub>, 1e<sup>−</sup>/1H<sup>+</sup>) | −390 | −331 | +59 |
| FMNH·/FMNH<sub>2</sub> (*E*<sub>2</sub>, 1e<sup>−</sup>/1H<sup>+</sup>) | −300 | −241 | +59 |
| N3 [4Fe-4S] (1e<sup>−</sup>) | −250 | −250 | 0 |
| N1a [2Fe-2S] (1e<sup>−</sup>) | −370 | −370 | 0 |
| UQ/UQH<sub>2</sub> (2e<sup>−</sup>/2H<sup>+</sup>) | +90 | +149 | +59 |
| UQ/UQ•<sup>−</sup> (1e<sup>−</sup>) | ≈ −160 | ≈ −160 | ~0 |
| H<sup>+</sup>/H<sub>2</sub> (P = 1 atm) | −414 | −355 | +59 |
| H<sup>+</sup>/H<sub>2</sub> (P = 10<sup>−6</sup> atm) | −237 | −177 | +59 |

**Notes:**
• Proton-coupled couples (FMN two-electron and semiquinone steps, UQ/UQH<sub>2</sub>, H<sup>+</sup>/H<sub>2</sub>) shift +59 mV per pH unit; **NAD<sup>+</sup>/NADH transfers 1 H<sup>+</sup> per 2 e<sup>−</sup> and therefore shifts only +29.6 mV per pH unit.**
• N3, N1a, and UQ/UQ•<sup>−</sup> are pure electron transfers and are pH-independent.
• H<sup>+</sup>/H<sub>2</sub> additionally depends on P(H<sub>2</sub>): *E* = −59.16·pH − 29.58·log<sub>10</sub> P(H<sub>2</sub>) (mV). The pH dependence is exactly +59.16 mV per unit (tabulated as +59); subtracting the rounded potentials in the table (−177 − (−237)) gives +60, a 1-mV rounding artifact.
• Unless otherwise stated, S4–S6 use pH 6.0 potentials; *E*<sub>1</sub> and *E*<sub>2</sub> are proton-coupled (1e<sup>−</sup>/1H<sup>+</sup>, +59 mV per pH unit) and are used at their pH 6.0 values throughout.
• **pKa caveat for *E*<sub>1</sub>/*E*<sub>2</sub>.** The +59 mV per pH correction assumes a constant 1e<sup>−</sup>/1H<sup>+</sup> slope over the pH 6–7 range. Sled et al. (1994) measured pKa values of 7.7 (flavosemiquinone) and 7.1 (fully reduced flavin) for CI-bound FMN. Because these pKa values lie near pH 7, the true potential–pH slope flattens slightly below pH ~7, so the simple +59 mV correction marginally overestimates the positive shift of *E*<sub>1</sub> and *E*<sub>2</sub> at pH 6.0 (by ~4–9 mV; with the pKa correction *E*<sub>2</sub> ≈ −250 mV and *E*<sub>1</sub> ≈ −335 mV). This does not change the sign or the qualitative conclusion of any ΔG in S4–S6.
• **FMN/FMNH<sub>2</sub> two-electron potential.** The two-electron midpoint potential of −340 mV is the commonly cited literature value for complex I-bound FMN (Sled et al., 1994); it differs by 5 mV from the average of the adopted one-electron potentials ((*E*<sub>1</sub>+*E*<sub>2</sub>)/2 = −345 mV; Sazanov & Hinchliffe, 2006), a discrepancy within measurement uncertainty that does not affect the sign or qualitative conclusion of any ΔG.

#### S2. Thermodynamic Feasibility of FMN-Dependent H₂ Release and the Cooperative FMN-N5/N1a Model

This section establishes the thermodynamic baseline for H₂ formation from reduced flavin under the experimental conditions (pH 6.0, hypoxia) and explains why FCCP abolishes H₂ despite the reaction being intrinsically favorable. A separate subsection (S2.6) summarizes the cooperative FMN-N5/N1a model discussed in the main text and analyzed in detail in S5.

##### S2.1 Standard free energy at pH 7.0 (reference)

###### For the overall two-electron reaction

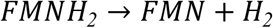

using pH 7.0 standard potentials:

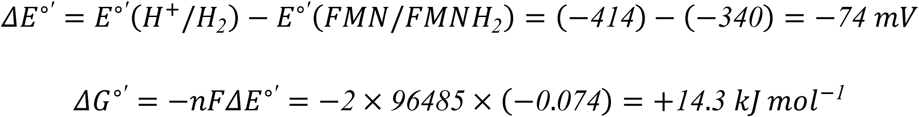

Under standard conditions (1 atm H₂, [FMNH₂]/[FMN] = 1), the reaction is unfavorable.

##### S2.2 Proton reduction potential at pH 6.0 and low P(H₂)

The experimental H₂ production occurs at pH 6.0. For the proton reduction half-reaction

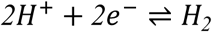

the potential at 25 °C is given by

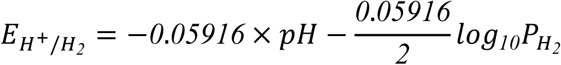

At pH 6.0 and (P(H₂) = 10⁻⁶) atm (conservative estimate from headspace detection limits):

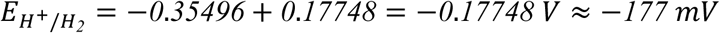

At 35 °C, the corresponding value is ≈ −183 mV. This value is used as the cathode potential throughout this section.

##### S2.3 FMN/FMNH₂ potential at pH 6.0 and high reduction ratio

The FMN/FMNH₂ couple is a 2e⁻/2H⁺ process. At pH 6.0, using the pH 7.0 value of −340 mV and a shift of +59.2 mV per pH unit:

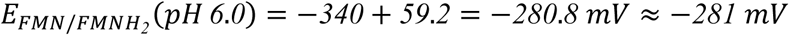

Under hypoxia, the ubiquinone pool becomes highly reduced (as O₂ becomes limiting) and NADH accumulates, establishing a strongly reducing FMN pool. For the direct two-electron pathway considered here, we assume a high [FMNH₂]/[FMN] ratio ≈ 100:1:

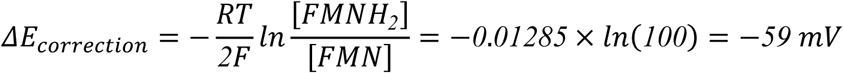

Thus,

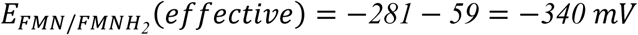

This is the anode potential for the direct two-electron FMNH₂ → H₂ reaction.

##### S2.4 Net driving force for direct FMNH₂ → H₂ at pH 6.0

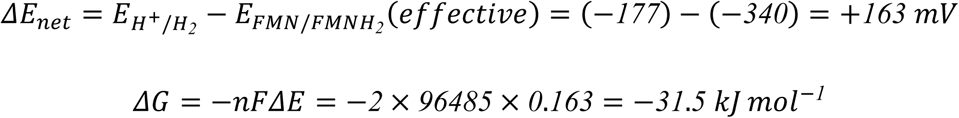

At 35 °C, Δ*E*_net ≈ +170 mV, giving ΔG ≈ −32.8 kJ mol⁻¹.

Thus, the direct two-electron oxidation of FMNH₂ to H₂ is strongly spontaneous under the experimental conditions. This calculation establishes a thermodynamic baseline but does not imply that FMN alone is the catalytic site.

##### S2.5 Thermodynamic consequences of Δp collapse: rationale for FCCP inhibition

The intrinsic FMN→H₂ reaction (S2.4) is favorable, so FCCP cannot act by altering the H₂-forming step itself. Instead, FCCP must act upstream, by removing the conditions that make the reaction favorable:

1. **RET eliminated.** Collapse of Δp abolishes reverse electron transport (CII → UQ → CI), removing the dominant NADH source for CI*.
2. **NADH pool depleted.** Without Δp backpressure, mature complex I operates at maximal forward rate, rapidly oxidizing the matrix NADH pool.
3. **FMNH₂/FMN ratio collapses.** As FMNH₂/FMN drops from ≈100 toward ≈1, the −59 mV correction vanishes, and the effective anode potential becomes −281 mV instead of −340 mV. The net driving force then becomes *ΔE* = −*177* − (−*281*) = +*104 mV*, still exergonic, but the reaction becomes kinetically limited by substrate availability and may cease if NADH supply is fully exhausted.

This interpretation is supported by the observed FCCP inhibition and the known role of Δp in driving RET. However, direct measurements of matrix NADH and FMN redox state under FCCP treatment were not performed in this study.

###### Conclusion

Δp’s role is **permissive** — maintaining the metabolic conditions (active RET, high NADH, high FMNH₂/FMN) under which the FMN→H₂ reaction is intrinsically thermodynamically favorable, rather than directly coupling transmembrane proton translocation to H₂ production. This subsection is therefore not an additional thermodynamic calculation but a condition analysis that links the thermodynamic feasibility (S2.4) to the observed pharmacology.

##### S2.6 Thermodynamics of the cooperative FMN-N5/N1a model (summary)

The main text considers a model in which H₂ is formed cooperatively by the FMN semiquinone and reduced N1a:

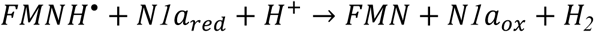

A detailed step-by-step analysis is given in S5. In brief, the two anodic half-reactions are FMNH· → FMN + e⁻ + H⁺ (*E*₁ = −331 mV at pH 6.0) and N1a(red) → N1a(ox) + e⁻ (*E* = −370 mV), giving an average anode potential of (−331 + (−370))/2 = −350.5 mV. With the cathode at *E*(H⁺/H₂) = −177 mV (pH 6.0, 10⁻⁶ atm H₂), the net driving force is Δ*E* = +173 mV, giving ΔG ≈ −33 kJ mol⁻¹.

This cooperative pathway is comparable in magnitude to the direct FMNH₂ → H₂ baseline (−31.5 kJ mol⁻¹), the small difference being within the uncertainty of the assumed midpoint potentials; it uses only one electron from FMNH· and one from N1a, matching the stoichiometry expected from a bifurcation-inspired mechanism in which one electron is delivered to the high-potential branch and one to N1a.

##### S2.7 Relation to main text models

The calculations in S2.4 and S2.6 show that both the direct FMN two-electron reaction and the cooperative FMN-N5/N1a reaction are thermodynamically favorable under the experimental conditions. However, they do not establish which pathway is kinetically competent. The main text presents both as testable hypotheses; the small difference in their calculated free energies (≈2 kJ mol⁻¹) is within the uncertainty of the assumed midpoint potentials and therefore does not discriminate between them. A more discriminating distinction is their pH dependence: the direct two-electron route carries no net proton stoichiometry and is pH-insensitive (S2.4), whereas the cooperative reaction consumes one matrix proton and becomes ≈5.7 kJ mol⁻¹ more exergonic per unit decrease in pH (S10.1), directly accounting for the experimentally observed acidic pH optimum. The cooperative model is therefore emphasized, in addition to its avoidance of the requirement for FMN to deliver two electrons to protons in a single step and its alignment with a bifurcation-inspired electron-partitioning architecture.

The high [FMNH₂]/[FMN] ratio assumed in S2.3 applies only to the conditional-potential estimate for the direct two-electron FMNH₂ → H₂ reaction and does not describe the redox state of the catalytic cycle in Model A. In the cooperative mechanism, FMN cycles between its oxidized, semiquinone, and reduced states during turnover, and the FMNH· required for H₂ formation is generated transiently by bifurcation in the second half of the ping-pong cycle (S6.1), rather than from a fully reduced FMN pool.

#### S3. One-Electron Potentials of CI-Bound FMN: Semiquinone Instability

##### S3.1 Measured one-electron potentials of the mature CI FMN

The FMN cofactor of complex I accepts two electrons from NADH and releases them one at a time to the Fe-S chain. Its two one-electron reduction potentials are:

- *E*₁: FMN + e⁻ + H⁺ → FMNH· (first reduction)
- *E*₂: FMNH· + e⁻ + H⁺ → FMNH₂ (second reduction)

Measured values for the intact enzyme are:

| **Source** | ***E*<sub>1</sub> (FMN/FMNH·)** | ***E*<sub>2</sub> (FMNH·/FMNH<sub>2</sub>)** | **Semiquinone** |
|---|---|---|---|
| Sled et al. (1994), pH 7.5 | −414 mV | −336 mV | unstable (*E*<sub>1</sub> < *E*<sub>2</sub>) |
| Sazanov & Hinchliffe (2006), pH 7 | −390 mV | −300 mV | unstable (*E*<sub>1</sub> < *E*<sub>2</sub>) |

We adopt the Sazanov & Hinchliffe (2006) values, *E*₁ = −390 mV and *E*₂ = −300 mV at pH 7.0.

##### S3.2 pH 6.0 correction

Both *E*₁ and *E*₂ are 1e⁻/1H⁺ processes. At pH 6.0 each shifts +59 mV:

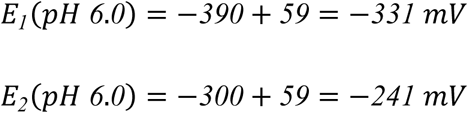

The ordering *E*₁ < *E*₂ (unstable semiquinone) is preserved at pH 6.0.

##### S3.3 Semiquinone stability

The semiquinone formation constant *K*_sf, defined by FMN + FMNH₂ ⇌ 2 FMNH·, relates to the potential difference by:

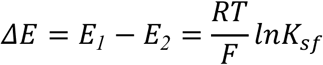

With *E*₁ − *E*₂ = −90 mV, *K*_sf ≈ 0.03.

*K*_sf < 1 indicates that the semiquinone is thermodynamically unstable at equilibrium and would disproportionate. This instability does not, however, preclude a catalytically competent semiquinone: Sled et al. (1994) detected the flavosemiquinone by EPR during turnover, indicating a lifetime sufficient for electron partitioning. As with many enzymatic reactions, a high-energy intermediate need not be thermodynamically stable to be kinetically trapped and productively converted.

##### S3.4 Implication for bifurcation

The mature CI semiquinone is thermodynamically unstable (*E*₁ < *E*₂, crossed potentials), a feature characteristic of flavin-based electron bifurcation (Buckel & Thauer, 2018). Using the measured (unstable-semiquinone) potentials directly, at pH 6.0 the high-potential branch (FMNH₂ → UQ) is exergonic and the low-potential branch (FMNH· → N1a) is endergonic — the thermodynamic signature of bifurcation. The transient semiquinone, even if thermodynamically unstable, is kinetically captured by N1a, which lies within electron-tunneling distance of FMN. Thus, complex I exhibits the thermodynamic signature of electron bifurcation (an exergonic high-potential branch coupled to an endergonic low-potential branch).

#### S4. Electron Bifurcation Energetics

Electron **bifurcation** partitions the two electrons of a single reduced flavin (FMNH₂) into two branches of different potential: one electron is delivered to the high-potential branch (N3 → UQ), the other to the low-potential branch (N1a). The exergonic high-potential transfer subsidizes the endergonic low-potential transfer. The one-electron potentials used below (*E*₁ = −331 mV, *E*₂ = −241 mV at pH 6.0) follow the simple +59 mV per pH correction; the pKa refinement (S1.1) shifts them by only ∼4–9 mV and does not change any sign or qualitative conclusion.

##### S4.1 High-potential branch: FMNH₂ → FMNH· + e⁻ → Fe-S chain → UQ

The first oxidation of FMNH₂ (reverse of *E*₂) releases one electron at *E*₂(pH 6.0) = −241 mV. This electron is relayed through N3 [4Fe-4S] (−250 mV) and the downstream Fe-S chain to ubiquinone, where the first electron transfer is a single-electron reduction:

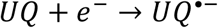

with *E*_*UQ*/*UQ*_•− ≈ −*160 mV* (vs SHE). The overall driving force of the high-potential branch is:

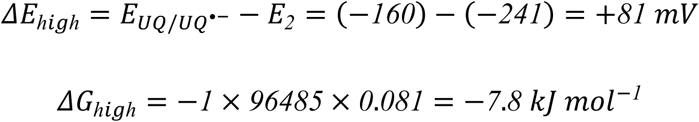

(The FMNH₂ → N3 step itself is near equilibrium at pH 6.0, Δ*E* = −9 mV; the exergonic drive accumulates along the Fe-S chain toward UQ•⁻.)

This exergonic release is the thermodynamic driving force that subsidizes the low-potential branch.

##### S4.2 Low-potential branch: FMNH· → FMN + e⁻ → N1a

The second oxidation (reverse of *E*₁) releases one electron at *E*₁(pH 6.0) = −331 mV to N1a (−370 mV):

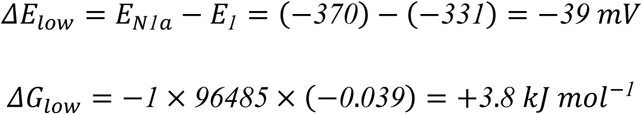

This step is **endergonic** — the hallmark of electron bifurcation, where an uphill transfer is driven by coupling to the exergonic branch.

##### S4.3 Bifurcation energy balance

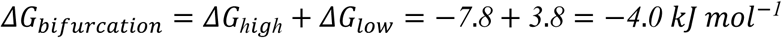

The overall bifurcation is exergonic (ΔG ≈ −4.0 kJ mol⁻¹) but only modestly so, consistent with a favorable yet regulated, low-flux side reaction. This net driving force supports a bifurcation-like mechanism in CI*, where the intact Q module and ubiquinone-binding site allow the high-potential branch to terminate at UQ, while AOX recycles UQH₂ under micro-oxic conditions.

##### S4.4 pH dependence of the low-potential branch — the pH ≈ 6.7 divide

###### Rationale

The endergonic character of the low-potential branch (S4.2) is not fixed but depends strongly on pH, because the two couples involved respond differently to proton activity. *E*₁ (FMN/FMNH·) is proton-coupled: the flavosemiquinone carries a proton at N5 (FMNH·), so its one-electron oxidation involves one proton and its midpoint potential shifts +59.16 mV per pH unit (S1.1). By contrast, N1a is a [2Fe-2S] cluster whose reduction is a pure electron transfer with no proton stoichiometry, so *E*(N1a) = −370 mV is pH-independent. The driving force of the low-potential branch,

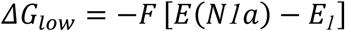

therefore varies with pH.

###### Derivation of the divide

With *E*₁(pH) = *E*₁(pH 7) + 59.16 mV × (7 − pH) = −390 + 59.16 × (7 − pH) mV:

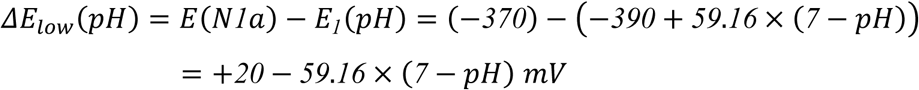

Setting Δ*E*_low = 0 gives 7 − pH = 20/59.16 = 0.338, i.e. **pH = 6.7**. Below this pH the branch is endergonic (uphill); above it, exergonic (downhill).

| **pH** | ***E*<sub>1</sub> (mV)** | **Δ*E*_low = *E*(N1a) − *E*<sub>1</sub> (mV)** | **ΔG_low (kJ mol<sup>−1</sup>)** | **Branch direction** |
|---|---|---|---|---|
| 6.0 | −331 | −39 | +3.8 | endergonic (uphill) |
| 6.5 | −360.5 | −9.5 | +0.9 | endergonic |
| 6.7 | −370 | 0 | 0 | divide |
| 7.0 | −390 | +20 | −1.9 | exergonic (downhill) |
| 7.5 | −419.5 | +49.5 | −4.8 | exergonic |
| ≥7.7 | −431 | +61 | −5.9 | exergonic (plateau) |

###### pKa refinement

The +59 mV per pH correction assumes the semiquinone remains protonated. Sled et al. (1994) measured pKa = 7.7 for the flavosemiquinone, above which FMNH· deprotonates to the anionic FMN·⁻ and *E*₁ no longer shifts with pH; ΔG_low therefore reaches a plateau of −5.9 kJ mol⁻¹ at pH ≥ 7.7 (*E*₁ = −431 mV). This refinement shifts *E*₁ by only ∼4–9 mV near pH 6–7 and does not move the divide (pH ≈ 6.7, well below the pKa) or change the sign of any value.

###### Two regimes

1. **pH 6.0 (experimental condition).** ΔG_low = +3.8 kJ mol⁻¹ is endergonic. The low-potential branch is a genuine “uphill” transfer that must be driven by the exergonic high-potential branch (ΔG_high = −7.8 kJ mol⁻¹). This energy coupling — an exergonic branch driving an endergonic one — is the defining feature of electron bifurcation. At pH 6.0 the FMN–N1a dyad therefore would satisfy the thermodynamic condition for bifurcation in the proposed model.
2. **Physiological pH (≥7.0).** ΔG_low is exergonic (−1.9 to −5.9 kJ mol⁻¹). The second electron would partition down-hill to N1a without requiring coupling, and the “bifurcation” would degenerate into simple two-electron partitioning. Consistent with this, in the canonical enzyme under normal conditions N1a is largely bypassed: the second electron follows the fast N3 main path (strong FMN–N3 spin coupling; Sled et al., 1994), and N1a reduction is observed only under strongly reducing conditions (Grivennikova et al., 2024), consistent with its midpoint potential lying below that of the NADH/NAD⁺ couple (Ohnishi et al., 2018).

###### Physiological significance

The divide at pH ≈ 6.7 places the acidic pH 6.0 optimum of our H₂ assays on the endergonic side of the divide. Matrix pH has not been measured directly in hypoxic plant mitochondria, but the cytosol of anoxic plant tissues acidifies to ≈ pH 6.8 (Roberts et al., 1984; Gout et al., 2001), and the matrix is expected to follow cytosolic acidification (Schwarzländer & Finkemeier, 2013). Acidic pH thus serves two functions simultaneously: (i) it makes the cooperative H₂-forming step strongly exergonic (S5.1), and (ii) it converts the N1a branch into the endergonic, high-potential-branch-coupled step that defines genuine bifurcation. At physiological matrix pH (∼7.5–8.0) neither condition holds: the H₂-forming step is endergonic (S5.1), and the second electron partitions down-hill to N1a without coupling. These two requirements are interlocking rather than independent: the high-pH partitioning regime is chemically silent for H₂ formation, so acidic pH is simultaneously the enabling condition for the chemistry and the enforcing condition for the coupling. This interlock is a key distinction between our model and a simple “electron partitioning” interpretation of the rotenone paradox (S7).

###### Caveat on *E*(N1a)

We use *E*(N1a) = −370 mV, the established *E*ₘ,₇.₀ for bovine complex I (Ohnishi, 1998; Ohnishi et al., 2018), throughout. The SI uses −370 mV consistently with S1.

(Values use the simple +59 mV per pH correction unless otherwise stated; the pKa refinement of S1.1 shifts *E*₁ by ∼4–9 mV and does not move the divide or change any sign.)

#### S5. H₂ Formation Step: Cooperative Proton Reduction by FMNH· and N1a(red) (electron confurcation)

Electron bifurcation (S4) does not, within a single turnover, generate FMNH· and N1a(red) together: the first branched turnover reduces N1a and regenerates oxidized FMN, and only in the second turnover is the flavin trapped as FMNH· while N1a remains reduced (S6.1). The H₂-forming step therefore combines two one-electron donors of different potential — FMNH· (*E*₁ = −331 mV) and N1a(red) (−370 mV) — converging on the 2H⁺/H₂ acceptor. At near-equilibrium H₂ pressure (pH 6.0, 1 atm; *E*(H⁺/H₂) = −355 mV, intermediate between the two donors) this constitutes a genuine electron **confurcation** — the reverse of bifurcation — in which exergonic donation from N1a(red) subsidizes the mildly endergonic donation from FMNH·. At the low experimental P(H₂) = 10⁻⁶ atm (*E* = −177 mV, higher than both donors), however, donation from either donor is independently favorable, and the step is more precisely a convergent two-electron proton reduction (Furlan et al., 2022). We consider two alternative models for this step, which share the same stoichiometry but differ in the catalytic site (S5.1 and S5.3).

##### Structural rationale

Unlike the electron-bifurcating [FeFe] hydrogenase HydABC, which uses a dedicated H-cluster to evolve H₂ (Furlan et al., 2022), CI* contains no hydrogenase cofactor. The CI* structure (Maldonado et al., 2020) retains all cofactors of the NADH-to-CoQ pathway — the FMN (in NDUV1), seven main-pathway FeS clusters, and the off-pathway N1a [2Fe-2S] cluster (in NDUV2) — but lacks the distal PD module. Among the available cofactors, the FMN N5 position (a hydride-transfer site) and the off-pathway, low-potential N1a cluster are therefore the most plausible H₂-forming components (Model A); alternatively, N1a itself could act as the catalytic site if a PD-loss-induced conformational change exposes its iron (Model B). We cannot exclude that a CI*-specific feature — such as the plant-specific γ-carbonic-anhydrase (γCA) domain retained in CI*, or a conformational change associated with PD loss — additionally participates in proton delivery to the H₂-forming site.

##### N1a-related energetics

All N1a calculations below use the pH-independent midpoint potential *E*(N1a) = −370 mV from S1. The reduction of N1a by FMNH· is treated in S4.2; the use of N1a(red) in H₂ formation is analyzed here in Models A and B.

##### S5.1 Model A — cooperative FMN-N5 / N1a catalysis

In this model, the N5-bound hydrogen of FMNH· provides one electron and one proton equivalent, while N1a(red) provides the second electron:

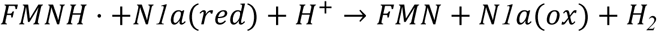

This 2e⁻/2H⁺ process can be decomposed into three half-reactions:

| **Half-reaction** | **Potential (mV)** | **Comment** |
|---|---|---|
| FMNH· → FMN + e<sup>−</sup> + H<sup>+</sup> | *E*<sub>1</sub> = −390 (pH 7); −331 (pH 6) | 1e<sup>−</sup> oxidation of semiquinone |
| N1a(red) → N1a(ox) + e<sup>−</sup> | *E*(N1a) = −370 | 1e<sup>−</sup> oxidation of [2Fe-2S] |
| 2H<sup>+</sup> + 2e<sup>−</sup> → H<sub>2</sub> | *E*°′(H<sup>+</sup>/H<sub>2</sub>) = −414 (pH 7, 1 atm) | 2e<sup>−</sup> proton reduction |

The overall driving force depends on the average potential of the two anodic processes:

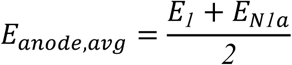

Under standard conditions (pH 7.0, P(H₂) = 1 atm):

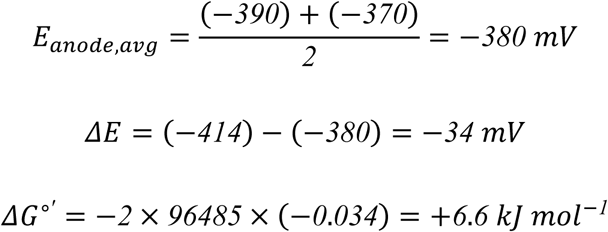

Under standard conditions, the cooperative reaction is **mildly unfavorable**.

Under experimental conditions (pH 6.0, P(H₂) ≈ 10⁻⁶ atm):

The hydrogen-electrode potential shifts positively:

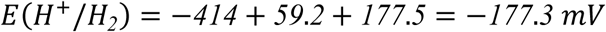

With *E*₁ as a proton-coupled process (1e⁻/1H⁺, +59 mV per pH unit):

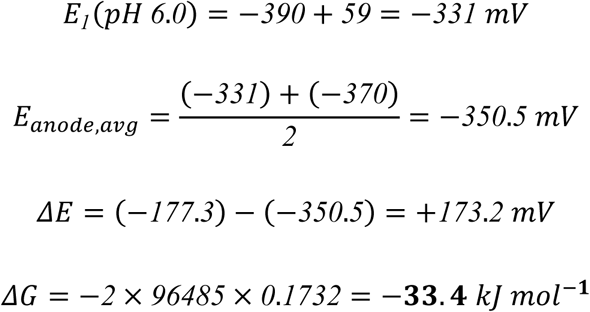

###### Conclusion

Under micro-oxic, acidic conditions, the cooperative reduction of protons by FMNH· and N1a(red) is **strongly exergonic** (ΔG ≈ −33 kJ mol⁻¹). The low-potential N1a cluster (−370 mV), far from being a thermodynamic dead-end, is proposed to serve as the second electron donor for H–H bond formation at the flavin site.

###### Mechanism of Model A — direct cooperative proton reduction (no FMNH₂ intermediate)

Electron bifurcation (S4) “charges” the low-potential N1a cluster (N1a(red)) at the expense of the exergonic high-potential branch; in the second turnover, N1a(red) cooperates with the FMNH· generated at the FMN N5 site to release H₂ by proton-coupled electron transfer (PCET), which may be concerted or tightly sequential, returning N1a to the oxidized state and closing the 2 NADH : 1 H₂ cycle:

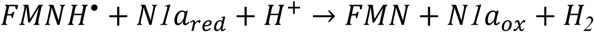

The N1a electron is delivered through the protein matrix to the FMN N5 site by electron tunneling (∼11.8 Å; S8) and combines with the N5-bound hydrogen and a solvent proton in a single coupled step, without generating a stable FMNH₂ intermediate. This direct route is consistent with the rotenone sensitivity (S7): by blocking quinone reduction, rotenone saturates the Fe-S chain and thereby precludes formation of FMNH·, the required co-substrate, abolishing H₂ production.

The chemical plausibility of this step rests on three independently supported elements: the N5 position of FMN is a well-established hydride-transfer site, analogous to the dihydropyridine C4 of NADH (Norcross et al., 1962; Walsh, 1980), and H–H bond formation may proceed by proton-coupled electron transfer (Hammes-Schiffer, 2015). None of these elements, however, constitutes a natural precedent for flavin-catalyzed H₂ formation: the N5–H of the neutral semiquinone is a hydrogen-atom equivalent coupled to the flavin radical electronic structure rather than a classical metal hydride, and whether the H–H bond-forming step is strictly concerted or asynchronous (with a short-lived intermediate) remains undetermined.

We note that electron back-transfer from N1a(red) to FMNH· to form FMNH₂ is thermodynamically favorable (Δ*E* = *E*₂ − *E*(N1a) = (−241) − (−370) = +129 mV; ΔG = −12.4 kJ mol⁻¹) and may occur as a non-productive off-pathway reaction. However, this back-transfer does not lead to H₂ formation in CI* under our conditions: if FMNH₂ were a productive H₂-forming intermediate, rotenone-induced FMNH₂ accumulation should have enhanced H₂ production, the opposite of the observed strong inhibition (S7). We therefore regard the back-transfer as an off-pathway branch that merely returns the electron to the flavin, not as the catalytic H₂-forming step.

###### Sensitivity to H₂ partial pressure

The H₂-forming driving force depends on P(H₂) through the Nernst term of the H⁺/H₂ couple; at 25 °C, *E*(H⁺/H₂) rises by 29.6 mV, and the two-electron ΔG falls by 5.71 kJ mol⁻¹, for every 10-fold decrease in P(H₂). Accordingly, at pH 6.0 the cooperative reaction becomes favorable (ΔG < 0) already below P(H₂) ≈ 0.70 atm (and is nearly neutral, +0.9 kJ mol⁻¹, even at 1 atm), whereas at pH 7.0 the threshold is ≈0.07 atm. By Henry’s law (k_H ≈ 800 μM atm⁻¹ at 25 °C), the pH 6.0 threshold corresponds to a dissolved H₂ concentration of ≈0.56 mM. The value P(H₂) = 10⁻⁶ atm (≈1 ppm; ≈0.8 nM dissolved) used above represents a strongly H₂-depleted, thermodynamically favorable limit (corresponding to a sealed-vial headspace in which H₂ does not accumulate), rather than a conservative assumption. Importantly, the calculated equilibrium threshold of ≈0.70 atm at pH 6.0 shows that the reaction is not predicted to require such an extremely low H₂ pressure: matrix acidification alone renders H₂ formation exergonic at near-atmospheric H₂, so the model retains substantial thermodynamic headroom against product (H₂) accumulation.

##### S5.2 Electron and proton stoichiometry

The cooperative mechanism rigorously conserves charge:

| Species | Electrons | Protons | Charge balance |
|---|---|---|---|
| FMNH· | 1e<sup>−</sup> | 1H<sup>+</sup> (N5-H) | Neutral |
| N1a(red) | 1e<sup>−</sup> | 0 | −1 (relative to ox) |
| H<sup>+</sup> (matrix) | — | 1H<sup>+</sup> | +1 |
| **Total reactants** | **2e<sup>−</sup>** | **2H<sup>+</sup>** | **0** |
| FMN | 0 | 0 | 0 |
| N1a(ox) | 0 | 0 | 0 |
| H<sub>2</sub> | 2e<sup>−</sup> | 2H<sup>+</sup> | 0 |
| **Total products** | **2e<sup>−</sup>** | **2H<sup>+</sup>** | **0** |

###### The two electrons in H₂ originate from two distinct one-electron donors

FMNH· (N5-bound hydrogen electron) and N1a(red) ([2Fe-2S] cluster electron). This resolves the electron-accounting paradox of the previous model, in which a single NADH (2e⁻) was simultaneously assigned to the high-potential branch, the low-potential branch, and H₂ formation.

##### S5.3 Model B — direct catalysis by the N1a iron (Fe–H⁻ mechanism)

As an alternative, the N1a [2Fe-2S] cluster itself could provide the catalytic site. If the absence of the PD module permits a local conformational change upon reduction of N1a, one iron atom may become solvent-exposed and form a low-coordinate Fe site analogous to the distal iron of [FeFe]-hydrogenases, catalyzing proton reduction through an Fe–H⁻ intermediate:

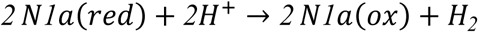

Because N1a is pH-independent (−370 mV), the driving force at pH 6.0 is:

- Standard state (P(H₂) = 1 atm): Δ*E* = (−355) − (−370) = +15 mV; ΔG = −2 × 96485 × 0.015 = **−2.9 kJ mol⁻¹**
- Low P(H₂) = 10⁻⁶ atm: Δ*E* = (−177) − (−370) = +193 mV; ΔG = −2 × 96485 × 0.193 = −37.2 kJ mol⁻¹

Both are favorable. Metal-hydride-mediated H₂ formation is well established in dedicated hydrogenases (Lubitz et al., 2014; Reijerse et al., 2017; Schilter et al., 2016), but its transferability to a fully ligated [2Fe-2S] cluster is unproven; this model would additionally require that N1a — which is fully coordinated and buried in the available structures (Maldonado et al., 2020) — undergo a conformational rearrangement that has not been observed. Moreover, this model requires two equivalents of N1a(red); because each CI* monomer contains a single N1a cluster, this would require either two successive catalytic turnovers or a transient dimerization of CI*, neither of which has been demonstrated — an additional limitation of Model B.

###### Two-step mechanism and intermediate lifetime

If N1a is the catalytic site, proton reduction would proceed in two sequential one-electron steps: (i) N1a(red) + H⁺ → N1a(ox) + Fe–H (a transient iron-bound hydride/hydrogen-atom intermediate); (ii) a second N1a(red) + H⁺ delivers the second electron, converting the Fe–H intermediate to H₂ (2 N1a(red) + 2 H⁺ → 2 N1a(ox) + H₂). For this to be kinetically competent, the putative Fe–H intermediate must persist long enough for the second electron to arrive. In [FeFe]-hydrogenases, metal-hydride intermediates are stabilized by a low-coordinate iron, a hydrophobic pocket, and CO/CN⁻ ligands (Lubitz et al., 2014; Schilter et al., 2016). In CrHydA1, the Hhyd state was reported to form with a time constant of approximately 60 μs and to decay on the order of 1 ms under photoinduced turnover conditions (Sanchez et al., 2019). If the second electron is delivered by a transiently associated CI* monomer, the relevant timescale is set by the encounter rate of two CI* complexes, which may be too slow for a stable Fe–H intermediate; if instead both electrons are supplied sequentially by the same N1a through two successive reduction events, the intermediate would need to persist across an entire turnover cycle, which is less likely for a solvent-exposed iron site. These considerations do not rule out Model B, but they show that it requires a yet-uncharacterized stabilization mechanism or a dedicated proton/electron-delivery pathway—an additional limitation beyond the structural observation that N1a is fully coordinated and buried (Maldonado et al., 2020).

##### S5.4 Comparison of models A and B

|  | **Model A (FMN-N5 / N1a cooperative)** | **Model B (N1a iron direct)** |
|---|---|---|
| H<sub>2</sub> 2e<sup>−</sup> source | FMNH· N5-H (1e<sup>−</sup>) + N1a(red) (1e<sup>−</sup>) | 2 × N1a(red) (2 × 1e<sup>−</sup>) |
| ΔG (pH 6.0, 10<sup>−6</sup> atm) | −33.4 kJ mol<sup>−1</sup> | −37.2 kJ mol<sup>−1</sup> |
| Chemical precedent | organic hydride (no natural flavin hydrogenase) | metal hydride ([FeFe] precedent) |
| Structural constraint | requires N5 solvent access (PD-less may favor) | N1a buried; requires conformational change |
| Reaction topology | intramolecular (FMN–N1a dyad, 11.8 Å) | intermolecular (two CI* monomers) |
| Discriminating test | EPR detects FMNH·; mutate N5 proton-channel residues | XAS / mutate N1a cysteine ligands |

Both models are thermodynamically favorable and consistent with a bifurcation-inspired architecture; they differ in the chemical identity of the H₂-forming site (see S13). We favor **Model A as the primary working model**: it proceeds intramolecularly within a single CI* monomer, requires no unobserved alteration of the N1a coordination, and assumes only that the N5 site is solvent-accessible (a plausible consequence of PD-module loss). **Model B**, by contrast, would require both a yet-unobserved N1a conformational rearrangement (exposure of a low-coordinate iron, absent in the available structures) and either two successive turnovers or a transient dimerization of CI* to supply two N1a(red) equivalents. We therefore retain Model B as a less-supported alternative — considered for transparency and as a target of the discriminating experiments in S13 — rather than as a co-equal candidate.

#### S6. Overall Energy Balance

##### S6.1 Net reaction stoichiometry

For the proposed UQ-coupled two-turnover cycle, electron conservation gives a stoichiometry of two NADH molecules per H₂ released:

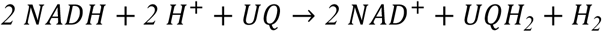

**Verification:**

- Electrons: 2 NADH (4e⁻) → UQH₂ (2e⁻) + H₂ (2e⁻) ✓
- Protons: Left: 2H⁺ + 2H (from NADH hydride) = 4H; Right: UQH₂ (2H) + H₂ (2H) = 4H ✓
- NAD⁺/NADH: 2 NADH consumed, 2 NAD⁺ produced ✓

This stoichiometry reflects the proposed ping-pong catalytic cycle at the FMN–N1a dyad. In the first turnover, one NADH reduces FMN to FMNH₂, which undergoes complete bifurcation—one electron to N3 (high-potential branch) and one to N1a (low-potential branch)—regenerating oxidized FMN while leaving N1a in the reduced state (N1a(red)). In the second turnover, a second NADH reduces FMN again; this time only one electron is released to N3, so the flavin is trapped as the semiquinone FMNH· (carrying a hydrogen at N5). The second electron cannot leave via the low-potential branch because N1a is already reduced from the first turnover and is therefore unavailable as an electron acceptor. The FMNH· produced in the second turnover then pairs with the N1a(red) accumulated from the first turnover to form H₂ (S5), thereby closing the 2 NADH : 1 H₂ : 1 UQH₂ stoichiometry of the proposed cycle. If FMNH₂ were instead converted directly to FMN and H₂, the corresponding stoichiometry would be 1 NADH : 1 H₂ and would not require coupled UQ reduction.

##### S6.2 Thermodynamics of the net reaction

Using the net reaction above:

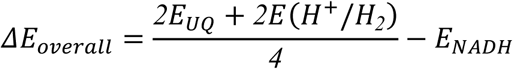

###### At pH 6.0, P(H₂) = 10⁻⁶ atm (experimental conditions)

using pH 6.0 potentials throughout (S1.1) — *E*(NAD⁺/NADH) = −290 mV, *E*(UQ/UQH₂) = +149 mV, *E*(H⁺/H₂) = −177 mV:

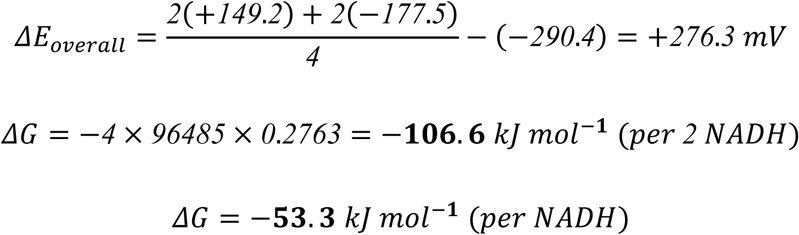

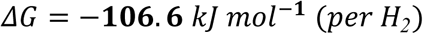

(At 35°C: ΔG ≈ −110 kJ mol⁻¹ per H₂)

The overall cycle is **strongly exergonic** under the assumed model conditions. For reference, the net reaction remains exergonic at pH 7.0 with standard P(H₂) = 1 atm (ΔG ≈ −61 kJ mol⁻¹), but the driving force is substantially larger at pH 6.0 with low P(H₂), underscoring that acidic pH and low H₂ partial pressure each contribute to the favorable thermodynamics (the net reaction consumes 2 H⁺ and therefore depends on pH).

#### S7. Model-Based Interpretation of the Rotenone Effect on H₂ Production

##### S7.1 Problem statement

Rotenone blocks N2 → UQ electron transfer. Within the proposed cooperative model, this blockade is predicted to inhibit H₂ formation through two coupled consequences:

1. **N1a(red) cannot be generated.** The low-potential branch of bifurcation is blocked because N3 cannot accept the high-potential electron from FMNH₂. Without prior bifurcation, N1a remains oxidized (N1a(ox)), depriving the H₂-forming step of its essential co-catalytic electron donor.
2. **FMNH· cannot be generated.** The first electron removal from FMNH₂ to N3 is blocked. FMNH₂ accumulates, but it remains in the fully reduced state, lacking the semiquinone electronic configuration required for cooperative proton reduction.

##### S7.2 Normal condition (Fe-S chain open)

As calculated in S4:

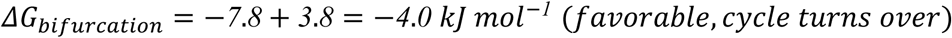

Bifurcation simultaneously generates **FMNH·** (via 1e⁻ loss to N3) and **N1a(red)** (via 1e⁻ acceptance from FMNH·). These two species then cooperate to reduce protons (S5).

##### S7.3 Rotenone-blocked condition (Fe-S chain saturated)

Rotenone blocks N2 → UQ. Electrons accumulate on the entire Fe-S chain, driving all centers toward their fully reduced states. When N3 is fully reduced, it cannot accept the high-potential electron from FMNH₂.

###### Can the high-potential electron go to N1a instead?

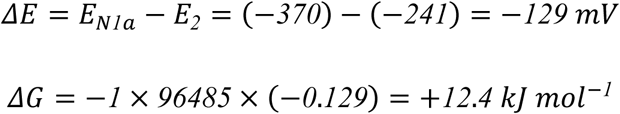

Strongly unfavorable, and N1a is a one-electron [2Fe-2S] carrier. After accepting one electron, N1a is fully reduced and unavailable. More critically, without N3 accepting the first electron, **FMNH· is never generated** — the prerequisite for the cooperative H₂-forming step is absent.

###### Can the second electron (FMNH· → FMN) go anywhere?

With both N3 (reduced by rotenone block) and N1a (would require FMNH· to reduce it) unavailable, no accessible acceptor exists within tunneling distance of FMN. The catalytic cycle cannot turn over.

###### Why doesn’t the non-bifurcating pathway (direct FMNH₂ → H₂) operate?

As shown in S2, the two-electron FMNH₂ → FMN + H₂ reaction is thermodynamically favorable (ΔG = −31.5 kJ mol⁻¹ at low P(H₂)). However, there is no evidence that CI* catalyzes direct two-electron H₂ release from FMNH₂; the proposed cooperative mechanism requires the prior one-electron oxidation to FMNH·, which generates the protonated N5 semiquinone configuration necessary for proton reduction. In the rotenone-blocked state this activation step cannot occur, and FMNH₂ retains both electrons without the semiquinone electronic configuration required for cooperative H–H bond formation.

This is analogous to the behavior of bifurcating [FeFe]-hydrogenases such as HydABC, where blocking the high-potential acceptor (ferredoxin or NAD⁺) abolishes H₂ production even though the overall reaction remains thermodynamically favorable (Furlan et al., 2022; Buckel & Thauer, 2018; Schuchmann & Müller, 2014).

##### S7.4 Summary

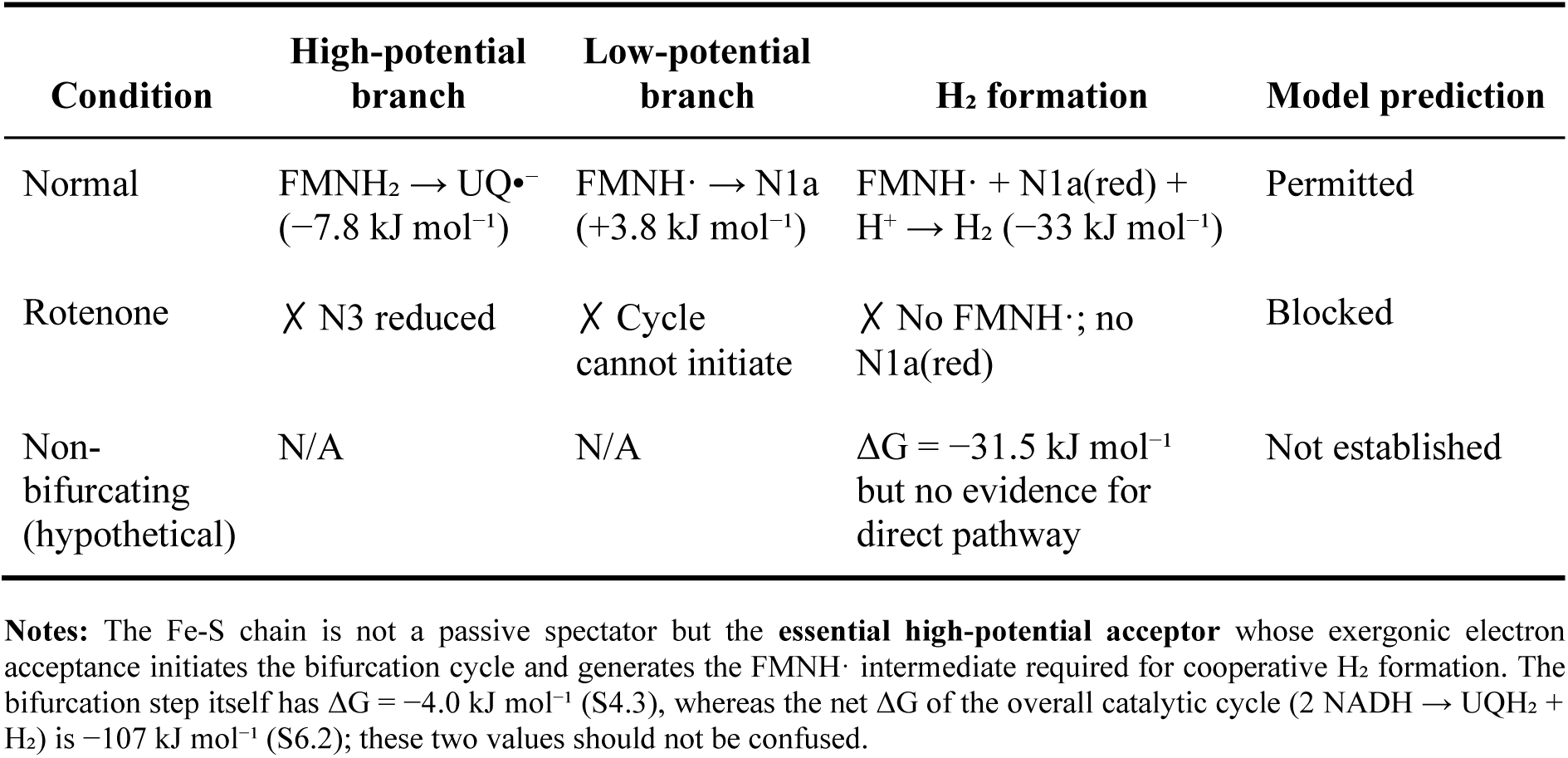

| Condition | High-potential branch | Low-potential branch | H₂ formation | Model prediction |
|---|---|---|---|---|
| Normal | FMNH₂ → UQ•⁻ (−7.8 kJ mol⁻¹) | FMNH· → N1a (+3.8 kJ mol⁻¹) | FMNH· + N1a(red) + H⁺ → H₂ (−33 kJ mol⁻¹) | Permitted |
| Rotenone | ✗ N3 reduced | ✗ Cycle cannot initiate | ✗ No FMNH·; no N1a(red) | Blocked |
| Non-bifurcating (hypothetical) | N/A | N/A | ΔG = −31.5 kJ mol⁻¹ but no evidence for direct pathway | Not established |

**Notes:** The Fe-S chain is not a passive spectator but the **essential high-potential acceptor** whose exergonic electron acceptance initiates the bifurcation cycle and generates the FMNH· intermediate required for cooperative H₂ formation. The bifurcation step itself has ΔG = −4.0 kJ mol⁻¹ (S4.3), whereas the net ΔG of the overall catalytic cycle (2 NADH → UQH₂ + H₂) is −107 kJ mol⁻¹ (S6.2); these two values should not be confused.

#### S8. Moser-Dutton Analysis: Electron Transfer Rates

The Moser-Dutton ruler estimates electron tunneling rates between cofactors:

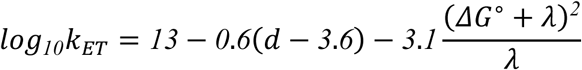

where *d* = edge-to-edge distance (Å), λ = reorganization energy (eV), ΔG° = driving force (eV). λ = 0.7 eV is used as standard value (Moser & Dutton, 1992). **Note:** We use an intercept of 13 rather than 15 for conservative estimation (see S8.6).

##### S8.1 FMN → N3 (main chain, high-potential branch)

- *d* = **6.3 Å** (edge-to-edge; Maldonado et al., 2020 — CI* structure)
- ΔG° = +0.009 eV (near equilibrium; from *E*₂ = −241 to N3 = −250 mV, pH 6; = +0.9 kJ mol⁻¹)
- λ ≈ 0.7 eV

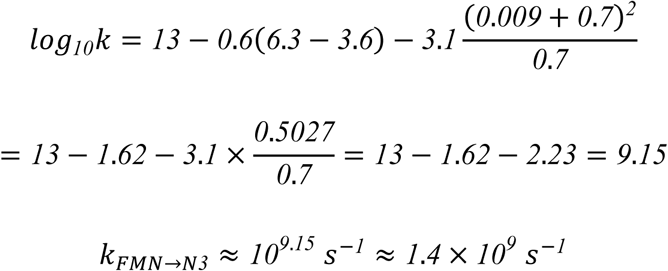

##### S8.2 FMN → N1a (low-potential branch)

- *d* ≈ **11.8 Å** (edge-to-edge; Maldonado et al., 2020 — CI* structure)
- ΔG° = +0.039 eV (endergonic; from *E*₁ = −331 to N1a = −370 mV, pH 6; = +3.8 kJ mol⁻¹, S4.2)
- λ ≈ 0.7 eV

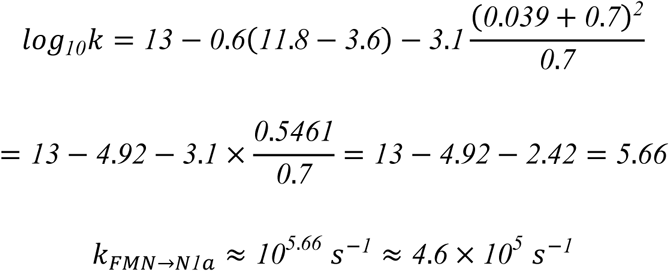

##### S8.3 N1a → FMN (off-pathway back-transfer)

In the cooperative model (S5.1), N1a(red) is consumed directly in H₂ formation together with FMNH·; the reverse N1a→FMN transfer is an off-pathway reaction (back-transfer to form FMNH₂, S5.1) that competes with, rather than constitutes, the catalytic step. Its rate therefore sets an upper bound on how much N1a(red) is lost to this competing branch. The driving force for this reverse transfer:

- *d* ≈ **11.8 Å** (same distance)
- ΔG° = −0.039 eV (exergonic reverse of FMN→N1a, pH 6)
- λ ≈ 0.7 eV

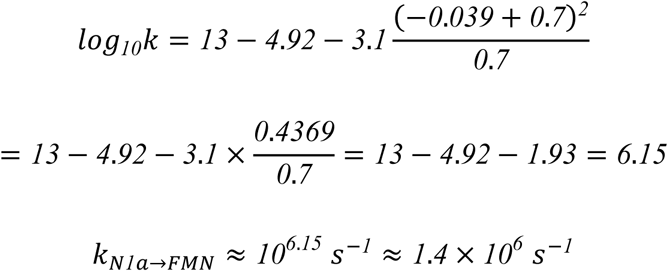

At pH 6.0 the reverse (N1a→FMN) transfer is exergonic and ∼3-fold faster than the forward (FMN→N1a) transfer. Back-transfer to form FMNH₂ is therefore kinetically accessible, so productive H₂ formation requires the N1a(red) electron to be captured by the coupled H₂-forming step before it can return to the flavin (see S8.4) — the same requirement underlying the rotenone sensitivity (S7).

##### S8.4 Interpretation

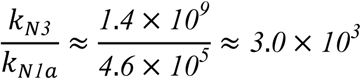

The N3 pathway remains ≈3×10³-fold faster, consistent with N3 being the kinetically preferred route for the high-potential electron. At pH 6.0 the forward FMN→N1a transfer is endergonic (+0.039 eV) and ∼3-fold slower than the reverse N1a→FMN transfer. Two consequences follow. First, even the slower forward rate (≈4.6 × 10⁵ s⁻¹) is more than two orders of magnitude faster than the overall CI turnover (≈10²–10³ s⁻¹). Thus, the estimated intrinsic FMN-to-N1a electron-transfer rate is faster than overall CI turnover, but this comparison does not establish whether net electron capture by N1a limits the proposed cycle. Second, because the reverse transfer is faster, accumulation of N1a(red) sufficient for H₂ formation requires the N1a electron to be sequestered from immediate back-transfer — plausibly by a fast confurcation (H₂-forming) step that consumes N1a(red), by conformational gating, or by a redox-balance shift when the high-potential branch is congested under micro-oxia. These remain open mechanistic questions for future kinetic studies.

The 11.8 Å FMN-to-N1a distance nevertheless yields forward and reverse rates (10⁵–10⁶ s⁻¹) well within the biologically competent range, consistent with N1a’s proposed role as the low-potential electron donor in the cooperative H₂-forming cycle.

##### S8.5 Tunneling feasibility of the FMN-to-N1a distance

The FMN-to-N1a edge-to-edge distance in CI* is 11.8 Å (Maldonado et al., 2020, Figure 5A). Although often cited as “too far for physiological electron transfer”, this distance is not beyond the range accessible to biological electron tunneling: physiological electron transfer over ∼14 Å through the protein matrix is well established, and the frequently quoted ∼14 Å reference is an empirical observation rather than a physical limit, with the protein environment (including enzyme-bound water) enhancing, not impeding, the tunneling rate (Page et al., 1999; Moser & Dutton, 1992). The FMN-to-N1a distance therefore does not exclude N1a from acting as the low-potential electron donor in the cooperative H₂-forming cycle.

##### S8.6 Interpretation limits

The Moser-Dutton calculation indicates that the selected cofactor distances do not exclude electron transfer on a biologically relevant timescale. However, the empirical model does not explicitly include detailed cofactor orientation, the actual protein tunneling pathway, conformational gating, donor/acceptor occupancy, proton-coupled electron transfer, electron back-transfer, or redox-state-dependent structural changes. The Moser–Dutton estimates suggest that the FMN–N3 and FMN–N1a separations are compatible with electron transfer on biologically relevant timescales under the assumed parameters; the distances alone do not establish the actual electron-transfer rates.

#### S9. Sensitivity Analysis

To assess the robustness of the thermodynamic conclusions, we varied key assumptions in the cooperative H₂-forming step (S5.1), whose baseline ΔG is −33.4 kJ mol⁻¹ (pH 6.0, P(H₂) = 10⁻⁶ atm, *E*₁ = −331 mV, *E*(N1a) = −370 mV).

| **Parameter** | **Baseline** | **Varied** | **ΔG_coop (kJ mol<sup>−1</sup>)** |
|---|---|---|---|
| P(H<sub>2</sub>) | 10<sup>−6</sup> atm | 10<sup>−4</sup> atm | −22.0 |
|  |  | 10<sup>−8</sup> atm | −44.8 |
| *E*(N1a) | −370 mV | −350 mV | −31.5 |
|  |  | −390 mV | −35.4 |
| *E*<sub>1</sub> (FMN/FMNH·) | −331 mV | −311 mV | −31.6 |
|  |  | −351 mV | −35.4 |

*E*₁ and *E*(N1a) enter the anode average (*E*₁ + *E*(N1a))/2 with equal weight, so their sensitivities are identical. In all cases the cooperative H₂-forming step remains exergonic, and the net catalytic cycle (S6.2, ΔG_net ≈ −106.6 kJ mol⁻¹) stays strongly exergonic (< −90 kJ mol⁻¹) under the tested variation of these parameters. Under the tested parameter variations, the qualitative conclusions — bifurcation feasibility, rotenone paradox resolution, Δp permissiveness, and cooperative H₂ formation — are unchanged.

Because the bifurcation step itself (S4.3) has the most modest driving force of the cycle (ΔG ≈ −4.0 kJ mol⁻¹), we additionally examined its sensitivity to the two assumptions that most directly affect it: the estimated one-electron potential of the first ubiquinone reduction (*E*(UQ/UQ•⁻), nominally −160 mV but ranging from −150 to −200 mV; S1) and the semiquinone potential *E*₂. The bifurcation free energy is ΔG_bifurcation = ΔG_high + ΔG_low, where ΔG_high = −F[*E*(UQ/UQ•⁻) − *E*₂] and ΔG_low = −F[*E*(N1a) − *E*₁] (S4.1–S4.3), with *E*₁ = −331 mV and *E*(N1a) = −370 mV held fixed:

| ***E*(UQ/UQ•<sup>−</sup>) (mV)** | **−150** | **−160** | **−200** |
|---|---|---|---|
| ΔG_bifurcation (kJ mol<sup>−1</sup>) | −5.0 | −4.0 | −0.2 |
| ***E*<sub>2</sub> (mV)** | **−241** | **−250 (pKa-refined)** |  |
| ΔG_bifurcation (kJ mol<sup>−1</sup>) | −4.0 | −4.9 |  |

Across this range the bifurcation remains exergonic (ΔG_bifurcation ≤ 0), although at the lowest *E*(UQ/UQ•⁻) estimate it approaches equilibrium (−0.2 kJ mol⁻¹). Even in that limiting case the complete cycle is driven by the strongly exergonic H₂-forming step (S5.1, −33.4 kJ mol⁻¹) and the overall NADH-to-H₂ reaction (S6.2, −106.6 kJ mol⁻¹), so the qualitative conclusion — a feasible, regulated low-flux bifurcation — is unchanged.

#### S10. Effect of pH on the H₂ Formation Step

##### S10.1 Thermodynamic effect of pH

The cooperative H₂-forming reaction (FMNH· + N1a(red) + H⁺ → H₂) involves one proton. Its free energy depends on pH:

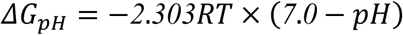

At 25°C, 2.303RT = 5.71 kJ mol⁻¹. Thus:

- A decrease of 1 pH unit makes ΔG more negative by 5.71 kJ mol⁻¹.
- An increase of 1 pH unit makes ΔG less negative by 5.71 kJ mol⁻¹.

Equivalently, for the 2H⁺/2e⁻ couple:

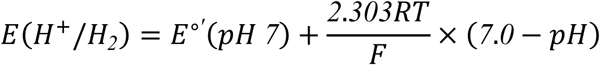

At 25°C, 2.303RT/F = 59.2 mV per pH unit.

##### S10.2 Combined effect of low P(H₂) and pH

Using P(H₂) ≈ 10⁻⁶ atm, the net free energy for the cooperative H₂ formation step at each pH:

| **pH** | **ΔG_pH (kJ mol<sup>−1</sup>)** | **ΔG_net (kJ mol<sup>−1</sup>)** | ***E*(H<sup>+</sup>/H<sub>2</sub>) (mV)** |
|---|---|---|---|
| 8.0 | +5.71 | −21.9 | −296 |
| 7.0 | 0 | −27.6 | −237 |
| 6.5 | −2.86 | −30.5 | −207 |
| 6.0 | −5.71 | −33.4 | −177 |
| 5.5 | −8.57 | −36.2 | −148 |

**Note:** *E*(H<sup>+</sup>/H<sub>2</sub>) is evaluated at P(H<sub>2</sub>) = 10<sup>−6</sup> atm (consistent with ΔG\_net). ΔG\_net = −2F·Δ*E*, where Δ*E* = *E*(H<sup>+</sup>/H<sub>2</sub>) − *E*\_avg(anode) and *E*\_avg(anode) = (*E*<sub>1</sub> + *E*(N1a))/2; *E*<sub>1</sub> is proton-coupled (1e<sup>−</sup>/1H<sup>+</sup>, +59 mV per pH unit) so *E*\_avg also shifts with pH, whereas *E*(N1a) = −370 mV is pH-independent. At pH 6.0, the H<sub>2</sub> formation step is thermodynamically favorable by ΔG\_net ≈ −33.4 kJ mol<sup>−1</sup>.

##### S10.3 Kinetic considerations

Beyond thermodynamics, lowering the assay pH from 7.0 to 6.0 increases proton activity tenfold and could facilitate proton delivery to FMN–N5. The observed maximum in H₂ accumulation at pH 6.0 is therefore consistent with combined thermodynamic and kinetic contributions, although the external assay pH does not directly report mitochondrial matrix pH.

The experimental observation that H₂ production is maximal at pH 6.0 reflects a balance between this kinetic acceleration and enzyme stability. Below pH 5.5, the matrix becomes too acidic, which may compromise protein structure and overall CI* activity.

##### S10.4 Biological implications

The pH dependence observed in the H₂ accumulation assays suggests that acidification during oxygen deprivation could favor CI*-mediated H₂ production. The mitochondrial matrix is typically alkaline under resting conditions (approximately pH 7.8 in rat liver mitochondria; Nicholls, 1974). In plants, oxygen limitation can reduce respiratory proton pumping, while the cytosol of anoxic tissues has been reported to acidify to approximately pH 6.8 (Roberts et al., 1984; Gout et al., 2001). These observations make matrix acidification plausible (Schwarzländer & Finkemeier, 2013), but cytosolic pH does not establish matrix pH or show that it reaches the pH 6.0 used in our assays.

At a fixed H₂ partial pressure, the model predicts that acidification makes the cooperative H₂-forming step more thermodynamically favorable (Section S10.2). It does not predict a kinetic optimum at pH 6.0. Matrix acidification could therefore contribute to the increase in H₂ production proposed for Phase III, but its magnitude and timing relative to H₂ accumulation remain to be established experimentally.

#### S11. Summary of Numerical Results

Unless otherwise stated, all values are calculated at 25 °C (298.15 K).

| **Calculation** | **Result** |
|---|---|
| Direct H<sub>2</sub> formation: FMNH<sub>2</sub> → FMN + H<sub>2</sub>, biochemical standard state (pH 7.0, P(H<sub>2</sub>) = 1 atm, unit flavin activities) (S2.1)<sup>a</sup> | ΔG°′ = +14.3 kJ mol<sup>−1</sup> |
| Direct H<sub>2</sub> formation, pH 6.0, P(H<sub>2</sub>) = 10<sup>−6</sup> atm, a(FMNH<sub>2</sub>)/a(FMN) = 1<sup>a</sup> | ΔG′ ≈ −20.0 kJ mol<sup>−1</sup> |
| Direct H<sub>2</sub> formation, pH 6.0, P(H<sub>2</sub>) = 10<sup>−6</sup> atm, a(FMNH<sub>2</sub>)/a(FMN) = 100<sup>a</sup> | ΔG′ ≈ −31.5 kJ mol<sup>−1</sup> |
| Cooperative H<sub>2</sub> formation (FMNH· + N1a(red)), pH 7.0, P(H<sub>2</sub>) = 1 atm | ΔG′ ≈ +6.6 kJ mol<sup>−1</sup> |
| Cooperative H<sub>2</sub> formation, pH 6.0, P(H<sub>2</sub>) = 1 atm | ΔG′ ≈ +0.9 kJ mol<sup>−1</sup> |
| Cooperative H<sub>2</sub> formation, pH 6.0, P(H<sub>2</sub>) = 10<sup>−6</sup> atm | ΔG′ ≈ −33.4 kJ mol<sup>−1</sup> |
| Net reaction: 2 NADH + 2 H<sup>+</sup> + UQ → 2 NAD<sup>+</sup> + UQH<sub>2</sub> + H<sub>2</sub>, pH 6.0, P(H<sub>2</sub>) = 10<sup>−6</sup> atm | ΔG′ ≈ −106.6 kJ mol<sup>−1</sup> |
| CI* FMN → N3 Moser-Dutton rate | 1.4 × 10<sup>9</sup> s<sup>−1</sup> |
| CI* FMN → N1a Moser-Dutton rate | 4.6 × 10<sup>5</sup> s<sup>−1</sup> |
| CI* N1a → FMN reverse rate | 1.4 × 10<sup>6</sup> s<sup>−1</sup> |

**Notes:** All values are calculated in this study (25 °C unless indicated); none are literature or measured values. Derivations: rows 1–3, S2.1–S2.5; rows 4–5, S5.1; row 6, S5.1 and S10.1–S10.2; row 7, S6.2; rows 8–10, S8.1–S8.3. Input midpoint potentials and their sources are listed in S1; the distance, reorganization energy and driving-force inputs for the rate estimates are given in S8.
<sup>a</sup> Rows 1–3 (direct H<sub>2</sub> formation): reference state and calculation. The standard transformed Gibbs energy (ΔG°′) was evaluated in S2.1 at pH 7.0 and 298.15 K, with p° = 1 atm and unit flavin activities. Using the pH 7.0 reduction potentials tabulated in S1 (vs SHE), ΔE°′ = (−0.414) − (−0.340) = −0.074 V, giving ΔG°′ = −2FΔE°′ = +14.28 kJ mol<sup>−1</sup>, reported as +14.3 kJ mol<sup>−1</sup>. Under the 2e<sup>−</sup>/2H<sup>+</sup> model adopted in S1.1, both half-cell potentials shift equally between pH 7.0 and pH 6.0, so their difference is unchanged at fixed hydrogen pressure and flavin activity ratio. Nonstandard values follow from the same potentials (S2.2–S2.4) or, equivalently, from ΔG′ = ΔG°′ + RT ln[(P(H<sub>2</sub>)/p°)(a(FMN)/a(FMNH<sub>2</sub>))], with activity coefficients assumed to cancel (S2.3); The value in row 3 follows the calculation in Section S2.4, using the rounded potential difference of +163 mV.

#### S12. Overall Conclusions

1. Under biochemical standard-state conditions at pH 7 and P(H₂) = 1 atm, the direct reaction FMNH₂ → FMN + H₂ is thermodynamically unfavorable: ΔG°′ = +14.3 kJ mol⁻¹.
2. At P(H₂) = 10⁻⁶ atm, the same reaction is favorable even when FMNH₂/FMN = 1: ΔG ≈ −20.0 kJ mol⁻¹. Under the 2e⁻/2H⁺ model used in S1.1, the two half-cell potentials shift equally between pH 7.0 and pH 6.0, so the calculated ΔG′ is unchanged at fixed hydrogen pressure and flavin activity ratio.
3. The **cooperative H₂ formation model** (FMNH· + N1a(red) + H⁺ → FMN + N1a(ox) + H₂) is thermodynamically favorable under experimental conditions: ΔG ≈ −33.4 kJ mol⁻¹ at pH 6.0, P(H₂) = 10⁻⁶ atm. This model rigorously conserves electrons (2e⁻ from two distinct one-electron donors) and protons.
4. The proposed UQ-coupled net catalytic cycle (2 NADH + 2H⁺ + UQ → 2 NAD⁺ + UQH₂ + H₂) is strongly exergonic: ΔG ≈ −106.6 kJ mol⁻¹ under the assumed experimental conditions.
5. we propose that FCCP-induced lowering of the FMNH₂/FMN ratio cannot, by itself, make the H₂-forming step thermodynamically unfavorable at very low P(H₂); rather, FCCP inhibits H₂ production by depleting the reduced intermediates (FMNH·, N1a(red)) required for the cooperative catalytic cycle (S2.5).
6. The proposed model offers a possible explanation for rotenone inhibition: blockade of the high-potential branch would prevent both FMNH· generation and N1a(red) formation, thereby suppressing the cooperative H₂-forming step even though direct FMNH₂ → H₂ remains thermodynamically favorable.
7. In the cooperative model (Model A), N1a serves as a co-catalytic electron donor, providing the second electron for H–H bond formation together with the N5-bound hydrogen of FMNH·; alternatively, N1a itself could act as the catalytic site through a metal-hydride (Fe–H⁻) pathway (Model B). We currently favor Model A on structural and mechanistic grounds but do not formally exclude Model B.
8. The Moser-Dutton rate estimates at pH 6.0 (intercept = 13) indicate that FMN→N3 (≈1.4 × 10⁹ s⁻¹), FMN→N1a (≈4.6 × 10⁵ s⁻¹), and N1a→FMN (≈1.4 × 10⁶ s⁻¹) are estimated to occur on timescales compatible with the proposed low-flux catalytic cycle.
9. Establishing this mechanism definitively would require independent evidence including: spectroscopic detection of FMNH· and N1a(red) under H₂-producing conditions; isotope-labeling experiments using H/D tracers; mutations of N1a ligands to test its essential role in H₂ formation; and direct testing of whether the overall reaction exhibits the 2 NADH : 1 H₂ : 1 UQH₂ stoichiometry predicted by the proposed UQ-coupled cycle.

#### S13. Discriminating Experiments

The following experiments test the proposed two-turnover cycle and, separately, seek to distinguish the two H₂-forming-site models (S5.1 vs S5.3). The highest-priority measurement is the overall stoichiometry: simultaneous quantification of NADH consumption, H₂ formation, and UQH₂ production. A reproducible 2 NADH : 1 H₂ : 1 UQH₂ ratio would be consistent with the two-turnover cycle (S6.1), but would not by itself establish electron bifurcation or distinguish the two catalytic-site models. Conversely, an approximately 1 NADH : 1 H₂ ratio with negligible UQH₂ production would favor a pathway that does not require coupled UQ reduction, such as direct two-electron H₂ formation from FMNH₂ (S2.4).

1. **N1a ligand mutagenesis.** Mutation of the N1a cysteine ligands is predicted to abolish H₂ production if the N1a iron is the direct catalytic site (Model B), whereas it would primarily impair electron delivery if FMN-N5 is the site (Model A). However, loss of the N1a cluster may also destabilize the adjacent NDUFV2 subunit or alter the FMN microenvironment, so this experiment alone cannot unambiguously distinguish the models. The most informative mutations are those that retain N1a assembly, global complex integrity, and canonical NADH oxidation while selectively altering H₂ formation (a functional-separation phenotype).
2. **Flavin semiquinone EPR.** EPR spectroscopy under H₂-producing conditions should detect a flavin semiquinone radical if Model A is operative; its absence, together with Fe–H⁻ signatures, would favor Model B.
3. **Proton-channel mutagenesis.** Mutation of residues controlling proton access to FMN–N5 is predicted to impair H₂ production in Model A by limiting proton delivery to the proposed catalytic site, although such an effect would not by itself distinguish the two models.
4. **Isotopic labeling.** D₂O or NADH-d₂ labeling can identify the hydrogen source and distinguish a flavin-hydride (N5) from an Fe–H⁻ pathway.
5. **X-ray absorption spectroscopy (XAS).** Fe K-edge XAS under turnover could reveal changes in the oxidation state or coordination environment of the N1a iron relevant to Model B
6. **Direct bifurcation detection.** Time-resolved EPR or ultrafast spectroscopy could directly follow the partitioning of electrons from photo-reduced FMNH₂ into N3 and N1a, providing the strongest test of bifurcation; if CI* is stabilized in a state where N1a accumulates in the reduced form in the absence of H₂ production, this would support bifurcation. This remains the key unresolved mechanistic step.

